# Bayesian Estimation of Allele-Specific Expression in the Presence of Phasing Uncertainty

**DOI:** 10.1101/2024.08.09.607371

**Authors:** Xue Zou, Zachary W. Gomez, Timothy E. Reddy, Andrew S. Allen, William H. Majoros

**Author notes:** To whom correspondence should be addressed. &. These authors contributed equally.

## Abstract

**Motivation:** Allele-specific expression (ASE) analyses aim to detect imbalanced expression of maternal versus paternal copies of an autosomal gene. Such allelic imbalance can result from a variety of cis-acting causes, including disruptive mutations within one copy of a gene that impact the stability of transcripts, as well as regulatory variants outside the gene that impact transcription initiation. Current methods for ASE estimation suffer from a number of shortcomings, such as relying on only one variant within a gene, assuming perfect phasing information across multiple variants within a gene, or failing to account for alignment biases and possible genotyping errors.

**Results:** We developed BEASTIE, a Bayesian hierarchical model designed for precise ASE quantification at the gene level, based on given genotypes and RNA-Seq data. BEASTIE addresses the complexities of allelic mapping bias, genotyping error, and phasing errors by incorporating empirical phasing error rates derived from Genome-in-a-Bottle individual NA12878. BEASTIE surpasses existing methods in accuracy, especially in scenarios with high phasing errors. This improvement is critical for identifying rare genetic variants often obscured by such errors. Through rigorous validation on simulated data and application to real data from the 1000 Genomes Project, we establish the robustness of BEASTIE. These findings underscore the value of BEASTIE in revealing patterns of ASE across gene sets and pathways.

**Availability and Implementation:** The software is freely available from https://github.com/x811zou/BEASTIE. BEASTIE is available as Python source code and as a Docker image.

**Supplementary information:** Additional information is available online.

## Introduction

Noncoding genetic variation has a major role in human traits and disease. For common and complex traits and diseases, genetic association studies typically identify non-coding genetic variants. Those associated variants have often been linked to impacts on gene regulatory activity [1] ; and the narrow sense genetic heritability of such common traits and diseases is also strongly enriched in gene regulatory regions [2]. Non-coding genetic variation also plays a major role in rare diseases that collectively impact an estimated 3-4% of the global population [3, 4]. For example, the diagnostic yield of rare diseases via DNA sequencing is 50% [5]. Supplementing DNA sequencing with gene expression measurements and RNA sequencing provides additional information about the potential disruption of a gene in a patient. In some cases, doing so has increased diagnostic rates by upwards of 15% through analysis of altered gene splicing, allele-specific expression, and unusual gene expression levels [6]. Together, these findings suggest that including gene expression measurements in studying traits and diseases can substantially improve our understanding of disease mechanisms.

Allele-specific expression (ASE), defined as an allelic imbalance in transcript quantity between maternal and paternal copies of autosomal genes, is an important indicator of a possible gene expression outlier. Altered gene expression can arise via many mechanisms including genetic variation in promoters and enhancers and loss-of-function mutations that impact mRNA splicing or stability and result in transcript degradation. Predicting such effects from DNA sequencing alone remains an unmet challenge. Compared to differential gene expression analysis, ASE analysis also has the advantage that it compares the expression of copies of a gene within the same nucleus, potentially increasing the power to detect expression anomalies. Recent advancements have illustrated the utility of incorporating ASE analysis in rare disease diagnostics, for example, elevating the diagnostic rate by 7.5% across various disease types [4, 7, 8]. Further, ASE analysis stands as a pivotal tool for identifying the existence of potential cis-regulatory effects in gene expression in both human and non-human organisms [4, 7, 9, 10, 11, 12].

The estimation of ASE can be strongly confounded by a number of issues, including (1) the availability of exonic heterozygous sites in a gene; (2) the need to combine read counts across sites in a phase-consistent manner; (3) the possibility of genotyping errors; and (4) allelic mapping bias. We discuss these in detail below.

### Availability of heterozygous sites

Among 445 individuals from the 1000 Genome Project, the average number of heterozygous (het) sites per gene is 3.3. Roughly 30% of genes have one exonic het SNP, while 34% have 4 or more exonic het SNPs. While some methods make use of only one variant in a gene, others attempt to make use of all sites by aggregating read counts across sites [13, 14]. When using short-read sequencing, individual reads may span only a subset of those variants, which presents the problem of how to combine read counts across sites within a gene, as using the counts at only one site to estimate ASE fails to make use of all available information. This is the phasing problem: determining which alleles at multiple het sites originate from the same chromosome.

### Combining read counts across sites in a phase-consistent manner

For short reads spanning individual het sites, allelic read counts need to be combined across sites in a manner that observes phasing, so that reads are summed across the maternal and paternal chromosomes separately. Unfortunately, the relative phasing of sites is often not known with certainty. One class of methods heuristically addresses this problem by utilizing pseudo-phasing [13, 15], which assumes that the alleles with the higher read count at each site always originate from the same gene copy. While there should be a statistical tendency for the alleles with higher counts to often come from the same gene copy in the presence of ASE, the assumption that they always do so is not generally correct, particularly under the null hypothesis (no ASE), and thus pseudo-phasing can result in overestimation and elevated Type I error. A more principled approach is to either predict a single best-supported phasing and use that as a proxy for the true phasing or to explicitly account for phasing uncertainty statistically when estimating ASE. Using a single predicted phasing (e.g., from methods such as phASER [16] or SHAPEIT2 [17]) has the shortcoming that the ASE estimate will be impacted by any errors in the predicted phasing. As we describe shortly, our approach instead explicitly accounts for phasing error rates when estimating the full posterior distribution of ASE, while marginalizing out the true phasing in a statistically rigorous way.

### Genotyping error

Genotyping errors occur when variant calling or genotyping algorithms mislabel genotypes, leading to a zero read count on an allele in RNA [18] and false signals of allelic imbalance. Various approaches have been proposed to remove sites with incorrect genotypes, including the use of customized genotype error filters that consider sequencing noise and coverage [19], developing a customized genotype caller to compute genotype at each SNP position [20], deploying a new genotype caller that only requires RNA-Seq data [21], jointly inferring genotypes in a ASE model [22], and filtering nonsense variants if the alternative allele ratio was low [23]. Some researchers have instead relied on having higher-quality source data [9, 14, 19, 24, 25, 26, 27], such as HapMap, 1000 Genomes, and GIAB, without additional base-calling correction, thereby constraining their analyses to data with established low base-calling errors.

### Allelic mapping bias

Allelic mapping bias arises when reads that contain the reference allele at a heterozygous site are successfully aligned to the reference genome at a higher rate than reads bearing the alternate allele, leading to skewed expression estimates that do not accurately reflect the true allelic expression ratios [23]. To address this challenge, prior studies have adopted various methods such as using variant-aware aligners [19, 28, 29], implementing adjusted binomial models to take into account the pre-existing allelic bias [13, 20], and filtering out sites through customized mappability filters or simulations [9, 14, 19, 22, 27, 30].

Here we introduce BEASTIE, a novel approach for estimating ASE that accounts for all of the foregoing challenges, in a statistically principled way. By leveraging a probabilistic Bayesian framework, this model allows for the incorporation of prior information to yield more precise posterior estimates of allelic imbalance. BEASTIE makes use of an external phasing algorithm but accounts for possible phasing errors in a locus-specific and variant-specific manner by studying local phasing error rates and using those to statistically marginalize over all possible phasings when estimating ASE. Doing so enables BEASTIE to produce more robust estimates than methods that either use only one site or assume a single predicted phasing without accounting for possible phasing errors. We also provide a processing pipeline that mitigates alignment bias by simulation and streamlines the processing of raw RNA-Seq fastq reads and VCF data to compute and output gene-level ASE, establishing BEASTIE as a comprehensive tool for the investigation of allelic imbalance.

## Material and methods

In order to integrate information across multiple heterozygous sites within a gene while accounting for phasing uncertainty, we designed a Bayesian probabilistic graphical model in which both ASE and phasing are treated as latent variables. Because these entities are both latent, we can do joint inference on them, thus allowing our estimate of ASE to take into account our uncertainty about phase. The resulting model, BEASTIE, is depicted in **Fig. 1**. Variable definitions are given in **Table 1**.

**Table 1.**
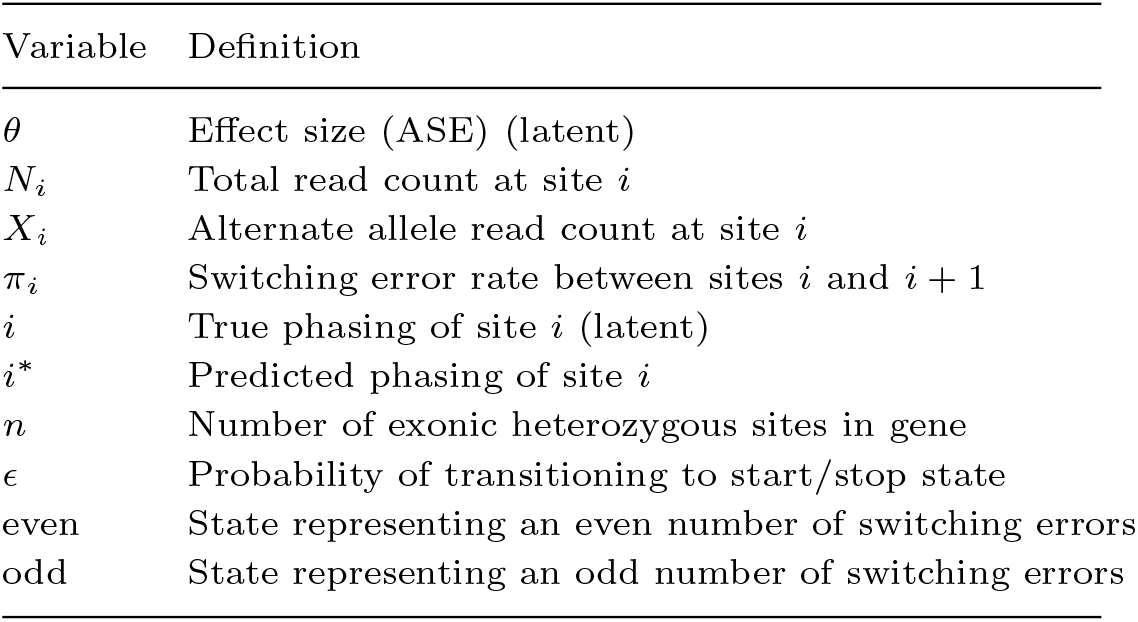
Definitions of the variables in Fig. 1.

| Variable | Definition |
| --- | --- |
| $\theta$ | Effect size (ASE) (latent) |
| $N_i$ | Total read count at site $i$ |
| $X_i$ | Alternate allele read count at site $i$ |
| $\pi_i$ | Switching error rate between sites $i$ and $i + 1$ |
| $i$ | True phasing of site $i$ (latent) |
| $i^*$ | Predicted phasing of site $i$ |
| $n$ | Number of exonic heterozygous sites in gene |
| $\epsilon$ | Probability of transitioning to start/stop state |
| even | State representing an even number of switching errors |
| odd | State representing an odd number of switching errors |

**Fig. 1.**
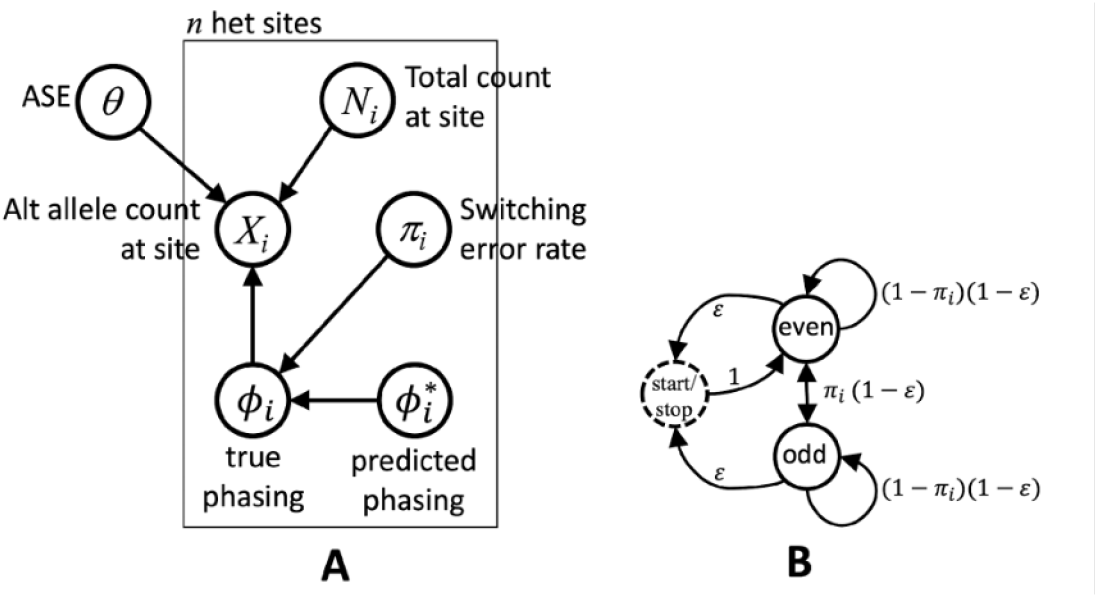
(A) The BEASTIE model for estimating ASE from multiple exonic heterozygous sites in a gene while accounting for phasing uncertainty. (B) An inhomogeneous hidden Markov model for summing in linear time over all possible phasings based on site-specific phasing error rates.

BEASTIE is intended to be applied to each gene independently to estimate the degree of ASE, expressed as an odds, *θ*:

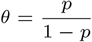

Where *p* is the relative proportion of expression of the gene originating from the maternal copy. Note that we arbitrarily label one chromosome as maternal, for simplicity, given that the two chromosomes are exchangeable under our statistical model.

Inputs to the model consist of *n*, the number of exonic heterozygous sites in the gene; *N*_*i*_, the total number of reads spanning site *i*; *X*_*i*_, the number of reads spanning site *i* that contain the alternate allele; *π*_*i*_, the expected per-site switching error rate (for phasing) between sites *i* and *i* + 1; and 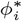, the predicted phasing of site *i*. Read counts *N*_*i*_ and *X*_*i*_ can be obtained by simple counting of reads aligned to the gene that spans each site; to avoid double-counting individual reads that span multiple sites, we provide a script to thin the list of heterozygous sites so that no two sites are within a read-length of each other in transcript distance.

The expected switching error rates *π*_*i*_ are provided by a separate model, SELR (Switching Error Logistic Regressor), detailed in **Supplementary Text 2.3**, which predicts site-specific switching error rates based on linkage disequilibrium, allele frequencies, and distances between sites. We provide a pre-trained version of the model tuned specifically for the statistical phaser SHAPEIT2 [17] ; a script is also provided for re-training the SELR model for arbitrary phasing programs. We also provide a version of BEASTIE that does not require site-specific error rate estimates. That version, called BEASTIE-latent, treats the phasing error rate as a latent variable, which is automatically marginalized out of the model during inference.

Due to the exchangeability of maternal versus paternal chromosome labels in our model, and with the relative nature of statistical phasing (i.e., statistical phasing of a given site is generally only meaningful in relation to the phasing of other sites), we model relative rather than absolute phasing. Thus it is sufficient to fix one site and phase all others relative to that site. To promote identifiability of the model, we choose the site with the highest coverage as the fixed site (details can be found in **Supplementary Fig. S15**); that is, we assume 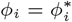 where *i* is the highest coverage site.

Phasing at this site serves as a reference, with the model assessing phasing for other sites in the gene relative to this one. This approach integrates phasing information into allele frequency estimation, accounting for uncertainties inherent in phasing data.

Given this arbitrary phasing of the site with the highest coverage, the phasing of all other sites relative to that site determines how the binomial likelihoods are evaluated. In particular, the likelihood of *X*_*i*_ is a binomial in which the probability parameter is either p (when the alternate allele is on the maternal chromosome) or 1 − *p* (when the alternate allele is on the paternal chromosome). Our BEASTIE model has two implementations. For the BEASTIE STAN version missing phasing error rates are treated as latent variables, as described in **Supplementary Text Section 2.1**. In the BEASTIE C++ version, phasing errors are incorporated as parameters determined by SELR or set as default parameters (5%), as outlined in **Supplementary Text Section 2.2**.

## Results

### Phasing ambiguity is an important problem in estimating ASE

We conducted an analysis of the number of exonic heterozygous (het) sites within each gene across 445 individuals from the 1000 Genomes Project and observed a long-tailed distribution (**Supplementary Fig. S1**), with 34% of genes containing four or more exonic het sites, and an average of 3.3 exonic het sites per gene. This distribution highlights the need for analytical methods that aggregate information across multiple genetic loci. The limitations of short-read sequencing technologies, which often capture only fragments of the genetic landscape, necessitate accurate assembly of these fragments.

Accurate aggregation of read counts across sites requires resolving their phase. Even in trio data, in which we can use parental genetic information to phase offspring’s het sites, triple heterozygous sites (triple-hets) remain unphased. In one trio from the 1000 Genomes Project (individuals NA19240, NA19238, NA19239; **Supplementary Text 1.4**), approximately 20% of sites were unphaseable due to triple-het configurations. This was consistent with simulations using trio transmission data derived from parental samples of diverse ancestries (**Supplementary Table S2**).

We evaluated the phasing accuracy of SHAPEIT2, a widely used statistical phaser, in three individuals. In NA19240, phased using trio phasing, we observed a 2.2% switching error rate for phasable child heterozygous sites. In addition, our analysis of individuals from two ancestries documented genome-wide switching error rates, finding an average rate of 4.13% for a Utah female (NA12878) against the GIAB gold standard reference phasing and 4.03% for a Yoruba male (GM19440), using experimentally determined haplotype phasing (**Supplementary Table S4, Supplementary Text 1.2 & 1.3**).

Despite generally low phasing errors observed, our study identified instances of high phasing errors, particularly in genes with larger numbers of exonic het sites,(**Supplementary Table S5**; Spearman *ρ* = 0.6854692, *p* = 0.002388332), SNP pairs separated by long distances, and SNPs with low minor allele frequencies (MAFs), as detailed below. We found a strong and statistically significant correlation between the number of exonic heterozygous sites within a gene and the likelihood of phasing errors. Our logistic regression analysis (error ∼ num.of.hets) revealed that as the number of heterozygous sites increases, the probability of encountering phasing errors also increases significantly (**Supplementary Fig. S2**). This relationship was confirmed by the ANOVA (code detailed in **Supplementary Text 4.3**), where the inclusion of the number of heterozygous sites as a predictor resulted in a substantial reduction in residual deviance (from 1734.2 to 1718.1) and an extremely low p-value (5.897e-05), indicating a highly significant effect. Specifically, we noted a percentage of genes with switching error of 7.4% for genes with two heterozygous sites, increasing to 8.5% for three sites, and peaking at 35.2% for genes with thirteen heterozygous sites (**Supplementary Table S5**). Examination of the MAF groups for each SNP pair demonstrated a contrast in average phasing error rates based on allele rarity (**Supplementary Fig. S3B**), with rare SNPs (MAF *<* 0.2%) exhibiting average phasing error rates as high as 48.55%, whereas common SNPs (MAF *>* 5%) had significantly lower average phasing error rates, at 3.12% (**Supplementary Table S6**). Furthermore, phasing error rates increased significantly from 4% to 8.37% when inter-SNP distances exceeded 1 kilobase, emphasizing the challenges in accurately phasing variants over larger genomic distances.

### BEASTIE performance

#### BEASTIE outperforms competing methods on realistic simulation data

We evaluated the performance of BEASTIE in comparison to two established methods: MajorSite (MS) and NaiveSum (NS), which build upon the methodologies reported in earlier studies [16, 19, 31]. MajorSite makes use of only one site within each gene – the site with the highest count. When multiple sites are present in a gene, MajorSite fails to make use of all available information. On the other hand, NaiveSum (NS) aggregates allelic read counts across exonic heterozygous sites within a gene, under the assumption of perfect phasing by SHAPEIT2 and drawing on concepts from previous studies [20]. However, the accuracy of this method is compromised by the existence of phasing errors, resulting in the summation of incorrect counts. A third approach is pseudo phasing, which assumes that the allele with the higher count at each site originates from the same haplotype, an approach introduced by MBASED [13]. This method is excluded from our comparison due to its inability to adequately control type 1 error rate.

This comparative analysis focused on each method’s proficiency in distinguishing positive cases (nonzero ASE) from negative cases (zero ASE) within simulated datasets having 10 het sites per gene. We simulated read depths ranging from 5 to 20 reads per variant and effect sizes of *θ* = 0.5 or 0.75. The discriminative efficacy was quantified via AUC values (details in **Supplementary Text 1.6**), particularly in distinguishing genes with *θ* = 0.5 or 0.75 from genes conforming to the null hypothesis (*θ* = 1).

Empirical findings demonstrated NaiveSum’s superiority over MajorSite at 5% or 10% switching error rates, demonstrating the utility of combining information across multiple sites. In scenarios devoid of switching errors, NaiveSum’s performance was on par with BEASTIE, while MajorSite was disadvantaged in multi-site contexts. Notably, BEASTIE outperformed NaiveSum in datasets exhibiting non-zero switching error rates (**Supplementary Fig. S5A**).

For genes with 10 heterozygous sites, the accuracy of BEASTIE with default 5% phasing error was predominantly influenced by the sequencing depth per site and the effect size. An increase in read depth from 5 to 20 reads per site substantially enhanced BEASTIE’s accuracy (AUC from 0.898 to 0.998), especially when the switching error rate was 5% and the effect size was 0.5. Elevating the switching error rate to 10% resulted in a diminished accuracy at lower read counts; however, at a simulated 20 reads per site, the accuracy of BEASTIE remained stable (AUC between 0.994 and 0.998) for data with switching error rates ranging from 0-10% (**Supplementary Fig. S5B**). Furthermore, a decrease in the effect size from *θ* = 0.5 to *θ* = 0.75 significantly reduced BEASTIE’s accuracy (AUC from 0.827 to 0.747) in datasets with 20 reads per site and switching error rates between 0-10%. While a depth of 5 reads per site was insufficient for reliable detection of subtler effects, enhancing the depth to 20 reads per site significantly improved accuracy (AUC at 0.747 for *θ* = 0.75), even in scenarios characterized by weaker effects and a 10% switching error rate (**Supplementary Fig. S5C**).

#### Phasing error rates are accurately predicted by genetic and genomic features

To enhance the accuracy of phasing error estimates within the BEASTIE framework, we developed the Switching Error Logistic Regressor (SELR), a logistic regression model designed to predict the switching error rate of SNP pairs processed by SHAPEIT2. This model was calibrated on data from the NA12878 sample, and SHAPEIT2 phasing errors were identified by comparing each SNP pair’s phasing against the gold standard phasing (from GIAB), with discrepancies marked as errors.

SELR utilizes a number of predictive features, including the minimum minor allele frequency (MAF) within the SNP pair, the differential MAF between SNP pairs, the log10-transformed inter-SNP distance, and the linkage disequilibrium (LD) metrics (*r*^2^ and *D*′), as well as second-order interaction terms between these variables (**Supplementary Table S21**). GIAB data was randomly split into two halves: the model was trained on 8,729 site pairs and validated against a disjoint set of 8,730 site pairs, each set comprising approximately 3.7% incorrectly phased site pairs. SELR attained an impressive AUC of 0.8425 on validation data set (**Supplementary Fig. S7B**), indicating a high level of discrimination between correctly and incorrectly phased site pairs. These features were selected based on their observed correlations with switching error rates, and the training/testing details as described in **Supplementary text 2.3**.

Our analysis on NA12878 data identified several trends: average SHAPEIT2 phasing error rates increase with greater inter-SNP distances and decrease in scenarios of high MAF and strong LD (*r*^2^, *D*′) (**Supplementary Fig. S3**). Similar trends were observed in the predicted switching error rate per SNP pair from SELR (**Supplementary Fig. S4**). These findings align with the expectation that for statistical phasing methods such as SHAPEIT2, rarer SNPs and those with greater distances between them are more prone to switching errors, as are SNP pairs characterized by low LD. Additionally, we observed a consistent trend across all chromosomes where higher MAF values correlate with reduced switching errors, while rare SNPs, particularly those with less than 0.2% MAF, are associated with increased switching errors (**Supplementary Fig. S8, S9, S10 and Tables S6, S7**). This pattern extends to LD values, where SNP pairs with lower LD are more likely to exhibit higher switching errors.

Upon integrating this model-predicted switching error rate into the BEASTIE framework, supplanting the fixed 5% phasing error parameter, we observed a notable performance enhancement. This optimized approach outperformed the BEASTIE model with its default 5% phasing error parameter, particularly evident in empirical simulation data derived from NA12878 (**Supplementary Fig. S6**).

### ASE traits on 1000 Genome dataset

In this study, BEASTIE was applied to a dataset of 445 samples from the 1000 Genomes Project (1KGP), representing five distinct ancestries. This dataset encompassed high-throughput RNA-Seq data from lymphoblastoid cell lines for each individual, coupled with complete 1000 Genomes phase3 genotype data. Our primary aim was to investigate the ASE landscape across different ancestries and examine ASE signals from gene sets of interest. Notably, prior ASE studies employing 1000 Genome data have assumed perfect phasing and genotyping from the 1000 Genome phase 1 dataset and Omni 2.5M SNP array data [32, 33]. In contrast, our analyses are based on our BEASTIE method that accommodates the potential for phasing errors, and are based on 1000 Genomes phase 3 data (details can be found in **Supplementary Text 1.1**).

#### BEASTIE identifies the most ASE genes

In our study analyzing this large cohort, we found that, per individual, an average of 7.23% (range: 3.14% to 19%; **Supplementary Table S10**) of genes exhibit ASE. This was observed in genes possessing at least one het site and having on average at least 10 reads per site. In comparison, when employing Naive Sum and Major Site, the proportions of genes exhibiting ASE were identified to be 6.39% and 4.37%, respectively.

#### ASE magnitude and enrichment between genesets

Our study analyzed ASE across twelve gene sets, including imprinted genes, genes tolerant to loss-of-function (LoF), genes associated with recessive conditions as per the Residual Variation Intolerance Score (RVIS), housekeeping genes, and several others related to haploinsufficiency and essential functions, as delineated in **Supplementary Table S12**. We assessed ASE rates by calculating the ratio of ASE-positive genes within each gene set for individual samples. A comparison of ASE rates between specialized gene sets and a defined “non-specialized” set of genes (not belonging to any of the gene sets considered here) revealed significant differences (Wilcoxon sign-rank test *p <* 0.05; **Supplementary Table S13**). Further analysis focused on ASE magnitudes (|log_2_(*θ*)|), revealing skewed distributions within each gene set, indicating diverse magnitudes of ASE in genes identified as having expression imbalance.

Known imprinted genes exhibited a notably high mean ASE magnitude of 3.49, reflecting strong epigenetic control, in contrast to housekeeping genes, which had a lower mean of 0.526, consistent with their role in maintaining cellular function. Genes with ASE showed a narrow mean ASE magnitude range (1.02-1.18) across gene sets (LoF tolerant, recessive, OMIM haploinsufficient, autosomal dominance) with all genes average to be 1.30 and non-specialized geneset to be 1.31 (**Supplementary Table S14**). Permutation tests comparing ASE magnitudes on ASE genes across gene sets confirmed these observations without relying on distributional assumptions, using over 10,000 randomizations for robustness. Interestingly, genes without ASE showed a narrow mean ASE magnitude range (0.213 to 0.265) across 12 gene sets (**Supplementary Table S15, Supplementary Fig. S14**).

The differentiation in ASE rates and magnitudes between gene sets, particularly between imprinted and stillbirth genes, illustrates the complexity of patterns in gene expression imbalance in our dataset. Specifically, the housekeeping gene set, with a lower ASE rate of 21.6% (versus imprinted genes of 63.8%), emphasizes their stable expression critical for cellular homeostasis. The mean ASE rate of all genes is 6.89. Non-specialized geneset (7.00), LoF geneset (13.4), recessive geneset (5.98), ClinGen haploinsufficient geneset (3.97), OMIM haploinsufficient geneset (5.29), All haploinsufficient (4.02), MGI essential geneset (6.21), and autosomal dominance geneset (6.47), stillbirth (1.64). The significant differences in ASE rates between gene sets such as stillbirth genes, ClinGen haploinsufficiency, MGI essential genes, and the non-specialized set further highlight the complex interplay of developmental and disease processes (**Supplementary Tables S13**).

#### Genes with extreme ASE magnitude

We employed a gene selection strategy to identify both known and potential novel candidates for imprinted genes in LCL, successfully highlighting the top 25 genes that exhibit extreme ASE, as detailed in **Supplementary Table S16, Supplementary Fig. S23**. Our selection criteria focus on genes with a high prevalence of extreme ASE and ASE magnitudes, consistently observed across individuals from diverse ancestries, which supports the reliability of their expression profiles. This set includes 6 previously documented genes in specific tissues: NAPIL5, HLA-DQA2, FAM50B, LPAR6, UTS2, and ZDBF2 (**Supplementary Table S17**), along with two well-known imprinted genes in PBL, PEG10 and SNRPN [34]. The remaining 17 genes (**Supplementary Table S18**), which include 3 pseudogenes, 2 RNA genes, and 6 immune gene segments, lack prior literature on their imprinting status, necessitating further validation to confirm their extreme ASE profiles. High allelic imbalance on immunoglobulin (Ig) gene loci can be caused by misalignment issues within the V(D)J recombination regions [35]. There is no expectation that the allele (reference versus alternate) would correlate with ASE status in imprinted genes, and indeed, no such association was observed (Fisher’s exact test I, all *p >* 0.1; **Supplementary Table S19**). Further analysis using the Variant Effect Predictor (VEP) [36] identified the primary SNP variants associated with ASE status in the genes IGHV3-9, WASH4P (Fisher’s exact test II, all *p <* 0.05; **Supplementary Table S19**), which include missense variants with a high PolyPhen score and low MAF score of 0.13; variants located in NMD transcripts with MAFs of 0.16, respectively. These findings are consistent with the ASE status of the genes identified by BEASTIE.

## Discussion

ASE analyses complement eQTL analyses by focusing on expression imbalance without assuming a specific mechanism. The internal control provided by comparing allelic counts within a single sample may offer stronger power from smaller sample sizes than eQTL analyses in certain cases. Furthermore, ASE analyses do not assume any particular mechanism for differences in allelic expression, making them flexible tools for exploring gene regulation.

Here we present BEASTIE, a novel Bayesian approach for detecting ASE that rigorously accounts for phasing uncertainty by treating phasing as a latent variable that is integrated out during inference. Previous solutions to estimating ASE rely on a single heterozygous site, a single predicted phasing, or pseudo phasing to combine read counts across sites. As we have shown, those strategies may not control false positive rates, are subject to incorrect phasing prediction, and ultimately miss opportunities to use all available phasing information. BEASTIE builds upon those previous studies by integrating information across exonic sites and incorporating additional information such as population allele frequencies, the distance between SNPs, and linkage disequilibrium. Through the BEASTIE modeling approach, we address several confounding challenges when estimating ASE. Specifically, we allow users to better assess the strength of evidence for allelic imbalance by reporting the full posterior distribution of ASE; we more precisely estimate phasing error rates; and we correct for systematic biases due to alignment errors and genotyping errors. Those features are particularly valuable for identifying issues such as multi-modality or highly dispersed posterior mass, which can inform the interpretation of ASE estimates.

We view this study as particularly impactful in two key applications of ASE analysis. First, there is often little phasing certainty for rare genetic variants, especially when they are more than 1 kb from a common variant. By integrating over that uncertainty, we can improve ASE inference involving such rare variants. That can potentially improve the use of RNA assays in diagnosing rare diseases and in understanding the effects of emerging variations in the human population. Second, there is also often more phasing uncertainty for genomes from populations and ancestries without established reference panels. Again, by integrating that potential uncertainty, we can improve ASE studies for more diverse populations. Both of these advances are important steps toward making genetics work for everyone.

BEASTIE’s ability to detect both known imprinted genes and genes with extreme ASE magnitudes underscores its utility in expanding our understanding of gene expression regulation. Specifically, we identified 25 additional genes with consistently extreme ASE across many individuals, including 2 known imprinting genes and 6 with imprinting status on other tissues mentioned in prior publications. Among those, all exhibited no significant association between haplotype and gene expression. Six of those genes are immunoglobulins. For those, the extreme ASE is likely due to a combination of V(D)J recombination, jackpotting in lymphoblastoid cell cultures, and alignment artifacts. The remaining 11 genes may be novel candidates for imprinting in the human genome, or may instead be subject to random inactivation or genetic alteration via other mechanisms. Our findings also reveal significant differences in ASE rates across various gene sets, reflecting the complex interplay between genetic regulation and functional categories. The observed higher ASE magnitudes in imprinted genes, compared to housekeeping genes, are consistent with strong epigenetic regulation in imprinted loci. In contrast, the narrow range of ASE across other gene sets indicates a degree of stability in expression imbalance that may be reflective of underlying genetic or regulatory constraints. The significant depletion of ASE in the stillbirth gene set in particular suggests stringent selective pressure to maintain stable expression in genes crucial for cellular homeostasis and organismal viability. These results align with the expected selection against disruption in essential genes, further illustrating the nuanced regulatory landscape that BEASTIE can help to elucidate.

The flexibility of our Bayesian modeling framework offers numerous opportunities for future improvements. Given the widespread use of short-read sequencing and the high costs of long-read sequencing, our approach remains relevant. There is a substantial amount of legacy short-read data, and long-read data is often too expensive for extensive quantitative use. Combining legacy short-read data with Bayesian updating methods is thus advantageous. As long-read sequencing becomes more prevalent, integrating previous short-read datasets with new long-read data will be crucial, and Bayesian posteriors offer a principled method for this integration. Additionally, performing joint inference across all genes simultaneously, though computationally challenging, could provide significant benefits by borrowing strength across gene groups and pathways in a principled way. Future work could also apply our methods to other assays, such as ATAC-Seq, to explore allele-specific chromatin accessibility, or to combine information from transcriptomic and chromatin data to identify coordinated changes indicative of regulatory mechanisms. Additionally, the Bayesian framework offers the possibility of incorporating gene-specific priors from cohorts of control individuals, a feature that could be incorporated in future versions of the model.

## Author contributions statement

XZ, ASA, and WHM designed the study, with WHM conceiving the original idea of the Bayesian model. XZ tested, implemented, and optimized the Bayesian model, and designed and implemented the regression models, simulators, and data processing pipeline. XZ also performed all analyses on real and simulated data under the guidance of ASA and WHM. ASA provided statistical advice. TER provided insights into practical issues related to the analysis of allelic imbalance in human gene expression data. ZWG contributed to the architecture and optimization of the pipeline. XZ, ASA, and WHM wrote the manuscript.

## Acknowledgments

The authors thank Thomas Cowart for his help in dockerizing the software, and Yuncheng Duan for sharing geneset lists. This work is supported in part by funds from the following grants: R35GM150404 to WHM and RM1HG011123 to ASA, TER.

## Supplementary Tables

**Table S1:**
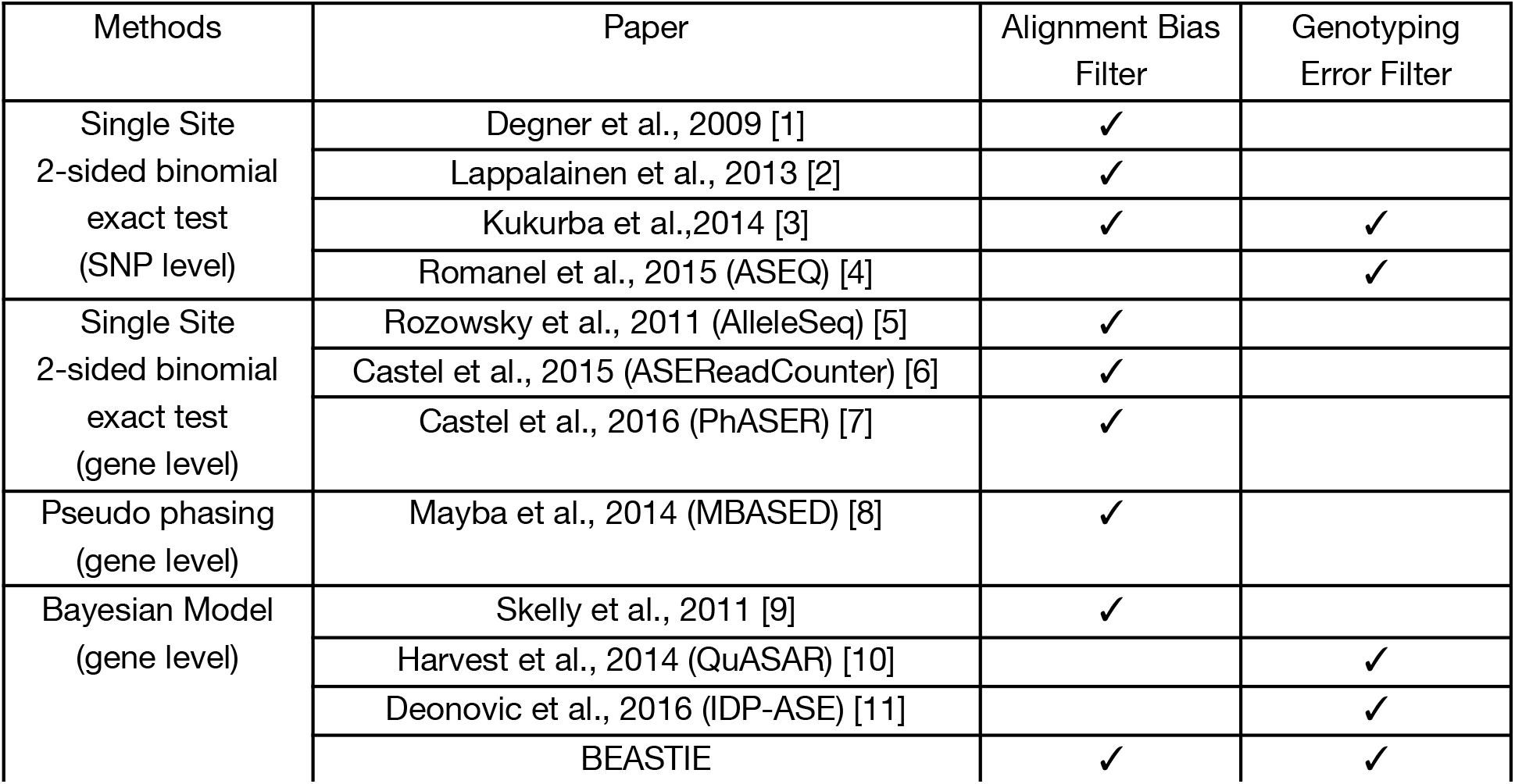
ASE Quantification Methods Used in Previous Studies.

| Methods | Paper | Alignment Bias Filter | Genotyping Error Filter |
| --- | --- | --- | --- |
| Single Site<br>2-sided binomial<br>exact test<br>(SNP level) | Degner et al., 2009 [1] | ✓ |  |
|  | Lappalainen et al., 2013 [2] | ✓ |  |
|  | Kukurba et al., 2014 [3] | ✓ | ✓ |
|  | Romanel et al., 2015 (ASEQ) [4] |  | ✓ |
| Single Site<br>2-sided binomial<br>exact test<br>(gene level) | Rozowsky et al., 2011 (AlleleSeq) [5] | ✓ |  |
|  | Castel et al., 2015 (ASEReadCounter) [6] | ✓ |  |
|  | Castel et al., 2016 (PhASER) [7] | ✓ |  |
| Pseudo phasing<br>(gene level) | Mayba et al., 2014 (MBASED) [8] | ✓ |  |
| Bayesian Model<br>(gene level) | Skelly et al., 2011 [9] | ✓ |  |
|  | Harvest et al., 2014 (QuASAR) [10] |  | ✓ |
|  | Deonovic et al., 2016 (IDP-ASE) [11] |  | ✓ |
|  | BEASTIE | ✓ | ✓ |

**Table S2.**
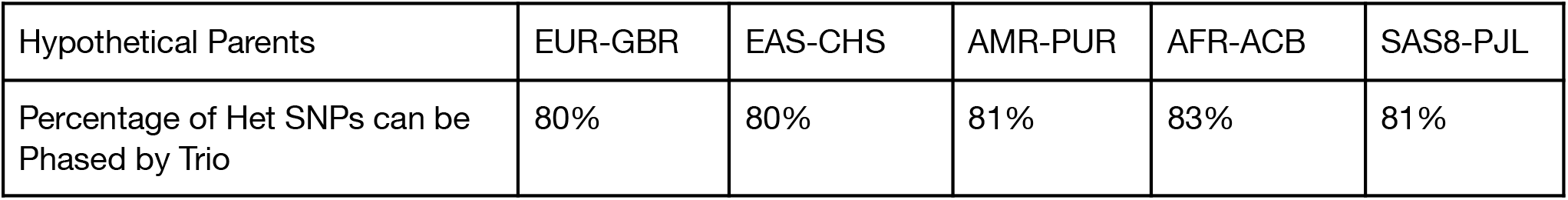
**Expected percentage of bi-allelic heterozygous SNPs that cannot be phased using trio-based phasing, in Thousand Genomes Project samples from five ancestries, based on transmission simulations**.

**Table S3:** 1000 Genome sample Used in Trio Phasing Assessment From Table S2.

| Sample | Ancestry | Sex |
| --- | --- | --- |
| HG00096 | EUR-GBR | M |
| HG00097 | EUR-GBR | F |
| HG00403 | EAS-CHS | M |
| HG00404 | EAS-CHS | F |
| HG00551 | AMR-PUR | F |
| HG00553 | AMR-PUR | M |
| HG02307 | AFR-ACB | M |
| HG02308 | AFR-ACB | F |
| HG03237 | SAS-PJL | M |
| HG03238 | SAS-PJL | F |

**Table S4.** Phasing error for each dataset. This phasing error is calculated using all phasable SNP pairs.

| Sample | Phasing method | Ancestry | Average SHAPEIT2 Switching Error per SNP pair | Shapeit2 reference panel |
| --- | --- | --- | --- | --- |
| NA12878 | GIAB gold standard VCF phasing | CEU | 4.03% (974 out of 24167 SNP pairs) | All reference panel after removing NA12878 |
| NA12878 | GIAB gold standard VCF phasing | CEU | 3.02% (731 out of 24167 SNP pairs) | European reference panel after removing NA12878 |
| GM19440 | Experimental VCF phasing | YRI | 4.02% (2074 out of 51585 SNP pairs) | African reference panel |
| NA19240 | 1000 Genome trio phasing (parents: NA19238, NA19239) | YRI | 2.2% (755 out of 34278 SNP pairs) | African reference panel |

**Table S5.** Number of hets within a gene increases with higher switching error in GIAB (NA12878). The spearman correlation coefficient between the number of hets and percentage of genes with phasing error is 0.6854692 with p value as 0.002388332 (binning genes with hets with >= 18 together and removing genes with only 1 het).

| n_distinct_hets | n_genes | n_genes_w_error | gene_w_error_percentage |
| --- | --- | --- | --- |
| 1 | 2765 | 0 | 0.000000 |
| 2 | 1577 | 117 | 7.419150 |
| 3 | 1009 | 86 | 8.523290 |
| 4 | 682 | 64 | 9.384164 |
| 5 | 420 | 36 | 8.571429 |
| 6 | 300 | 43 | 14.333333 |
| 7 | 202 | 29 | 14.356436 |
| 8 | 157 | 26 | 16.560510 |
| 9 | 108 | 10 | 9.259259 |
| 10 | 88 | 13 | 14.772727 |
| 11 | 45 | 10 | 22.222222 |
| 12 | 44 | 11 | 25.000000 |
| 13 | 17 | 6 | 35.294118 |
| 14 | 19 | 4 | 21.052632 |
| 15 | 14 | 1 | 7.142857 |
| 16 | 10 | 3 | 30.000000 |
| 17 | 12 | 3 | 25.000000 |
| 18 | 5 | 1 | 20.000000 |
| 19 | 5 | 0 | 0.000000 |
| 20 | 1 | 0 | 0.000000 |
| 21 | 1 | 0 | 0.000000 |
| 24 | 1 | 0 | 0.000000 |
| 26 | 1 | 1 | 100.000000 |
| 29 | 3 | 1 | 33.333333 |
| 30 | 1 | 1 | 100.000000 |
| 32 | 1 | 1 | 100.000000 |

**Table S6.**
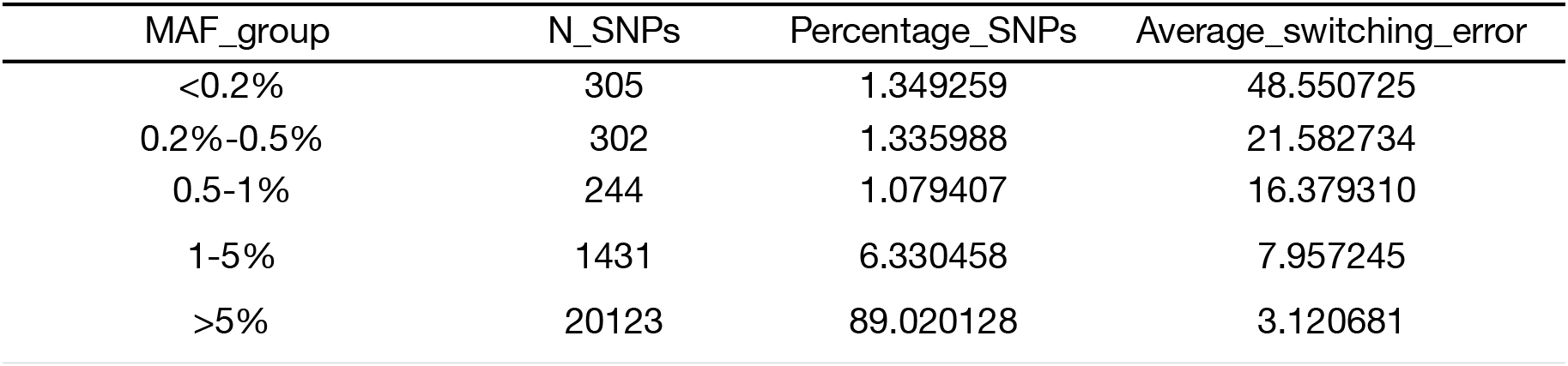
Switching error decreases with high MAF in GIAB (NA12878)

| MAF_group | N_SNP | Percentage_SNP | Average_switching_error |
| --- | --- | --- | --- |
| <0.2% | 305 | 1.349259 | 48.550725 |
| 0.2%-0.5% | 302 | 1.335988 | 21.582734 |
| 0.5-1% | 244 | 1.079407 | 16.379310 |
| 1-5% | 1431 | 6.330458 | 7.957245 |
| >5% | 20123 | 89.020128 | 3.120681 |

**Table S7.** Switching error increases with high inter-distance between SNP Pair in GIAB (NA12878)

| Log10(distance) group | N_SNP | Percentage_SNP | Average_switching_error |
| --- | --- | --- | --- |
| <1 | 9 | 0.0398142 | 0.000000 |
| 1-2 | 673 | 2.9772174 | 2.678571 |
| 2-3 | 6659 | 29.4580845 | 2.543648 |
| 3-4 | 4915 | 21.7429772 | 3.968903 |
| >4 | 2925 | 12.9396151 | 8.367769 |

**Table S8.** Switching error for each chromosome in GIAB (NA12878)

| chrN | num_SNP | percent_SNP | average_switching_error |
| --- | --- | --- | --- |
| 1 | 2192 | 9.696970 | 5.020921 |
| 2 | 1642 | 7.263880 | 3.052065 |
| 3 | 1218 | 5.388188 | 3.855721 |
| 4 | 987 | 4.366291 | 4.148784 |
| 5 | 1049 | 4.640566 | 5.825243 |
| 6 | 995 | 4.401681 | 4.097117 |
| 7 | 1036 | 4.583057 | 3.202329 |
| 8 | 851 | 3.764654 | 6.271777 |
| 9 | 761 | 3.366512 | 6.329114 |
| 10 | 929 | 4.109710 | 2.446982 |
| 11 | 1305 | 5.773059 | 3.665521 |
| 12 | 1287 | 5.693431 | 2.311436 |
| 13 | 368 | 1.627958 | 7.111111 |
| 14 | 783 | 3.463835 | 4.863813 |
| 15 | 793 | 3.508073 | 1.795332 |
| 16 | 1249 | 5.525326 | 2.863688 |
| 17 | 1383 | 6.118115 | 4.468085 |
| 18 | 523 | 2.313647 | 7.068063 |
| 19 | 1667 | 7.374475 | 4.967801 |
| 20 | 476 | 2.105729 | 2.816901 |
| 21 | 300 | 1.327140 | 1.951220 |
| 22 | 811 | 3.587702 | 4.166667 |

**Table S9.** Switching error rate decrease with high LD in GIAB (NA12878)

| **d_group**<br><fctr> | **num_SNPs**<br><int> | **percent_SNPs**<br><dbl> | **average_switching_error**<br><dbl> |
| --- | --- | --- | --- |
| <0.2 | 328 | 1.451006 | 20.743034 |
| 0.2-0.4 | 251 | 1.110374 | 19.678715 |
| 0.4-0.6 | 253 | 1.119221 | 11.067194 |
| 0.6-0.8 | 374 | 1.654501 | 8.086253 |
| >0.8 | 13354 | 59.075426 | 2.999323 |

**B**
| **r2_group**<br><fctr> | **num_SNPs**<br><int> | **percent_SNPs**<br><dbl> | **average_switching_error**<br><dbl> |
| --- | --- | --- | --- |
| <0.2 | 4741 | 20.973236 | 9.096692 |
| 0.2-0.4 | 1779 | 7.869940 | 2.317694 |
| 0.4-0.6 | 1263 | 5.587259 | 1.352426 |
| 0.6-0.8 | 1176 | 5.202389 | 1.794872 |
| >0.8 | 5601 | 24.777704 | 1.163415 |

**Table S10:** Average %(#) ASE genes detected by different methods among 445 individuals from 1000 Genome Project with different average count per het cutoff.

| ASE rate per ind %**#** | quick BEAST | Naive Sum | Major Site |
| --- | --- | --- | --- |
| average count per het $\geq 30$ | 10.00% (193) | 9.10% (176) | 6.32% (121) |
| average count per het $\geq 10$ | 7.23% (271) | 6.39% (240) | 4.37% (163) |
| average count per het $\geq 1$ | 4.26% (311) | 3.34% (245) | 2.12% (155) |

**Table S11:** Summary of 1000 Genome Project Dataset Analysis.

| n (95%CI) | Utah (CEU) | Finnish (FIN) | British (GBR) | Tuscan (TSI) | Yoruba (YRI) |
| --- | --- | --- | --- | --- | --- |
| # individual | 89 | 92 | 86 | 91 | 87 |
| RNA total sequencing depth | 58085673 | 59294884 | 59908207 | 58410071 | 62235536 |
| # genes | 3430<br>(3354,3505) | 3503<br>(3420, 3587) | 3583<br>(3504, 3662) | 3529<br>(3448, 3611) | 4387<br>(4284,4489) |
| # het snps | 10945<br>(10611,11280) | 11294<br>(10922,11665) | 11658<br>(11299,12016) | 11381<br>(11038,11724) | 15474<br>(14968,15980) |
| average # hets | 3.18 | 3.21 | 3.24 | 3.21 | 3.51 |
| # ASE genes | 267<br>(251,283) | 247<br>(233,261) | 249<br>(236,269) | 253<br>(236,269) | 342<br>(314,370) |
| % ASE genes | 7.68<br>(7.65,7.72) | 6.95<br>(6.92,6.98) | 6.86<br>(6.83,6.90) | 7.03<br>(7.00,7.07) | 7.64<br>(7.58,7.69) |

**Table S12:**
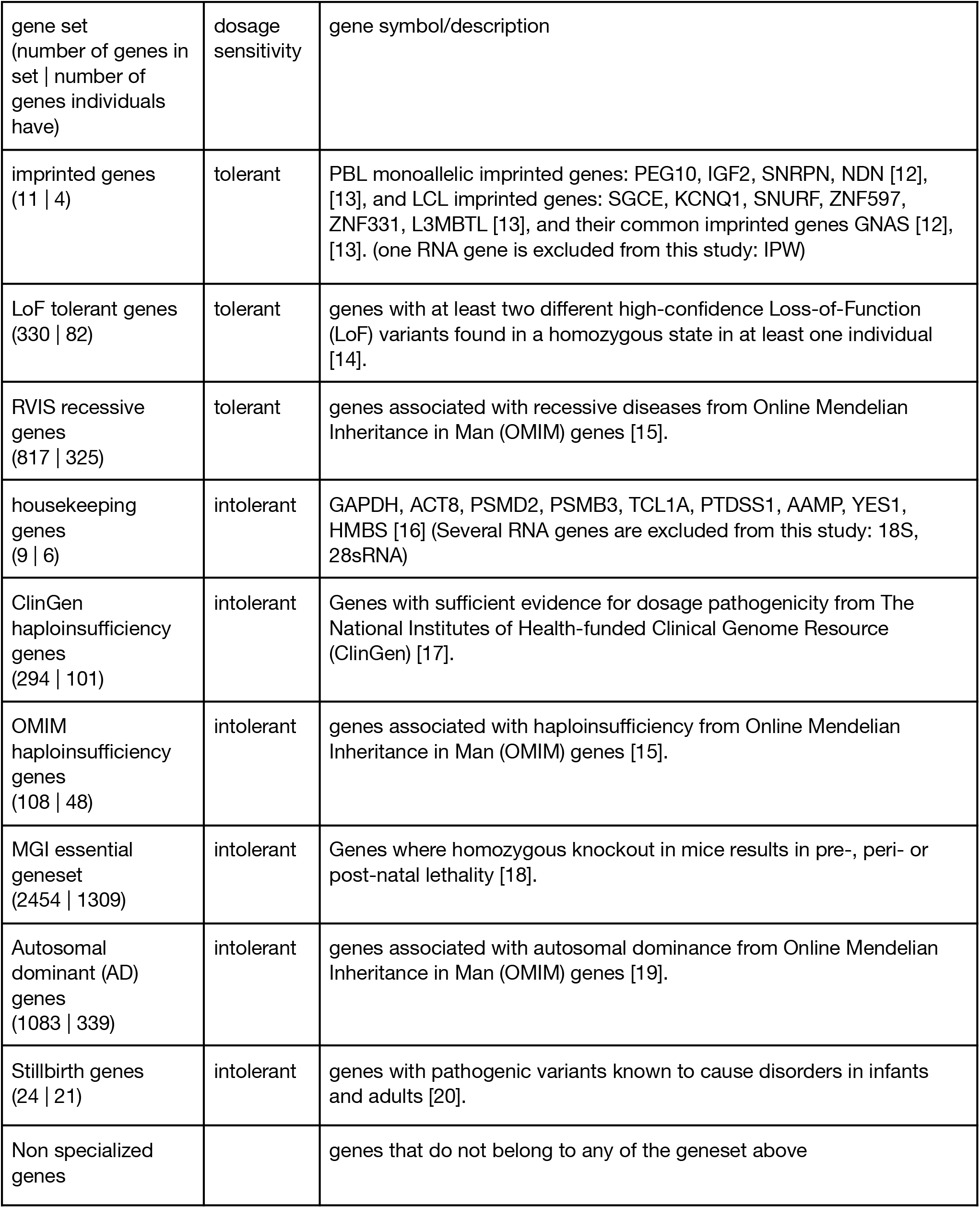
geneset description. All numbers of genes in the table above are reflecting the number of genes that have at least 10 read counts per het in at least two of the individuals per ancestry group.

**Table S13:** **ASE Rate Statistics and Wilcoxon Test P-values for Defined-Geneset genes. P-values from the Wilcoxon Signed Rank test are conducted between defined-geneset genes and non-specialized genes**.

| Geneset | Mean (Variance) of<br>ASE Rate | 95% CI of<br>ASE rate | Wilcoxon Sign Rank<br>Test P values |
| --- | --- | --- | --- |
| All genes | 6.89 (3.11) | (6.73 , 7.06) | / |
| Non-specialized geneset | 7.00 (3.09) | (6.84 , 7.17) | / |
| Imprinted | 63.8 (1027) | (60.7 , 66.9) | 1.20099e-65 |
| LoF tolerant | 13.4 (35.8) | (12.8 , 14.0) | 1.186255e-61 |
| Housekeeping | 21.6 (706) | (19.0 , 24.1) | 4.91415e-16 |
| Recessive | 5.98 (7.29) | (5.73 , 6.23) | 1.394209e-24 |
| Clingen haploinsufficient | 3.97 (9.83) | (3.68 , 4.26) | 1.85044e-55 |
| OMIM haploinsufficient | 5.29 (25.6) | (4.82 , 5.76) | 1.142098e-16 |
| All haploinsufficient | 4.02 (9.26) | (3.74 , 4.30) | 1.01169e-56 |
| MGI essential | 6.21 (4.11) | (6.21 , 6.73) | 6.690214e-35 |
| Autosomal Dominance (AD) | 6.47 (7.77) | (6.21 , 6.73) | 5.334317e-08 |
| Still birth | 1.64 (20.1) | (1.22 , 2.06) | 4.036315e-52 |

**Table S14:**
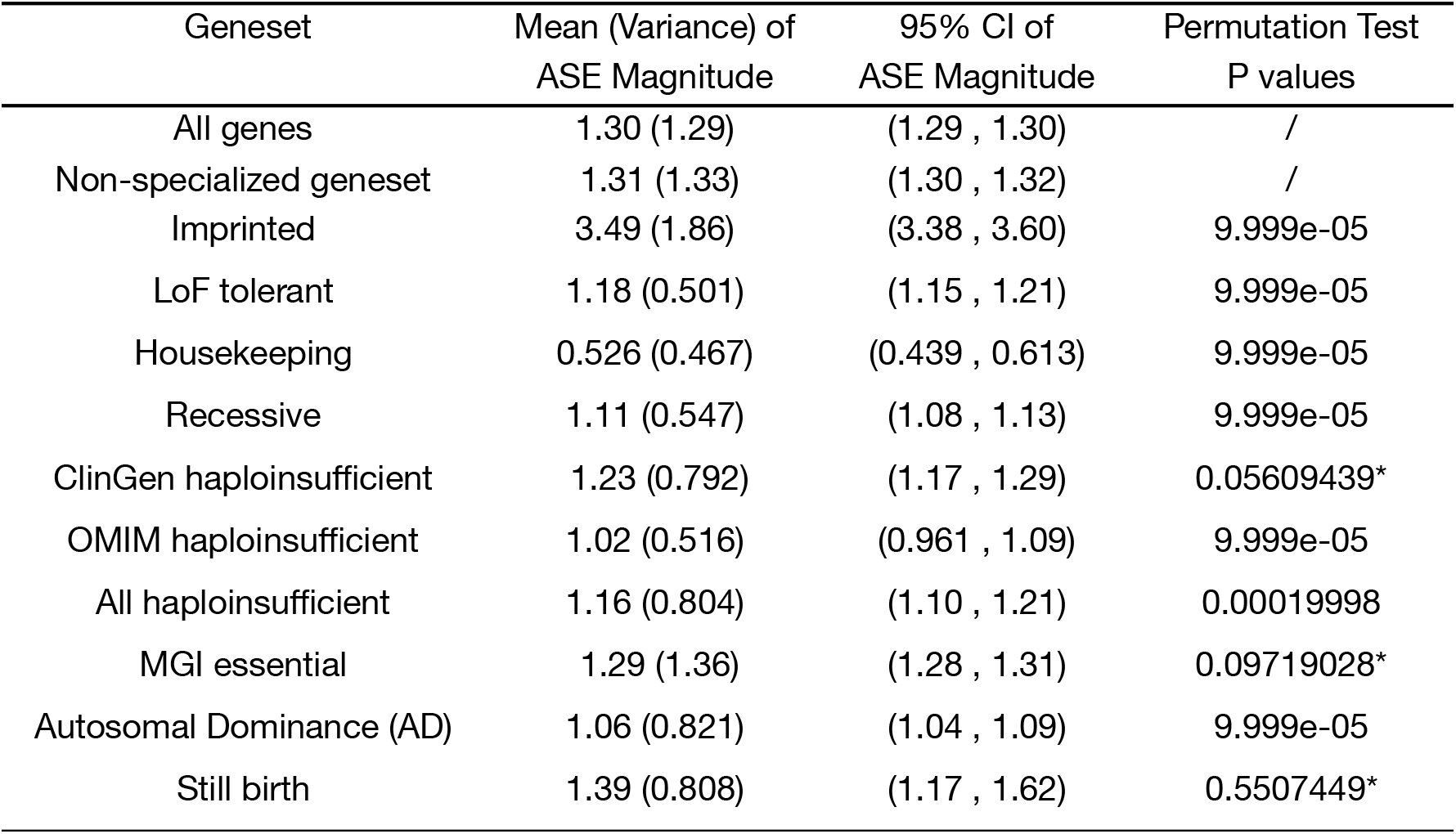
**ASE magnitude Statistics and Permutation Test P-values for Defined-Geneset ASE genes. Permutation Test is conducted between ASE genes from defined-genesets and non-specialized genes. * indicates non-signifiance**.

| Geneset | Mean (Variance) of<br>ASE Magnitude | 95% CI of<br>ASE Magnitude | Permutation Test<br>P values |
| --- | --- | --- | --- |
| All genes | 1.30 (1.29) | (1.29 , 1.30) | / |
| Non-specialized geneset | 1.31 (1.33) | (1.30 , 1.32) | / |
| Imprinted | 3.49 (1.86) | (3.38 , 3.60) | 9.999e-05 |
| LoF tolerant | 1.18 (0.501) | (1.15 , 1.21) | 9.999e-05 |
| Housekeeping | 0.526 (0.467) | (0.439 , 0.613) | 9.999e-05 |
| Recessive | 1.11 (0.547) | (1.08 , 1.13) | 9.999e-05 |
| ClinGen haploinsufficient | 1.23 (0.792) | (1.17 , 1.29) | 0.05609439* |
| OMIM haploinsufficient | 1.02 (0.516) | (0.961 , 1.09) | 9.999e-05 |
| All haploinsufficient | 1.16 (0.804) | (1.10 , 1.21) | 0.00019998 |
| MGI essential | 1.29 (1.36) | (1.28 , 1.31) | 0.09719028* |
| Autosomal Dominance (AD) | 1.06 (0.821) | (1.04 , 1.09) | 9.999e-05 |
| Still birth | 1.39 (0.808) | (1.17 , 1.62) | 0.5507449* |

**Table S15:** ASE magnitude Statistics and Permutation Test P-values for Defined-Geneset non-ASE genes.

| Geneset | Mean (Variance) of<br>ASE Magnitude | 95% CI of<br>ASE Magnitude |
| --- | --- | --- |
| All genes | 0.251 (0.0394) | (0.250 , 0.251) |
| Non-specialized geneset | 0.252 (0.0398) | (0.252 , 0.253) |
| Imprinted | 0.213 (0.0564) | (0.186 , 0.241) |
| LoF tolerant | 0.265 (0.0421) | (0.262 , 0.269) |
| Housekeeping | 0.249 (0.0357) | (0.236 , 0.261) |
| Recessive | 0.250 (0.0397) | (0.248 , 0.252) |
| Clingen haploinsufficient | 0.243 (0.0379) | (0.240 , 0.246) |
| OMIM haploinsufficient | 0.241 (0.0384) | (0.236 , 0.245) |
| All haploinsufficient | 0.238 (0.0374) | (0.236 , 0.241) |
| MGI essential | 0.241 (0.0370) | (0.240 , 0.242) |
| Autosomal Dominance (AD) | 0.241 (0.0384) | (0.239 , 0.242) |
| Still birth | 0.251 (0.0374) | (0.245 , 0.257) |

**Table S16:** Top 25 genes identified by ‘prioritization method’ statistics. The ranking order is determined by using the ratio of individuals with extreme ASE and the ratio of individuals with ASE per gene. ASE status is determined by FDR adjusted p-values from quickBEAST with a significance threshold of 0.05. And each gene has at least 10 read depth per het. There are 3 pseudogenes (MTRNR2L12, UBBP4, WASH4P), 2 RNA genes (FLVCR1-DT, AC021317.19), 6 immune gene segments (IGHV3-9, IGHV3-20, IGKV3-15, IGLV3-1, IGHV3-21, IGLC7).

|  | gene_name | gene_ID | avg_hets | total | num_ASE | num_extremeASE | extremeASE_ratio | ASE_ratio |
| --- | --- | --- | --- | --- | --- | --- | --- | --- |
| 1 | SNRPN | ENSG00000128739 | 1.124424 | 217 | 217 | 217 | 1.0000000 | 1.00000000 |
| 2 | MTRNR2L12 | ENSG00000269028 | 1.315068 | 146 | 146 | 146 | 1.0000000 | 1.00000000 |
| 3 | PEG10 | ENSG00000242265 | 2.276753 | 271 | 271 | 271 | 1.0000000 | 1.00000000 |
| 4 | POTEJ | ENSG00000222038 | 1.000000 | 15 | 15 | 15 | 1.0000000 | 1.00000000 |
| 5 | NAP1L5 | ENSG00000177432 | 1.406593 | 91 | 91 | 91 | 1.0000000 | 1.00000000 |
| 6 | EIF5AL1 | ENSG00000253626 | 1.825000 | 40 | 40 | 40 | 1.0000000 | 1.00000000 |
| 7 | HLA-DQA2 | ENSG00000237541 | 1.697674 | 86 | 86 | 86 | 1.0000000 | 1.00000000 |
| 8 | UBBP4 | ENSG00000263563 | 1.000000 | 59 | 59 | 59 | 1.0000000 | 1.00000000 |
| 9 | FAM50B | ENSG00000145945 | 1.141414 | 99 | 98 | 99 | 1.0000000 | 0.98989899 |
| 10 | LPAR6 | ENSG00000139679 | 1.051546 | 97 | 95 | 97 | 1.0000000 | 0.97938144 |
| 11 | UTS2 | ENSG00000049247 | 1.576923 | 26 | 25 | 26 | 1.0000000 | 0.96153846 |
| 12 | IGHV3-9 | ENSG00000211940 | 1.117647 | 34 | 32 | 34 | 1.0000000 | 0.94117647 |
| 13 | IGHV3-20 | ENSG00000211946 | 1.312500 | 48 | 45 | 48 | 1.0000000 | 0.93750000 |
| 14 | FLVCR1-DT | ENSG00000198468 | 2.165714 | 175 | 146 | 174 | 0.9942857 | 0.83428571 |
| 15 | GEMIN2 | ENSG00000092208 | 1.969163 | 227 | 196 | 224 | 0.9867841 | 0.86343612 |
| 16 | AC021317.19 | ENSG00000174093 | 1.126697 | 221 | 216 | 218 | 0.9864253 | 0.97737557 |
| 17 | ZDBF2 | ENSG00000204186 | 4.637037 | 135 | 132 | 133 | 0.9851852 | 0.97777778 |
| 18 | PDE4DIP | ENSG00000178104 | 9.114754 | 61 | 58 | 60 | 0.9836066 | 0.95081967 |
| 19 | IGKV3-15 | ENSG00000244437 | 1.162791 | 43 | 37 | 42 | 0.9767442 | 0.86046512 |
| 20 | AP3S1 | ENSG00000177879 | 1.936508 | 378 | 368 | 369 | 0.9761905 | 0.97354497 |
| 21 | IGLV3-1 | ENSG00000211673 | 1.225000 | 40 | 37 | 39 | 0.9750000 | 0.92500000 |
| 22 | IGHV3-21 | ENSG00000211947 | 1.145161 | 62 | 59 | 60 | 0.9677419 | 0.95161290 |
| 23 | IGLC7 | ENSG00000211685 | 1.032967 | 91 | 81 | 88 | 0.9670330 | 0.89010989 |
| 24 | FAM153A | ENSG00000170074 | 3.035714 | 28 | 26 | 27 | 0.9642857 | 0.92857143 |
| 25 | WASH4P | ENSG00000234769 | 1.178571 | 56 | 41 | 54 | 0.9642857 | 0.73214286 |

**Table S17:**
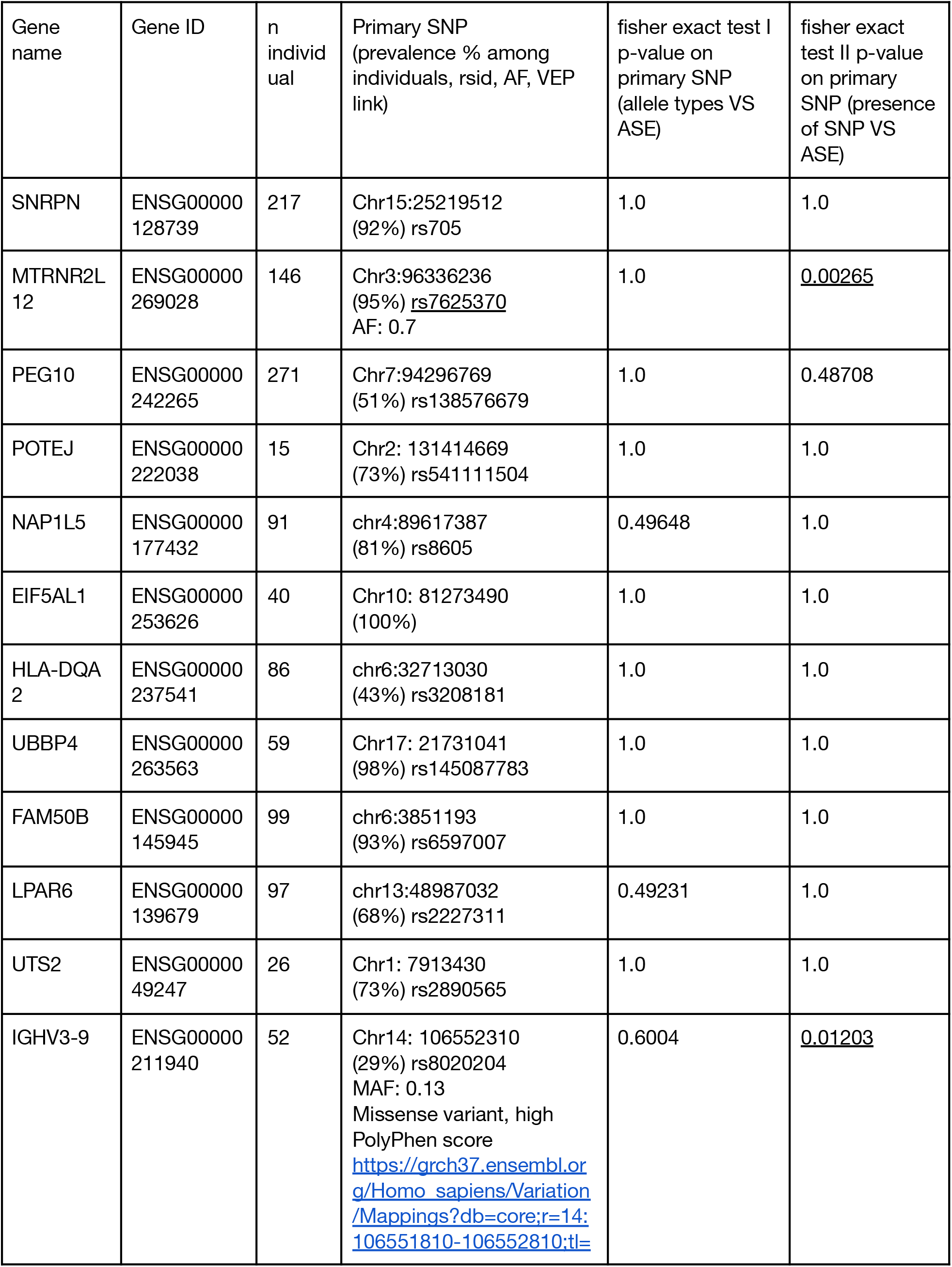
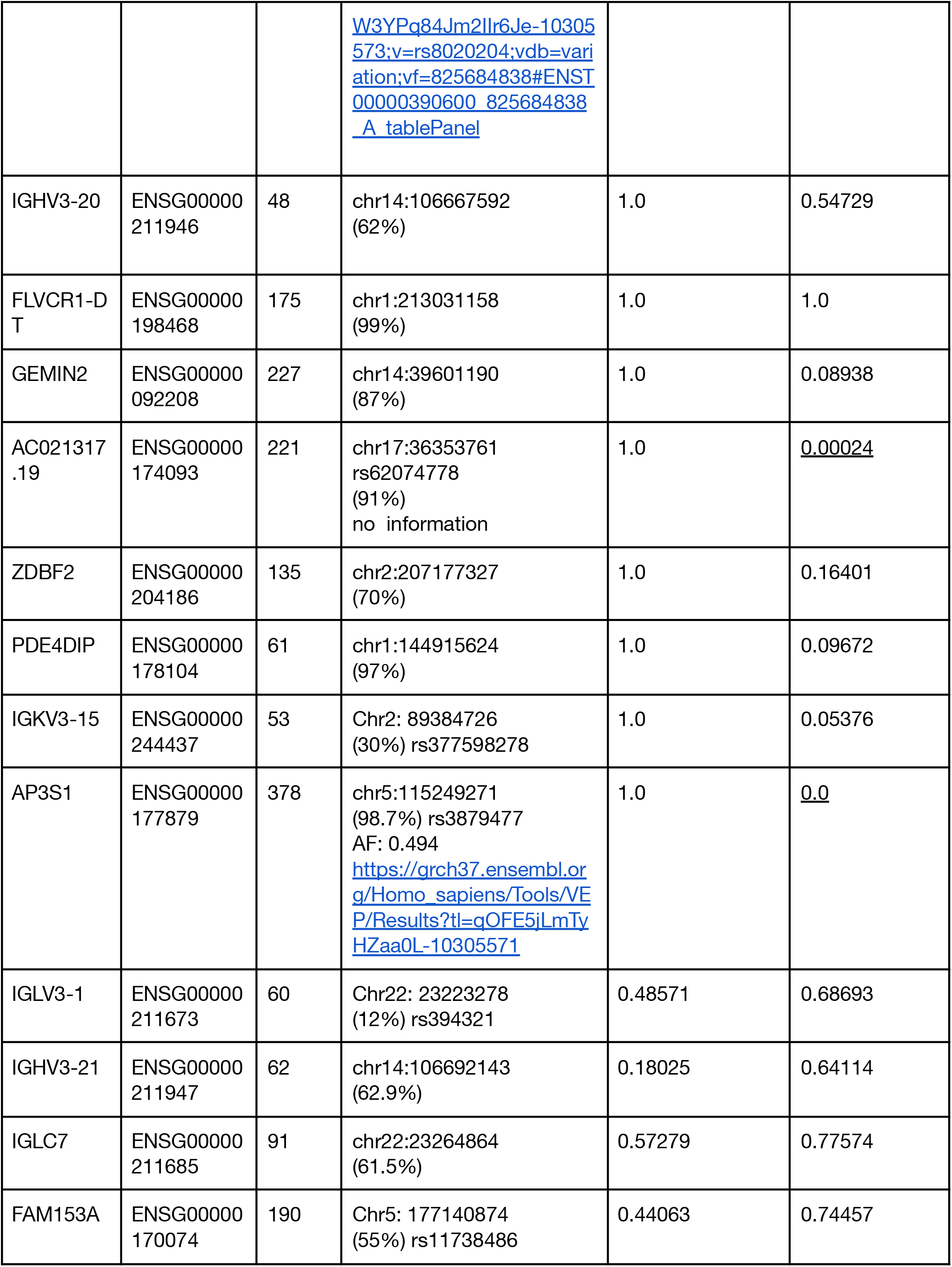
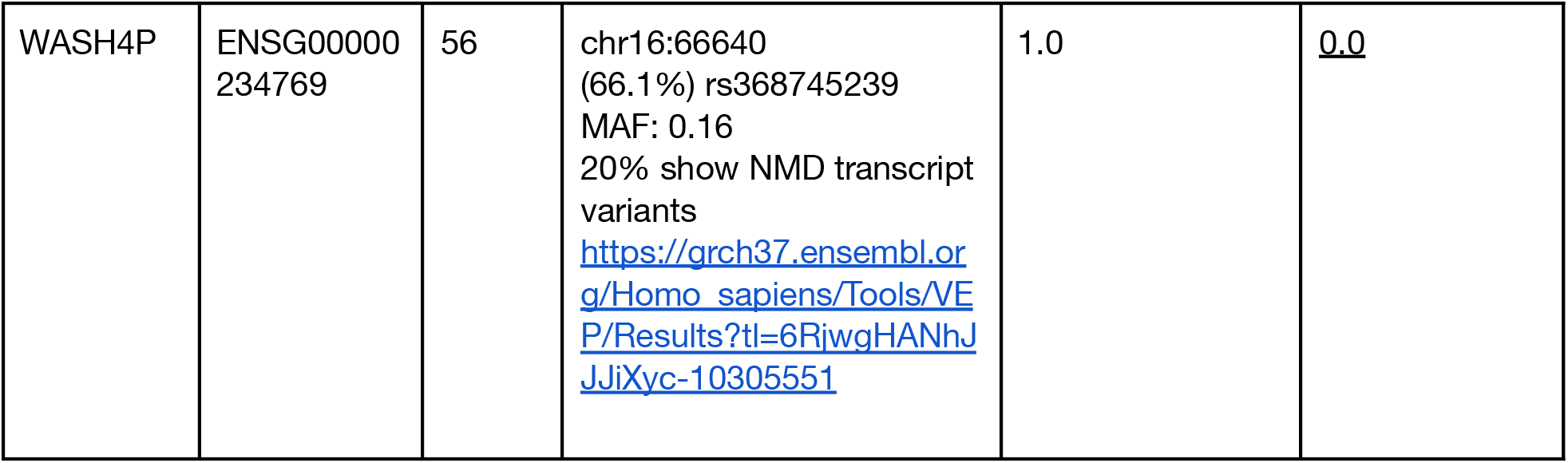
Top 30 genes identified by ‘prioritization method’ fisher exact tests. Each SNP has to have at least 10 reads, and shared in at least 1 individual. ‘n_individual’ indicates the number of individuals having at least 10 reads on a given SNP.

**Table S18:**
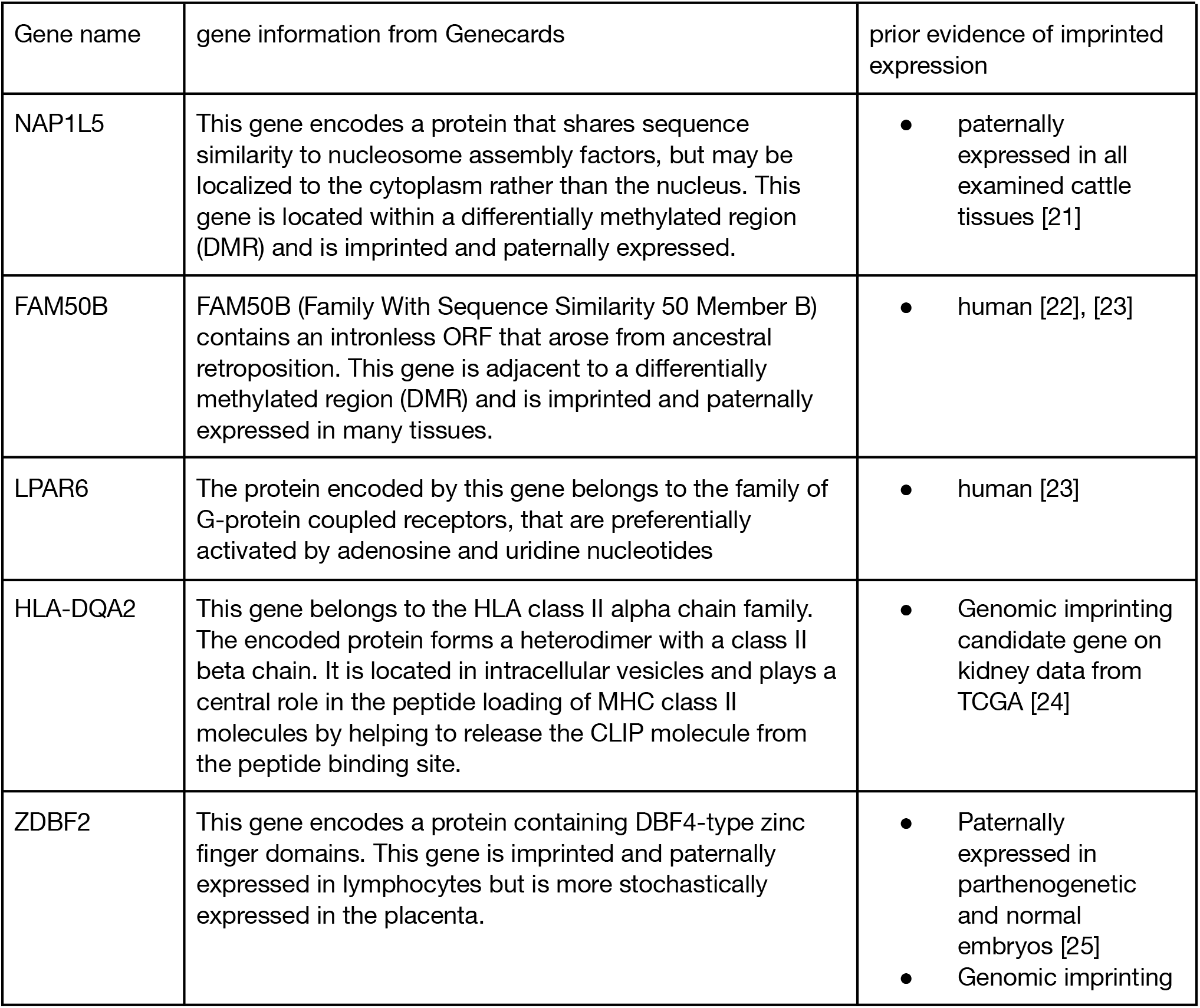
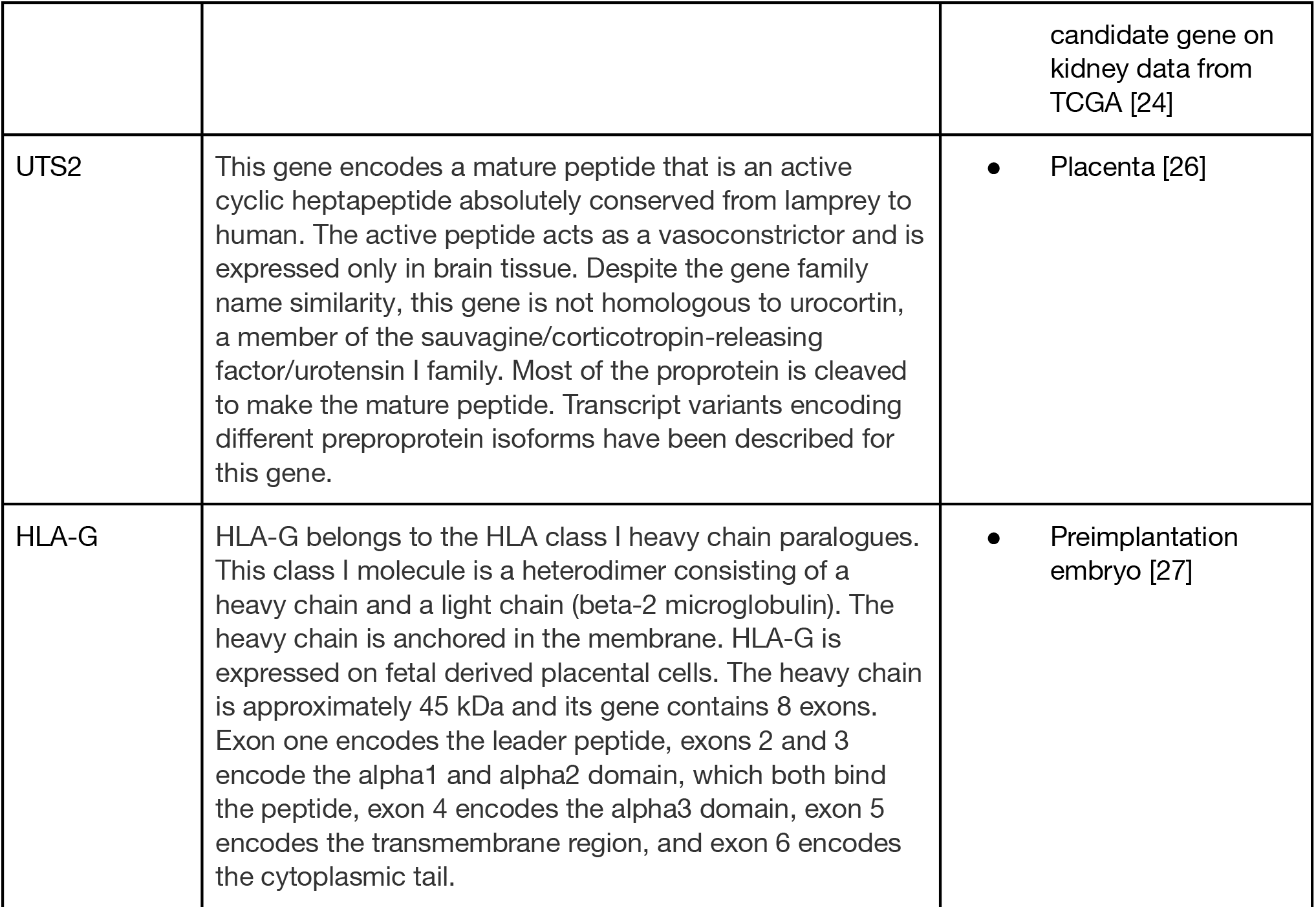
7 imprinted gene candidates reported in previous publication. There are among the top 30 genes from Table S16 arranged based on decreasing order of extreme ASE ratio, and ASE ratio. And all genes are having at least 10 read counts on average of each het.

**Table S19:**
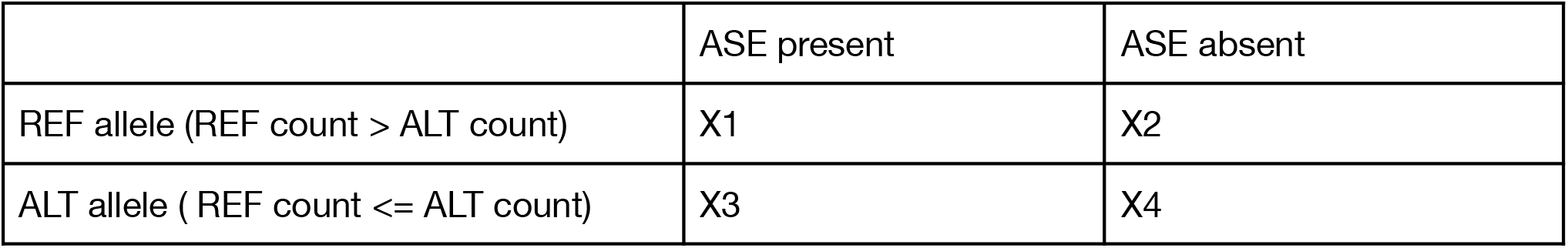
Two by two table examining the association between Allele Type and ASE status. The table displays the number of individuals with alleles that have higher count in genes with different ASE status. Fisher’s Exact Test was applied to assess the statistical significance of this association in **Table S16**.

|  |  |  |
| --- | --- | --- |
|  | ASE present | ASE absent |
| REF allele (REF count > ALT count) | X1 | X2 |
| ALT allele ( REF count <= ALT count) | X3 | X4 |

**Table S20.**
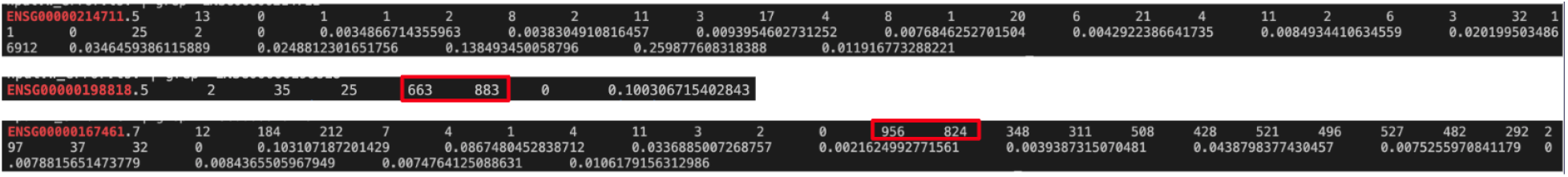
**Three genes in model input format used (in Supplementary Figure S15) to investigate the difference between fixing first site and fixing highest coverage site in qb** **(A) gene ENSG00000214711** **(B) gene ENSG00000198818** **(C) gene ENSG00000167461**

**Table S21:** variables with non-zero coefficients in fitted model with regularization. We use R function cva.glmnet and family “binomial” with 10 fold cross validation for 10 times on full model in training dataset and calculate the average alpha 0.7163 as the best. And then we use R function cv.glmnet and family “binomial” with 10 fold cross validation and best alpha in training dataset again to find the best lambda of 0.000326700507361396. This fitted model is then applied to testing dataset for switching error prediction.

| Variables used in fitted model | coefficient |
| --- | --- |
| intercept | -4.1059315 |
| Minimum MAF in SNP pair | -6.0251976 |
| Difference in MAF between SNP pair | 1.8274357 |
| Log10(inter distance in SNP pair) | 0.5546364 |
| r2 | -0.4825783 |
| Minimum MAF in SNP pair × Log10(inter distance in SNP pair) | 0.7363362 |
| Minimum MAF in SNP pair × d | -1.8066884 |
| Difference in MAF between SNP pair × r2 | 2.0147905 |
| r2 × d | -0.6216824 |
| Minimum MAF in SNP pair × Difference in MAF between SNP pair × Log10(inter distance in SNP pair) | 1.5263432 |
| Minimum MAF in SNP pair × Difference in MAF between SNP pair × d | -18.8610889 |
| Minimum MAF in SNP pair × Difference in MAF between SNP pair × Log10(inter distance in SNP pair) × r2×d | 0.8808806 |

**Table S22:** Parameters in Parameterized Simulator.

| $\pi$ | $\theta$ | n | m | d |
| --- | --- | --- | --- | --- |
| Rate of switching errors | Amount of ASE | Number of genes | Number of heterozygous site | Read depth per het site |

**Table S23.** Definitions of the variables in JAGS model.

| Variable | Definition |
| --- | --- |
| p | Probability of having major haplotype allele count at zero-count het SNP j |
| $\alpha$ | Sum of minor haplotype counts from all non zero-count sites at gene i |
| $\beta$ | Sum of minor haplotype counts from all non zero-count sites at gene i |
| n | Total count at zero-count het SNP j |
| X | Predicted major haplotype allele count at zero-count het SNP j |
| var | Variance of total count from all het SNPs genomewide |
| mu | Mean of total count from all het SNPs genomewide |

**Table S24.** Type 1 Error/Power Comparison Among Three Different Models using 1000 Stimulated Genes with 5% phasing error and 10 hets per gene. Fisher’s Exact test and the beta binomial test do not control Type 1 Error, but that the GEM model does

| Test | total count: 100<br>allele frequency: 0.5 | total count: 100<br>allele frequency: 0.9 | total count: 100<br>allele frequency: 0.1 |
| --- | --- | --- | --- |
| Fisher Type1 | 0.088 | 0.162 | 0.021 |
| Betabino Type1 | 0.125 | 0.162 | 0.057 |
| New model Type1 | 0.061 | 0.019 | 0.032 |
| JAGS Type1 | 0.036 | 0.019 | 0.049 |
| Fisher Power | 0.945 | 0.99 | 0.276 |
| Betabino Power | 0.954 | 0.99 | 0.398 |
| New model Power | 0.935 | 0.98 | 0.323 |
| JAGS Power | 0.908 | 0.979 | 0.416 |

**B**
| Test | total count: 5<br>allele frequency: 0.5 | total count: 5<br>allele frequency: 0.9 | total count: 5<br>allele frequency: 0.1 |
| --- | --- | --- | --- |
| Fisher Type1 | 0.039 | 0.003 | 0.064 |
| Betabino Type1 | 0.086 | 0.012 | 0.255 |
| New model Type1 | 0.007 | 0.0 | 0.037 |
| JAGS Type1 | 0.002 | 0.0 | 0.039 |
| Fisher Power | 0.709 | 0.957 | 0.167 |
| Betabino Power | 0.883 | 0.975 | 0.675 |
| New model Power | 0.637 | 0.92 | 0.131 |
| JAGS Power | 0.65 | 0.859 | 0.405 |

**Table S25:** Type 1 Error and power between qb, NS, MS and Pseudo Phasing. Type 1 error is using p values calculated from skewed t distribution fitted to 19M simulation data (calculation details in **Supplementary Text 4.1**) under the null with 5% phasing error and high coverage (10 hets per gene, read count 100 per het). The version of BEASTIE used here is quickBEAST.

|  | Power | Type 1 Error |
| --- | --- | --- |
| BEASTIE | 1.000 | 0.061 |
| Naive Sum | 0.921 | 0.046 |
| Major Site | 0.903 | 0.035 |
| Pseudo Phasing | 1.000 | 0.926 |

**Table S26:** Observed Type 1 error and expected type 1 error with different alpha values in BEASTIE. Type 1 error is calculated using p values from skewed t distribution fitted to 19M simulation data ((calculation details in **Supplementary Text 4.1 (2)**) under the null with 5% phasing error and high coverage (10 hets per gene, read count 100 per het). The version of BEASTIE used here is quickBEAST.

| Alpha | Observed Type 1 error | Expected Type 1 error | Absolute difference |
| --- | --- | --- | --- |
| 5e-02 | 6.058631e-02 | 5e-02 | 1.058631e-02 |
| 5e-03 | 7.686211e-03 | 5e-03 | 2.686211e-03 |
| 5e-04 | 9.272225e-04 | 5e-04 | 4.272225e-04 |
| 5e-05 | 1.084970e-04 | 5e-05 | 5.849700e-05 |
| 5e-07 | 1.457733e-06 | 5e-07 | 9.577332e-07 |

## Supplementary Figures

**Figure S1.**
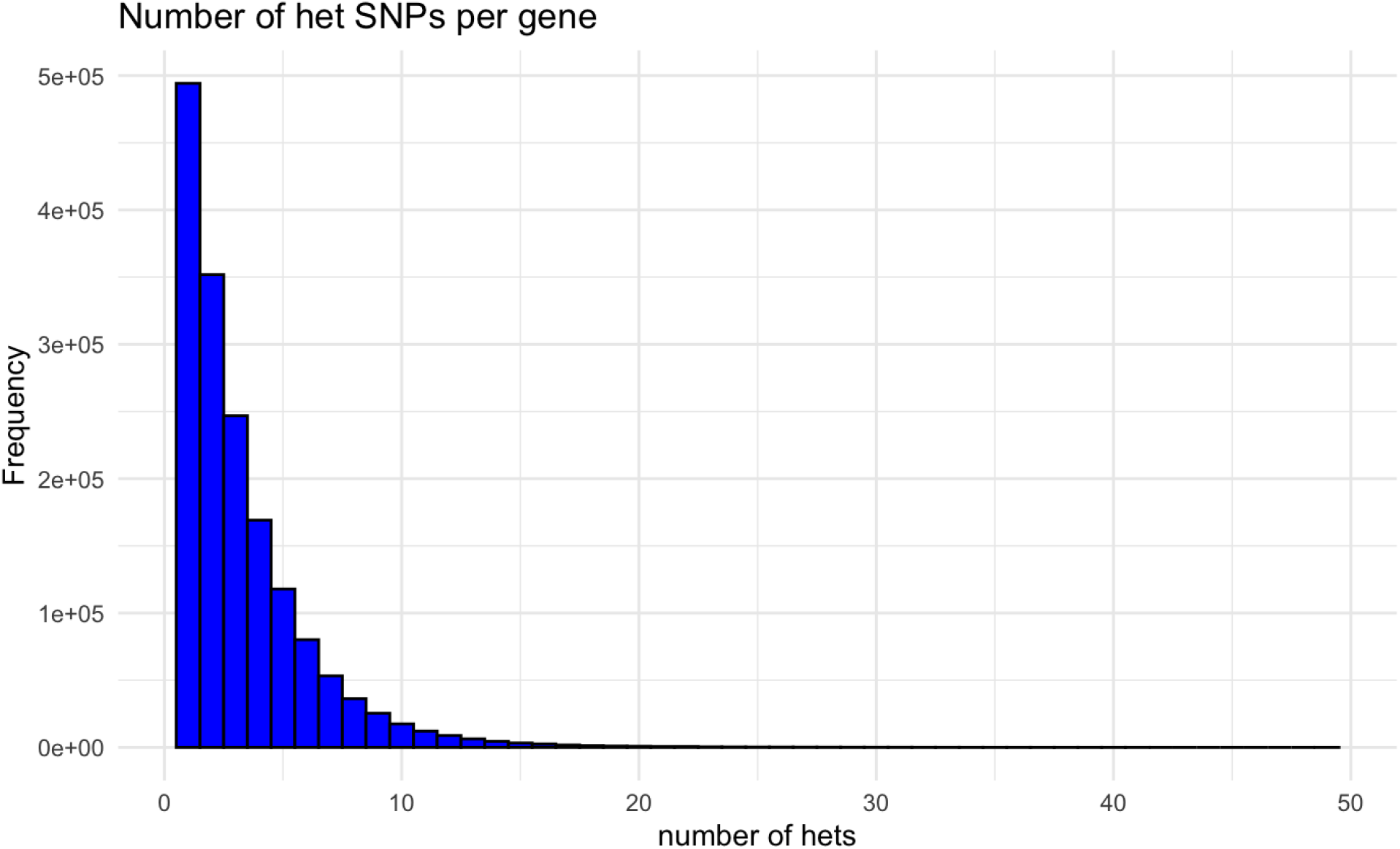
density plot of the number of exonic het SNPs per gene on 1000 genome samples.

**Figure S2.**
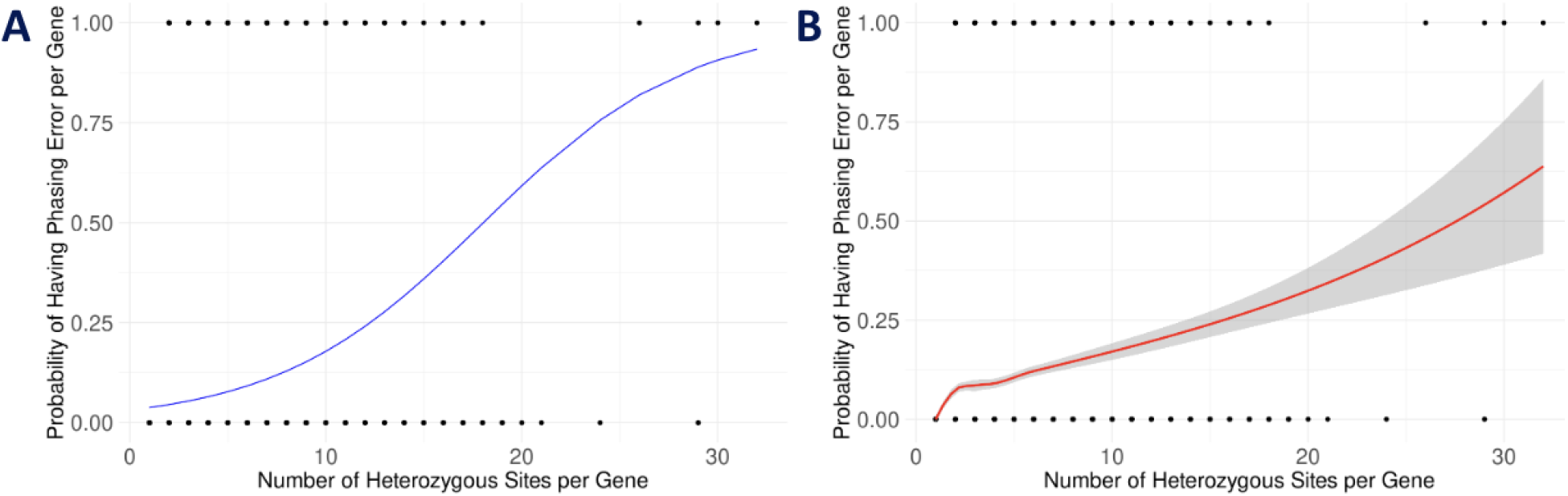
**The relationship between the number of heterozygous site per gene and the probability of having phasing error based on gold standard phasing from GIAB data**. **(A) Logistic Regression curve** **(B) LOWESS curve**

**Figure S3.**
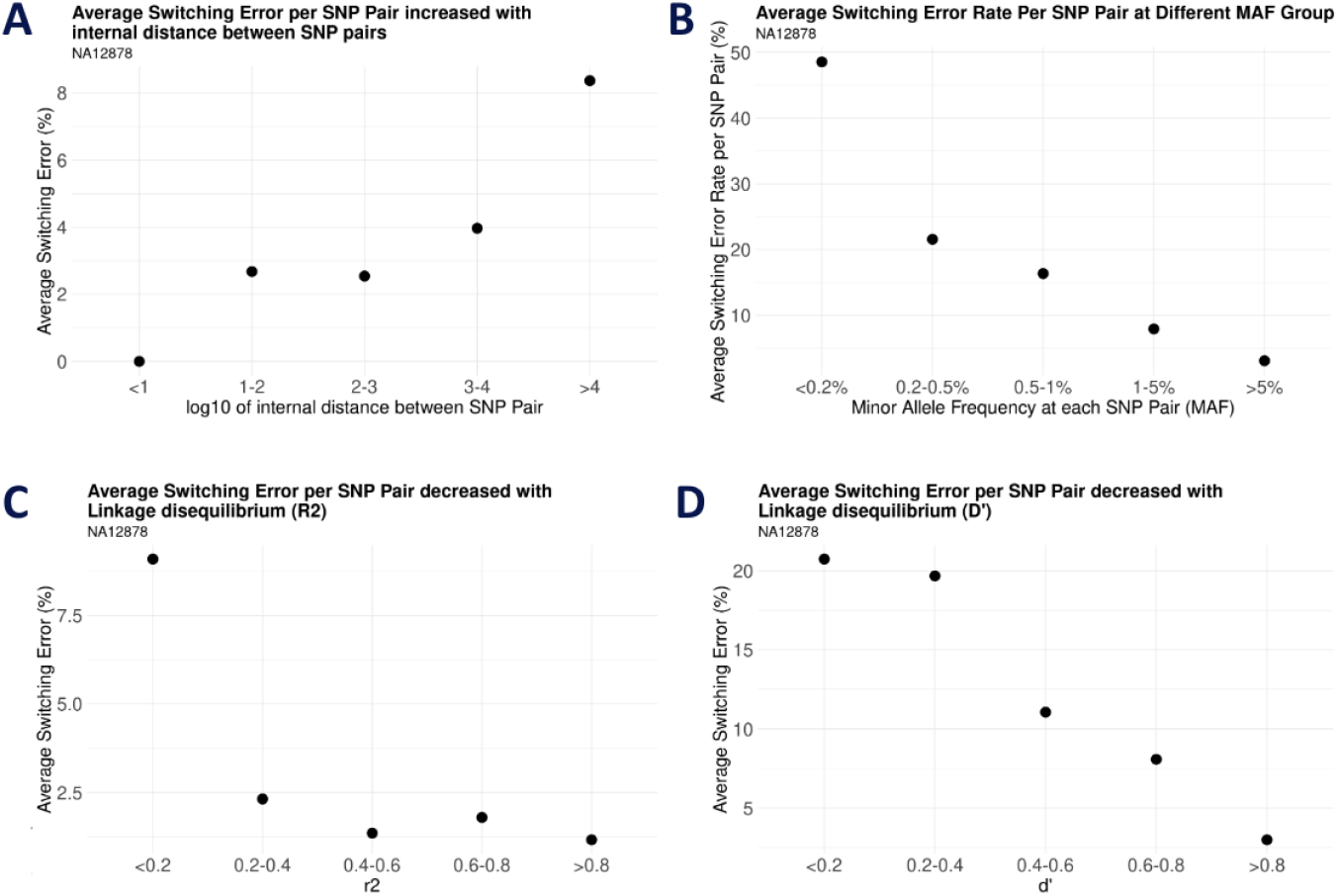
relationship of each variables (in SELR) with switching error. (A) Switching error increases with high inter-SNP pair distance (B) Switching error decreases with high MAF (C) Switching error decreases with high LD values (r2) (D) Switching error decreases with high LD values (d’)

**Figure S4.**
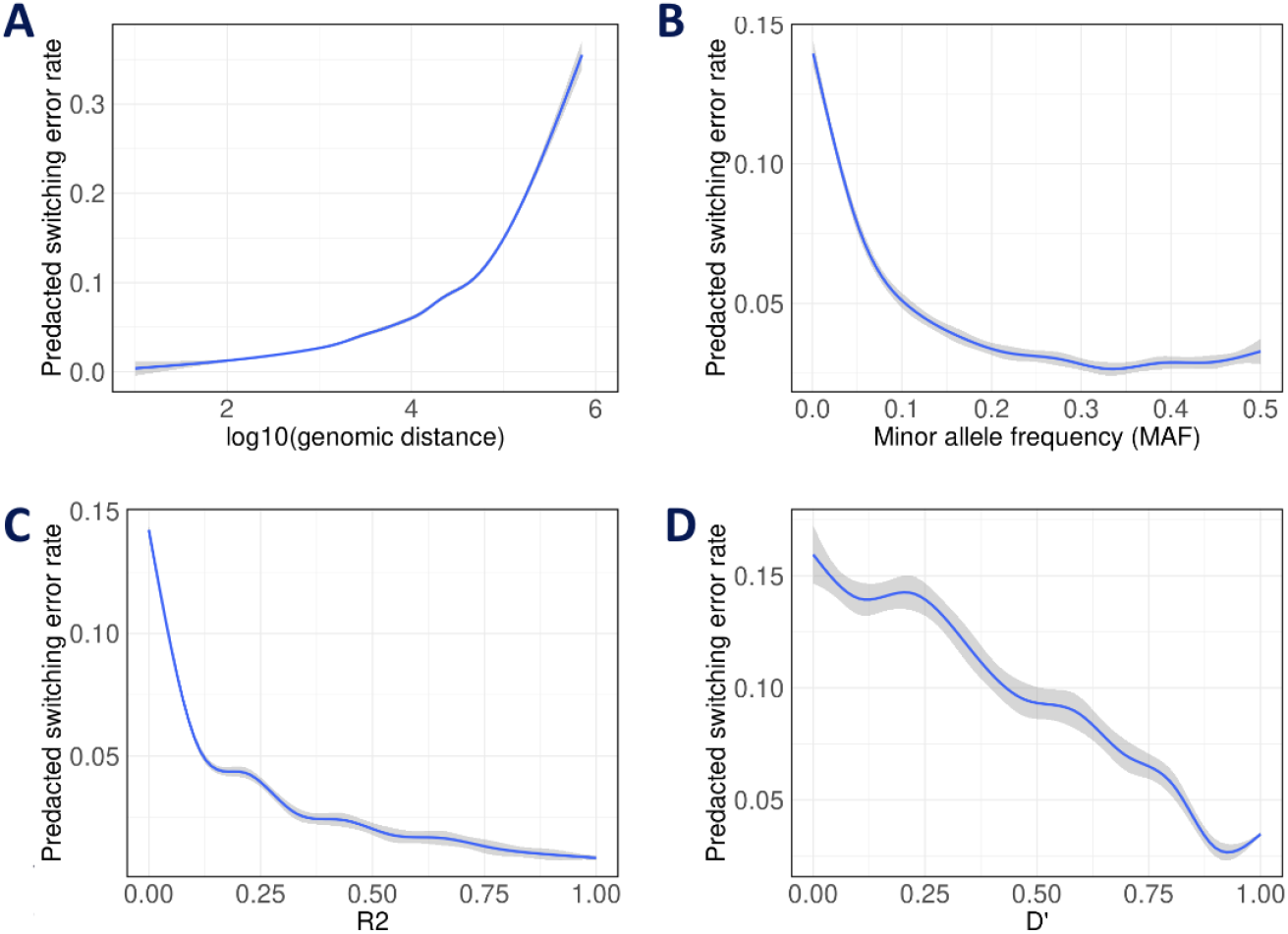
GAM smoothed line plot visualizing the relationship of each variables (in SWLR) with predicted switching error. (A) Predicted switching error increases with high inter-SNP pair distance between a SNP pair (B) Predicted switching error decreases with high min_MAF (minimum MAF between a SNP pair) (C) Predicted switching error decreases with high LD values (r2) (D) Predicted switching error decreases with high LD values (d’)

**Figure S5.**
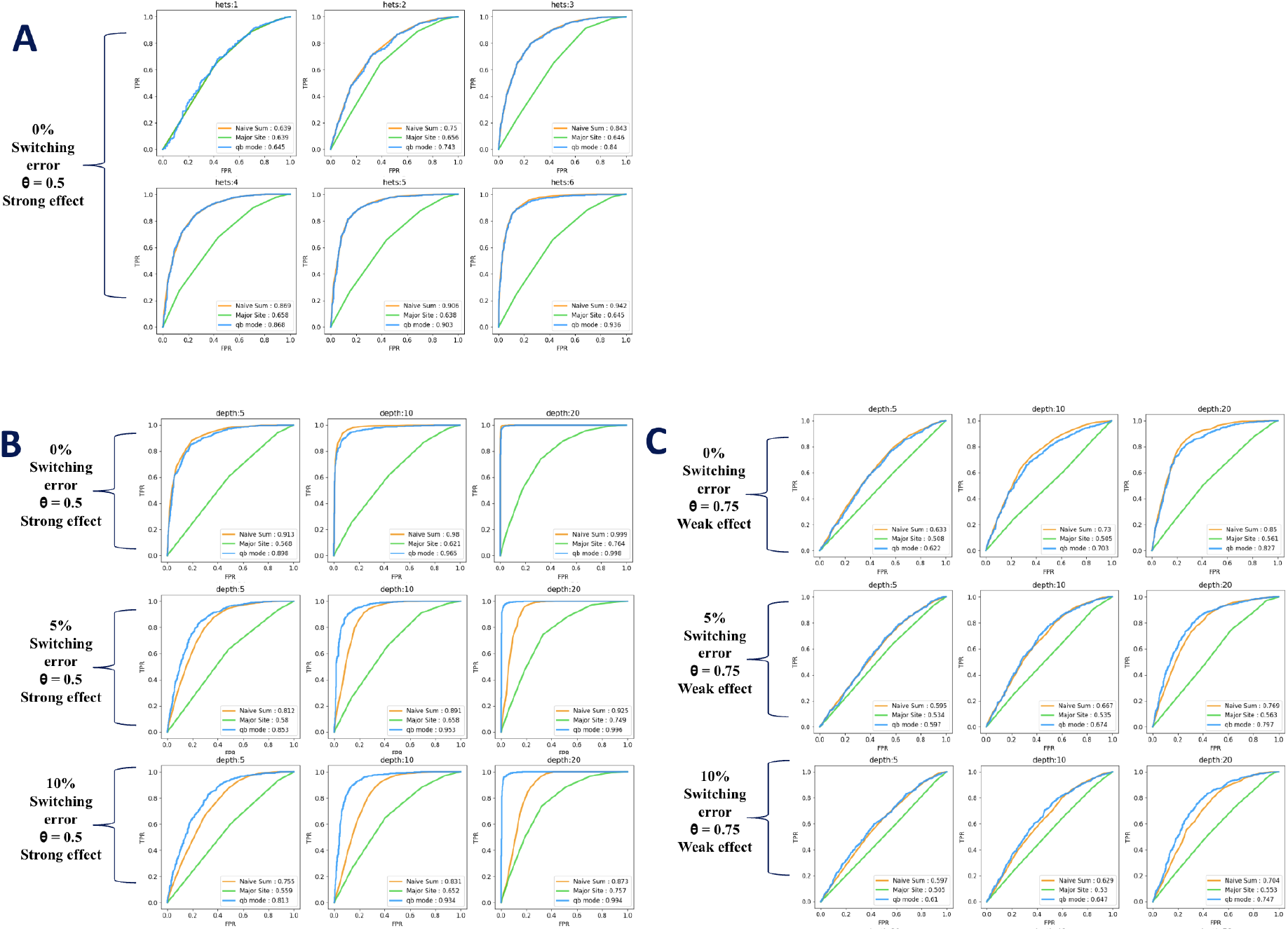
**ROC curves for BEASTIE with default 5% phasing error and two baselines based on the binomial test (NaïveSum: naively summing across sites; MajorSite: using the site with highest total count) using simulated data by parametrized simulator. All results were generated using an optimized version of BEASTIE described in Supplementary Text Section 2.2**. **a. ROC curves on simulated data with 0% switching error rate, 10 reads per site, for 1,2,3,4,5,6 heterozygous sites per gene**, **at strong ASE effect (θ=0.5)**. **b. ROC curves on simulated data with 0%, 5%, 10% switching error rate, 10 heterozygous sites per gene, for 5,10,20 reads per site**, **at strong ASE effect (θ=0.5)**. **c. ROC curves on simulated data with 0%, 5%, 10% switching error rate, 10 heterozygous sites per gene, for 5,10,20 reads per site, at weak ASE effect (θ=0.75)**. **All panels: numbers in legend are AUC values. Simulated data have 1000 genes.**

**Figure S6.**
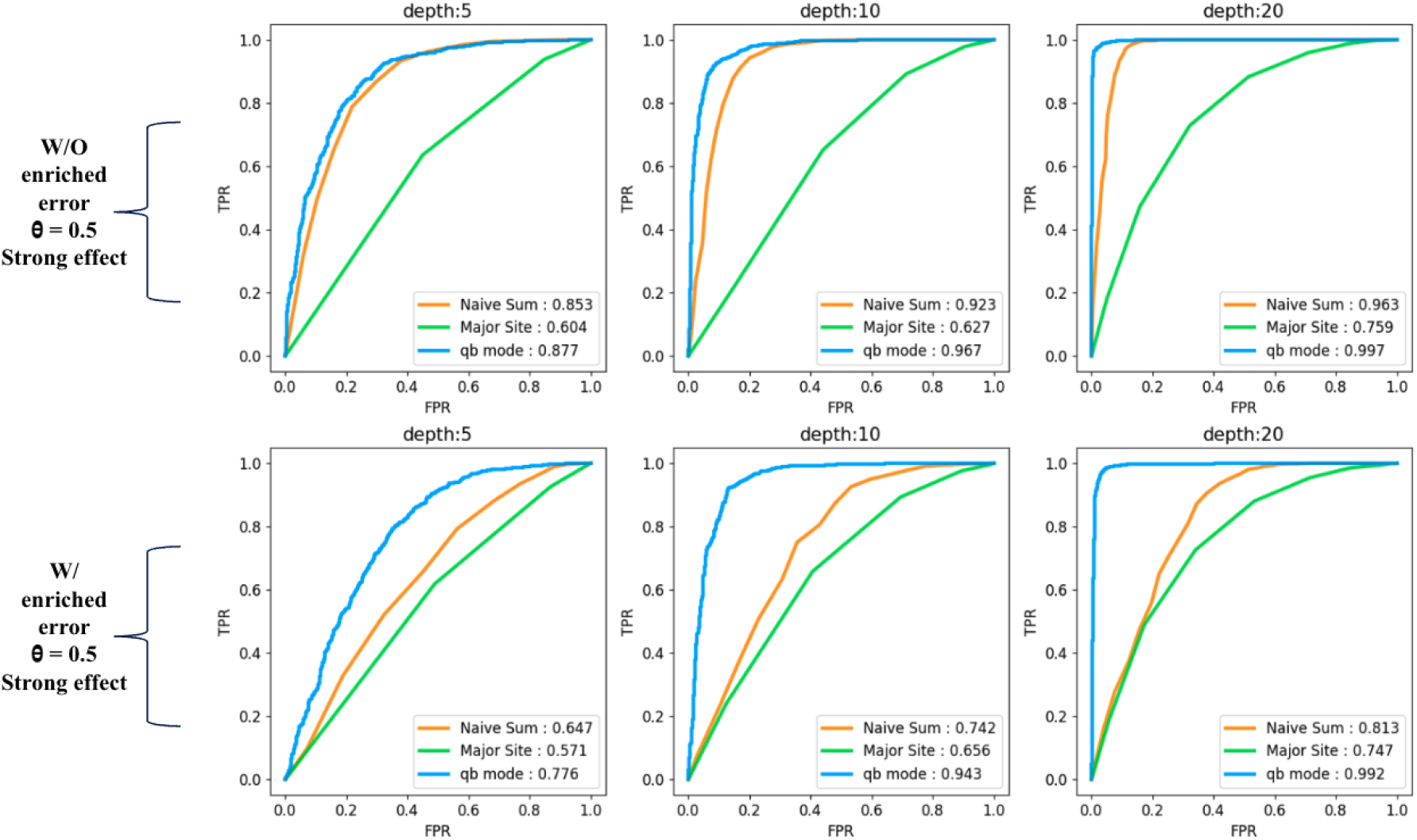
**ROC curves for BEASTIE with predicted phasing error, BEASTIE with default 5% phasing error, and two baselines using simulated data by semi-empirical simulator. All results were generated using an optimized version of BEASTIE described in Supplementary Text Section 2.2** (A) ROC curves on simulated data based on real GIAB data that has ∼3.8% switching error rate, 10 heterozygous sites per gene, for 5,10,20 reads per site, at a strong ASE effect (θ=0.5). (B) ROC curves on simulated data sampled from real GIAB data with every gene has odd number of switching error that has 12% switching error rate (predicted by model fitted with GIAB real data), 10 heterozygous sites per gene, for 5,10,20 reads per site, at a strong ASE effect (θ=0.5). All panels: numbers in legend are AUC values. Empirical simulated data have 1000 genes. Semi-empirical simulators use GIAB data to generate 1000 genes with 10 heterozygous sites per gene.

**Figure S7.**
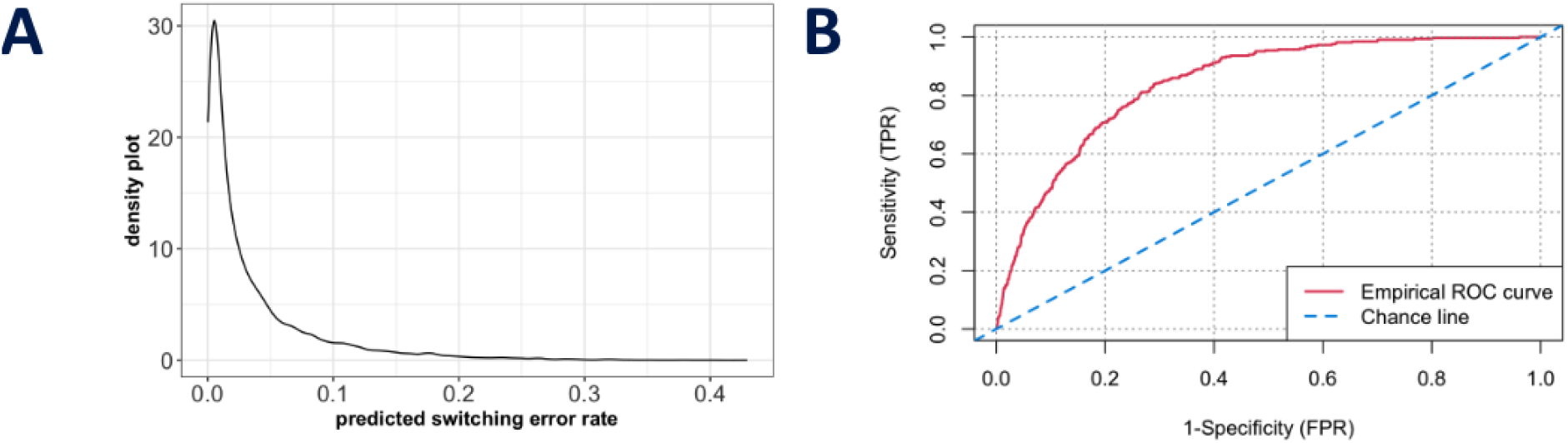
SELR model performance. (A) Scatterplot plot of predicted SHAPEIT2 switching error rate (by SELR) on sample GIAB (NA12878) as evaluated against GIAB gold standard phasing. (B) ROC curve for classification of genes having ASE in test dataset (AUC = 0.8425). The model was trained on 8,729 site pairs and validated against a disjoint set of 8,730 site pairs, each set comprising approximately 3.7% incorrectly phased site pairs.

**Figure S8.**
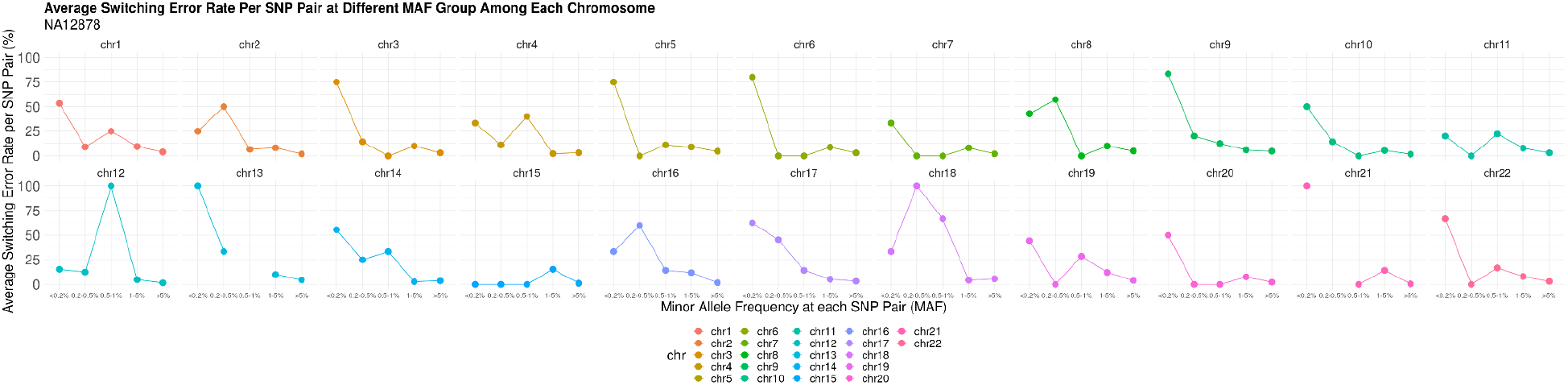
Switching error decreases with high MAF for each chromosome.

**Figure S9.**
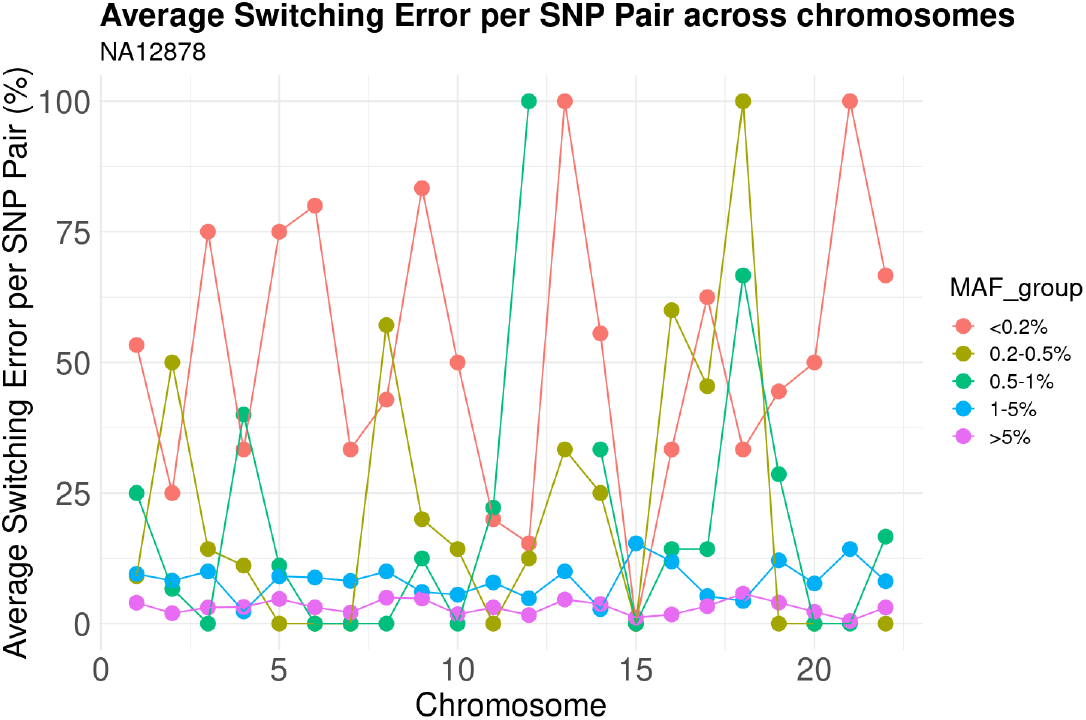
Switching error decreases with high MAF across chromosomes.

**Figure S10.**
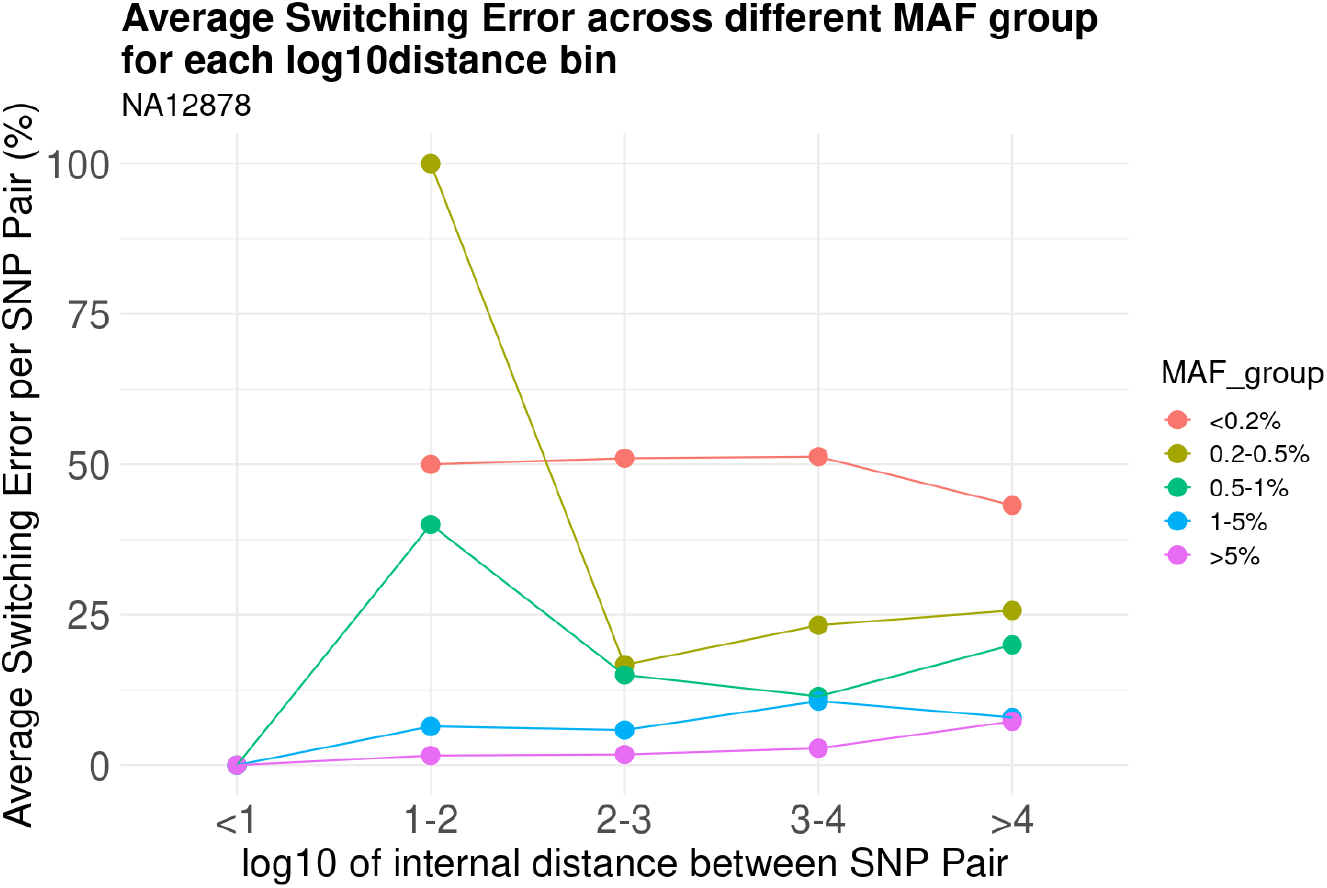
Switching error decreases with high MAF across inter-distance per SNP Pair.

**Figure S11.**
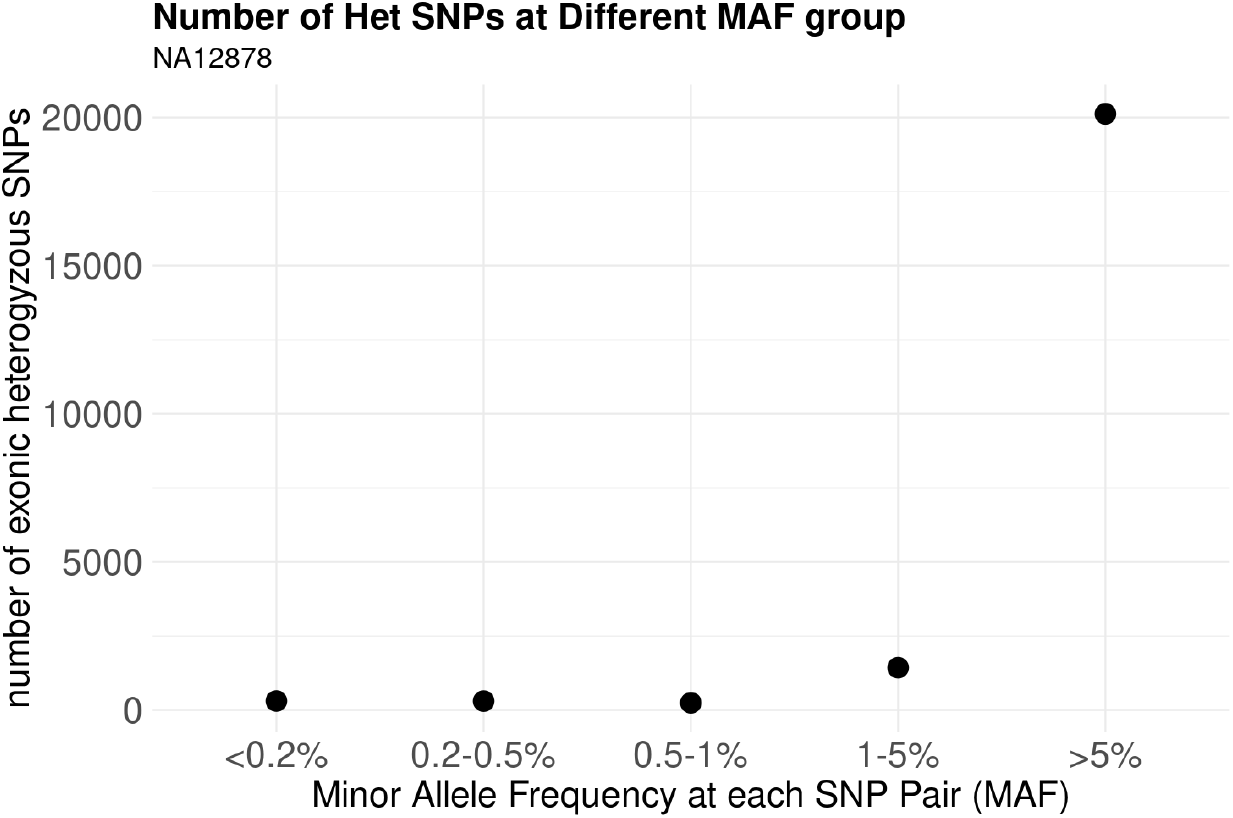
number of SNPs increases with high MAF.

**Figure S12:**
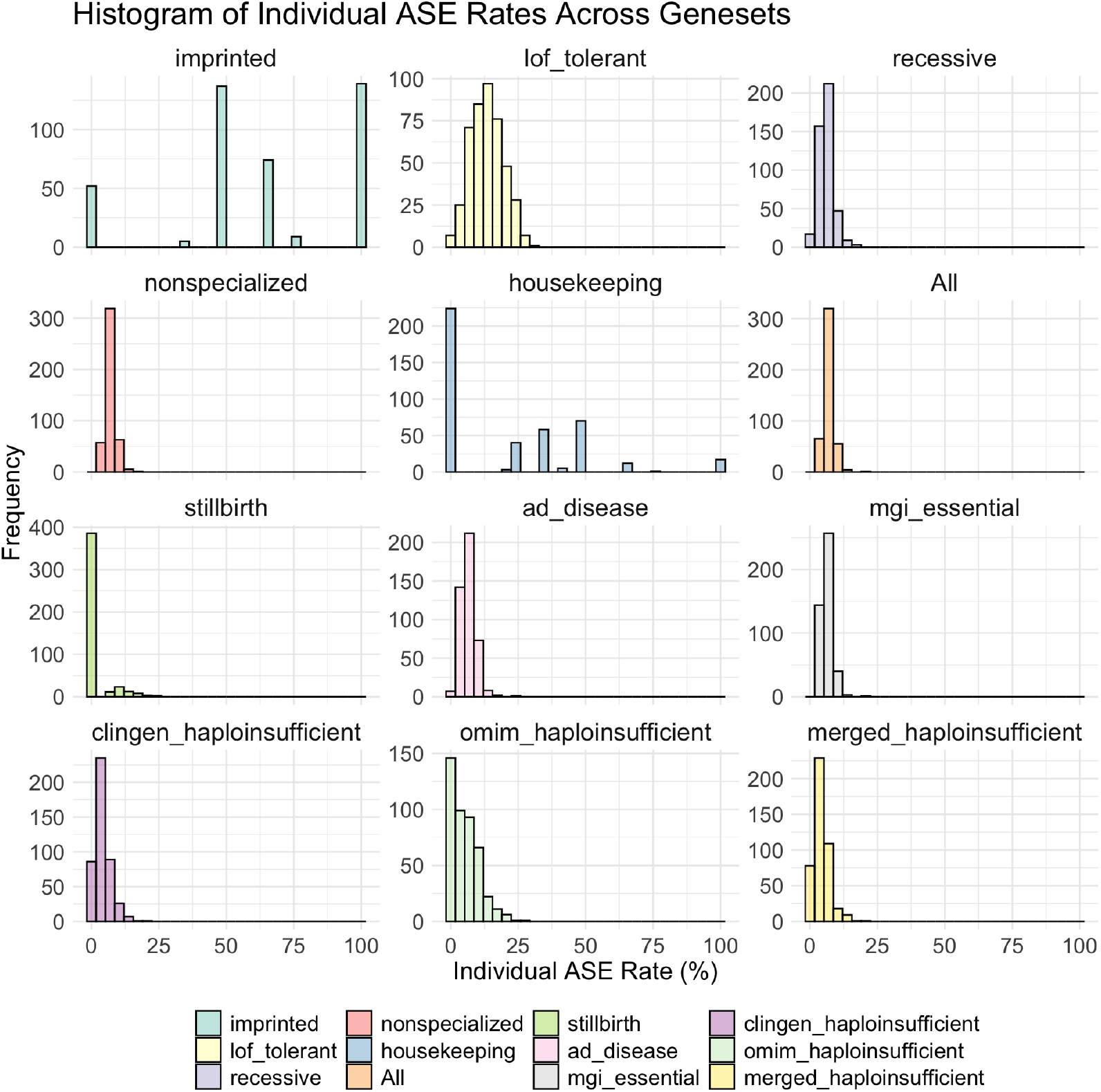
ASE Rate Histograms for all genes from different genesets.

**Figure S13:**
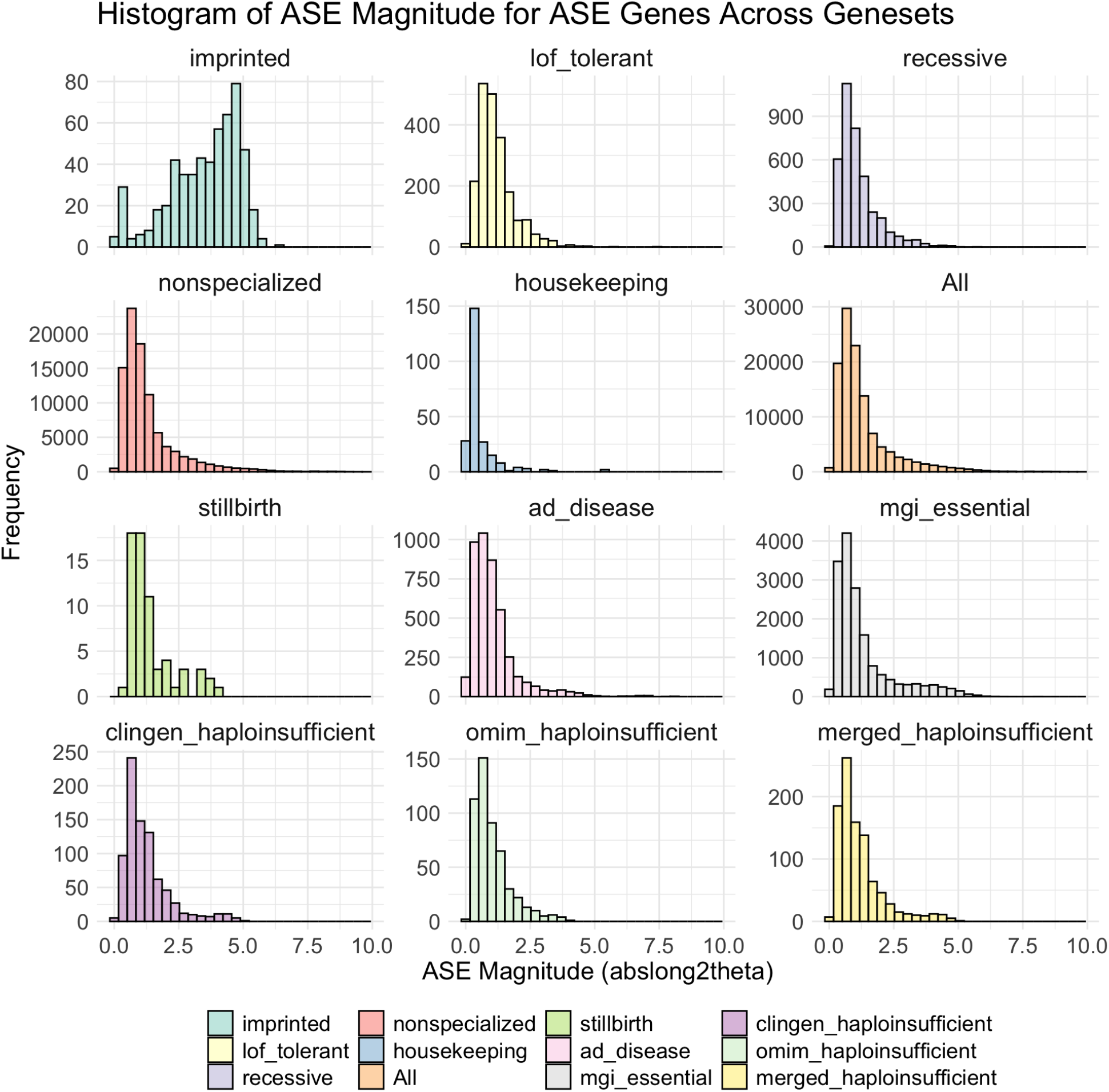
ASE magnitude histograms in ASE genes from different genesets.

**Figure S14:**
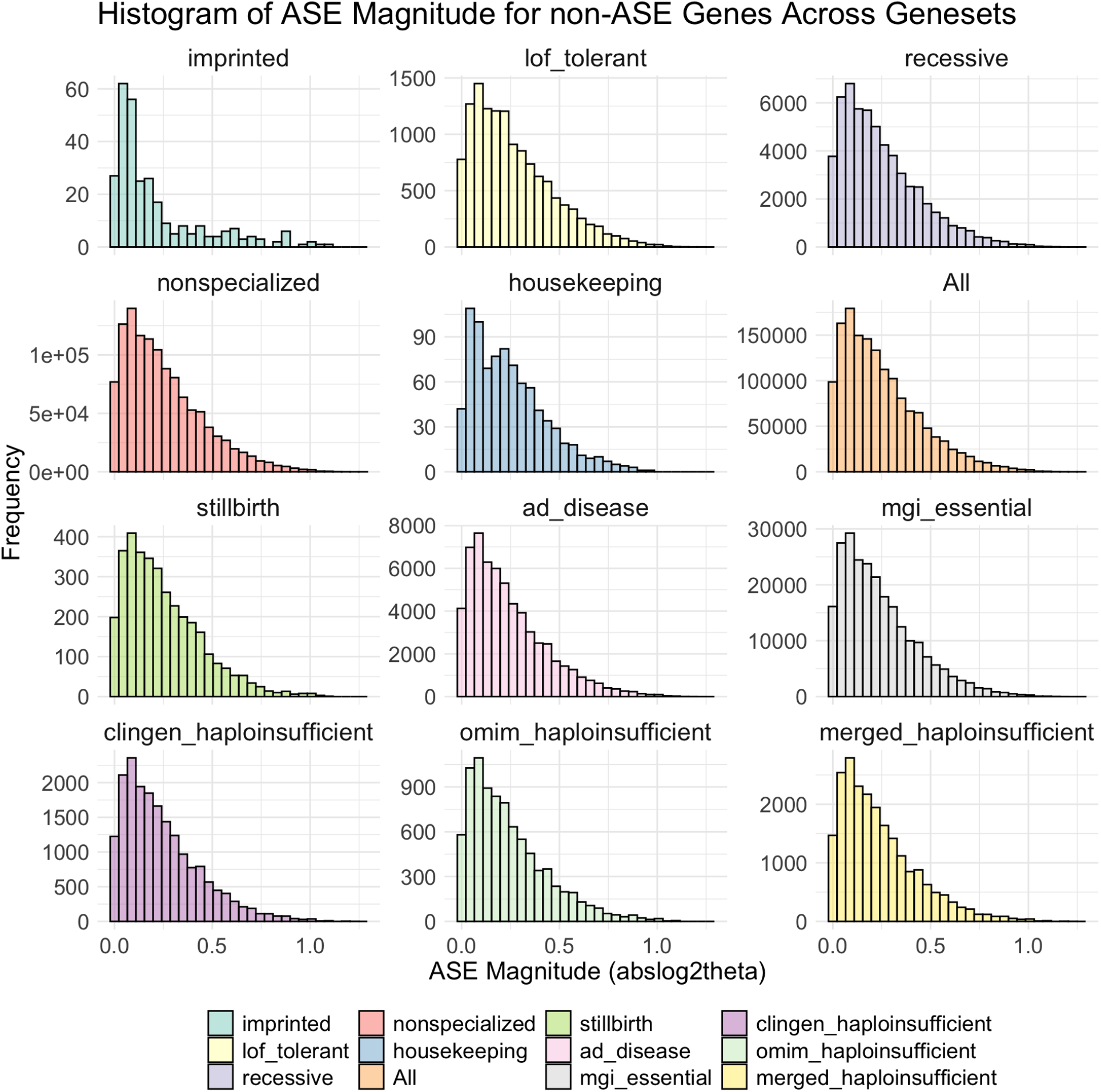
ASE magnitude histograms in non-ASE genes from different genesets.

**Figure S15.**
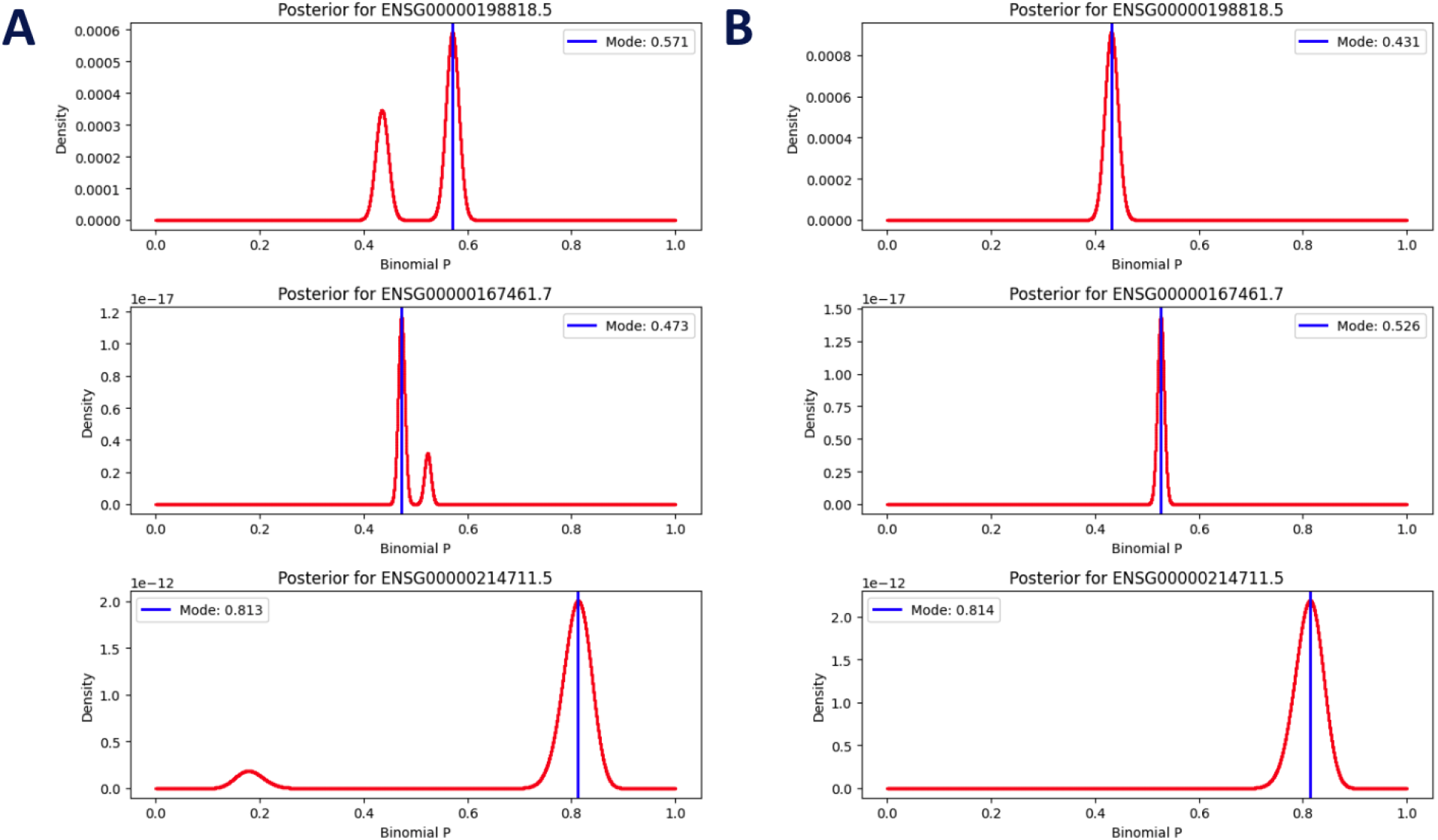
**Fixing first site VS fixing highest coverage site in QuickBeast on three genes from NA12878 sample**. **(A) qb with fixing the first site (multiple local maximum)** **(B) qb with fixing the highest coverage site (single maxima)**

**Figure S16.**
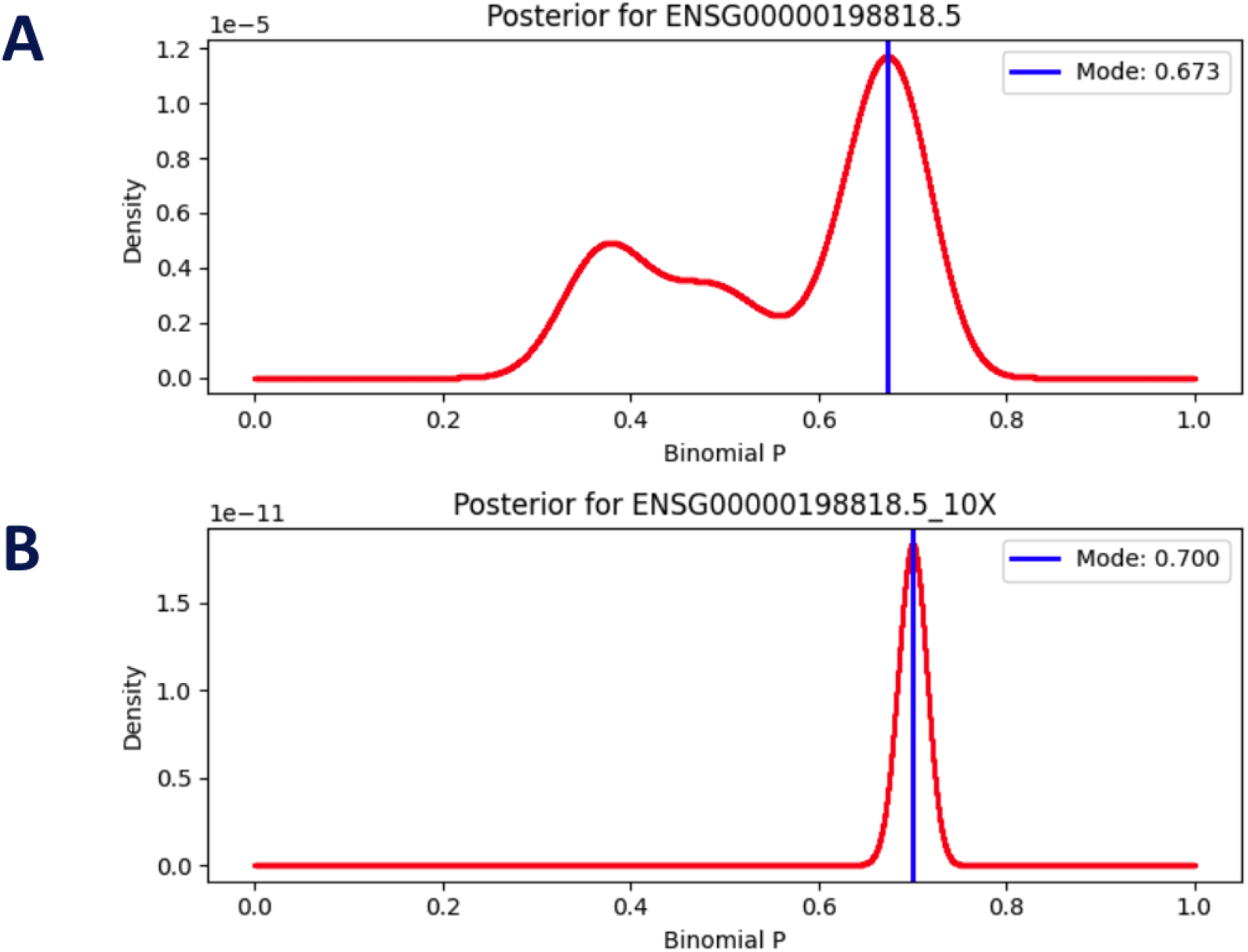
Trimodal distribution changes to Unimodel distribution after 10X coverage increases from NA12878 sample data of gene ENSG00000198818.5.

**Figure S17.**
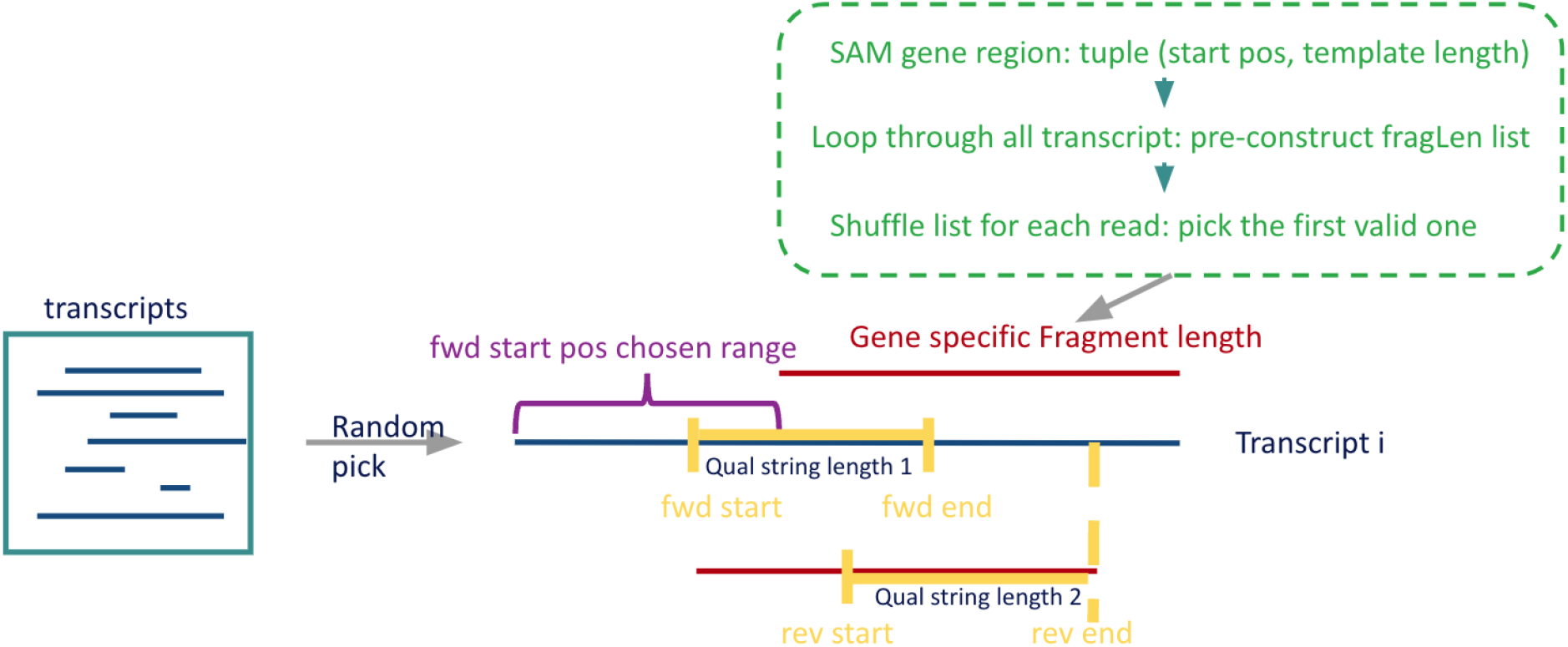
Diagram for unbiased splice reads simulator.

**Figure S18:**
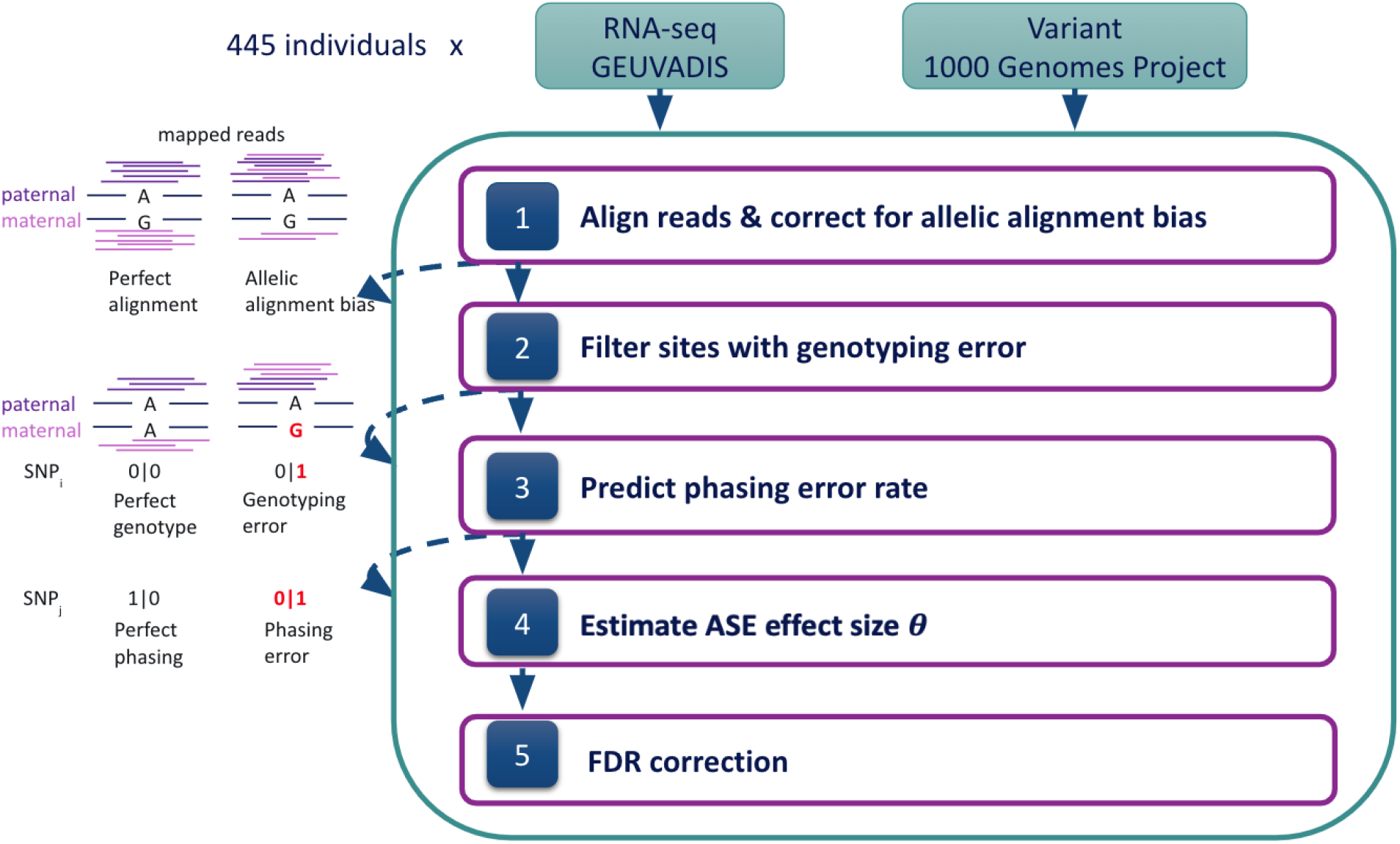
pipeline overview. It contains all major steps processing the steps that are explained in Methods.

**Figure S19:**
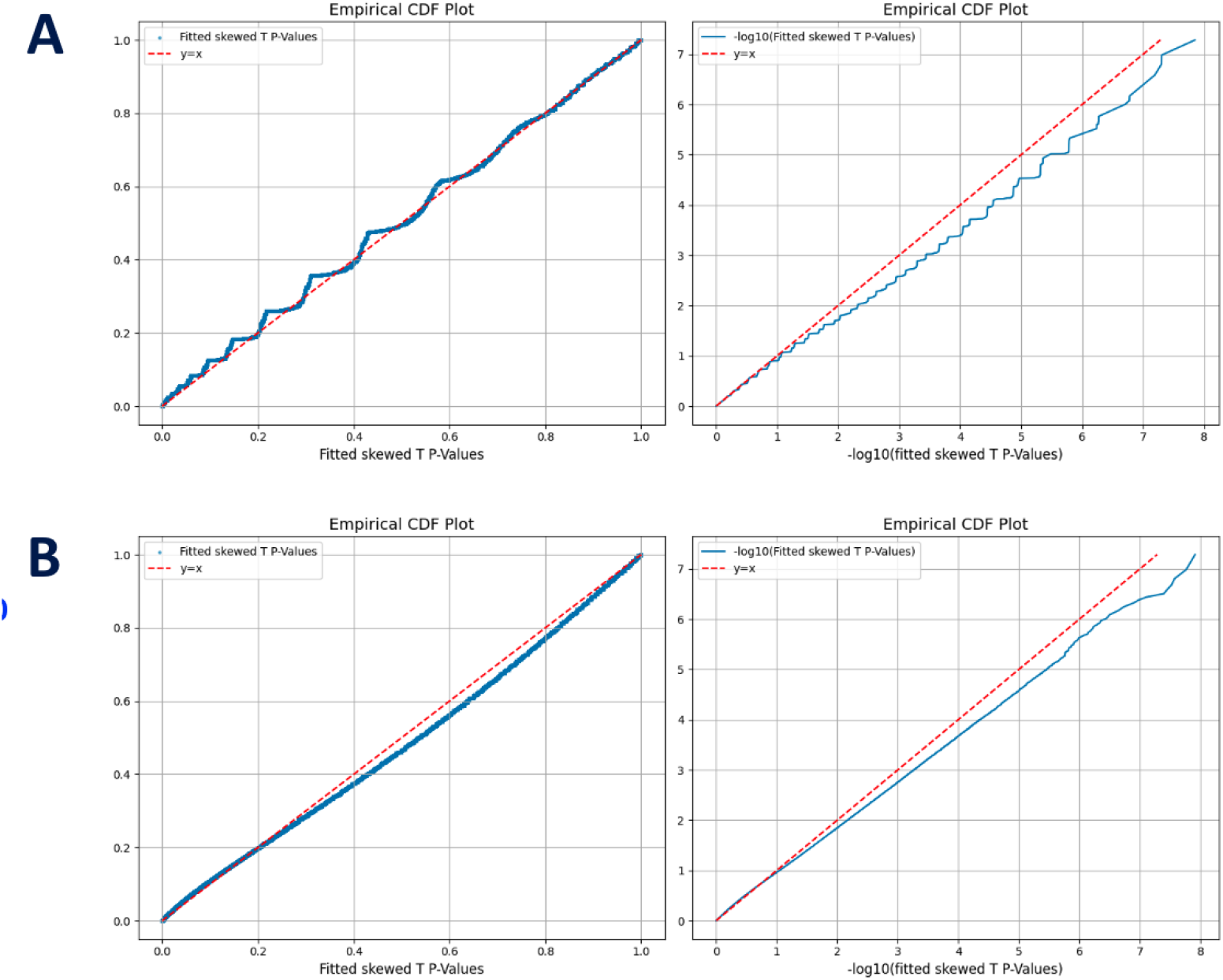
**Empirical CDF using p values from 19M null simulations** (calculation details in (**Supplementary Text 4.1 (2)**) **fit to skewed t distribution** **A: ECDF of fitted skewed t distribution p values and -log10 (p values) from 19207904 null stimulated genes with 3 hets and 30 read depth per het and theta = 1** **B: ECDF of fitted skewed t distribution p values and -log10 (p values) from 19207904 null stimulated genes with 10 hets and 100 read depth per het and theta = 1**

**Figure S20:**
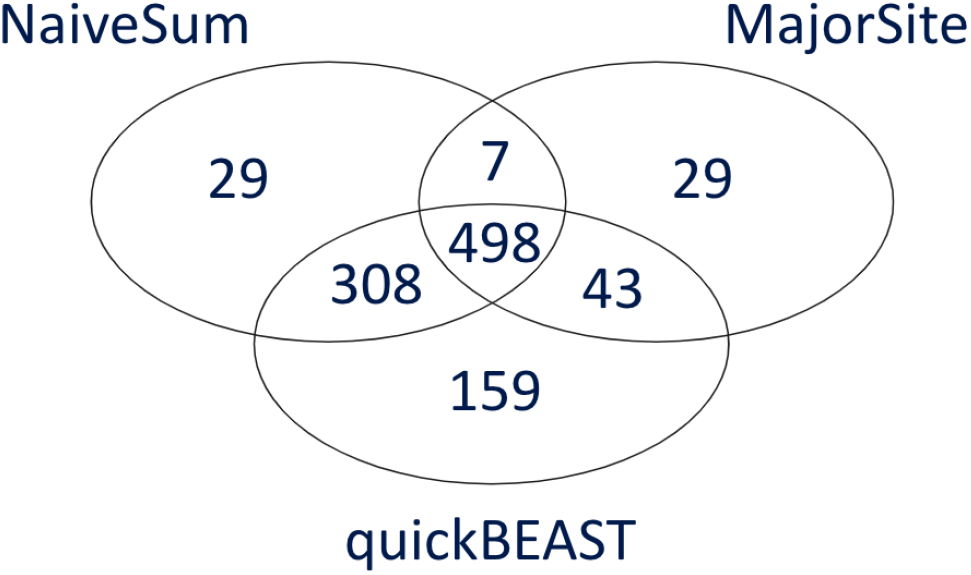
ASE results and comparative analysis on NA12878. Comparison with NaiveSum and MajorSite. Concordance of ASE genes detection is shown between BEASTIE, NaiveSum and MajorSite. The version of BEASTIE used here is the latest version called quickBEAST.

**Figure S21.**
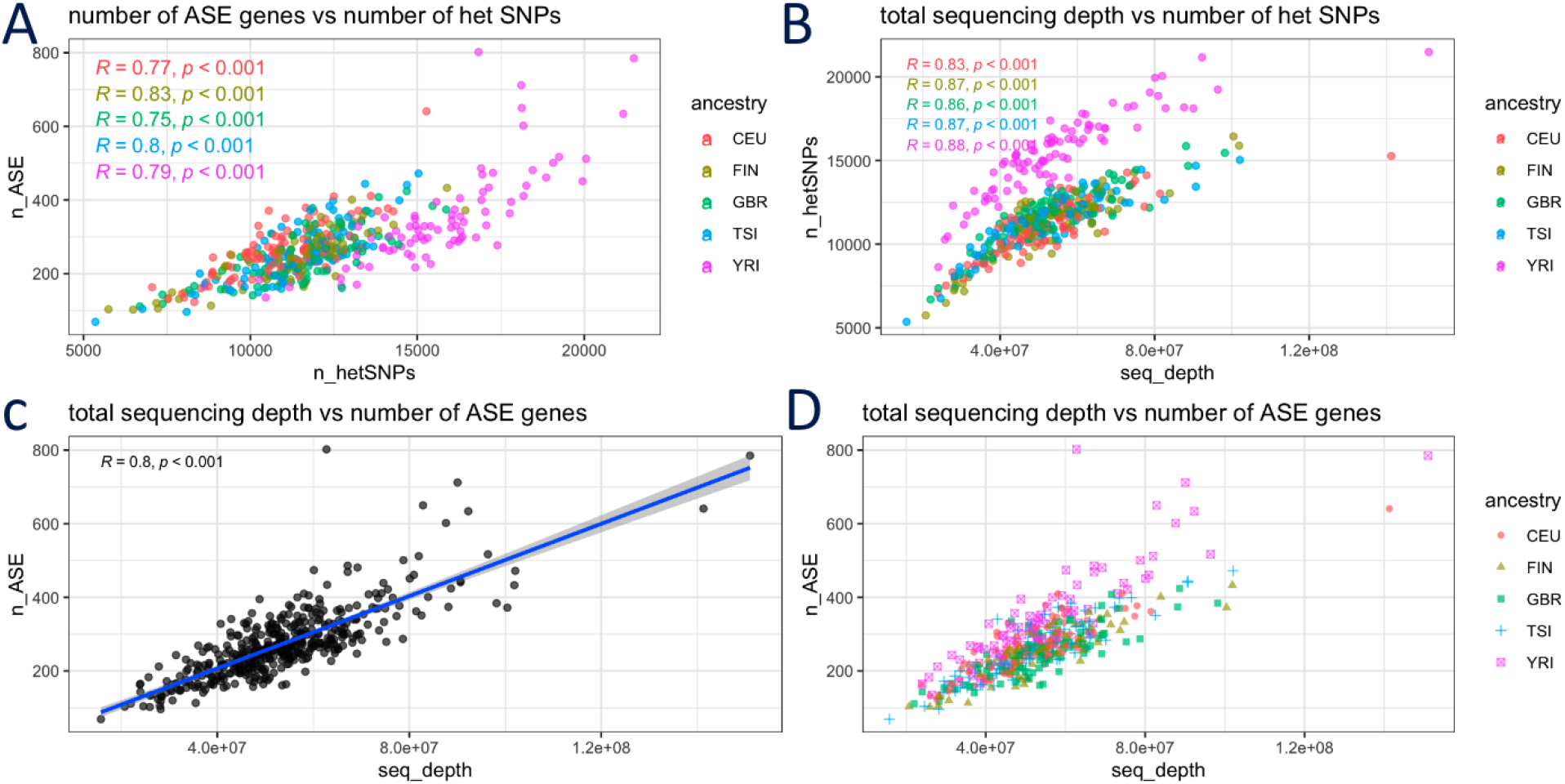
Positive pearson correlation between the total number of reads and ASE genes. (A)Scatterplot of the number of total heterozygous SNPs versus the number of ASE genes per individual. (B)Scatterplot of the total number of reads from aligned BAM files versus the number of heterozygous sites per individual. (C)Scatterplot of the total number of reads from aligned BAM files versus the number of ASE genes per individual. All panels: legend indicates the color/shape for each ancestry. Pearson correlation coefficient and p-value are labeled in the plot.

**Figure S22:**
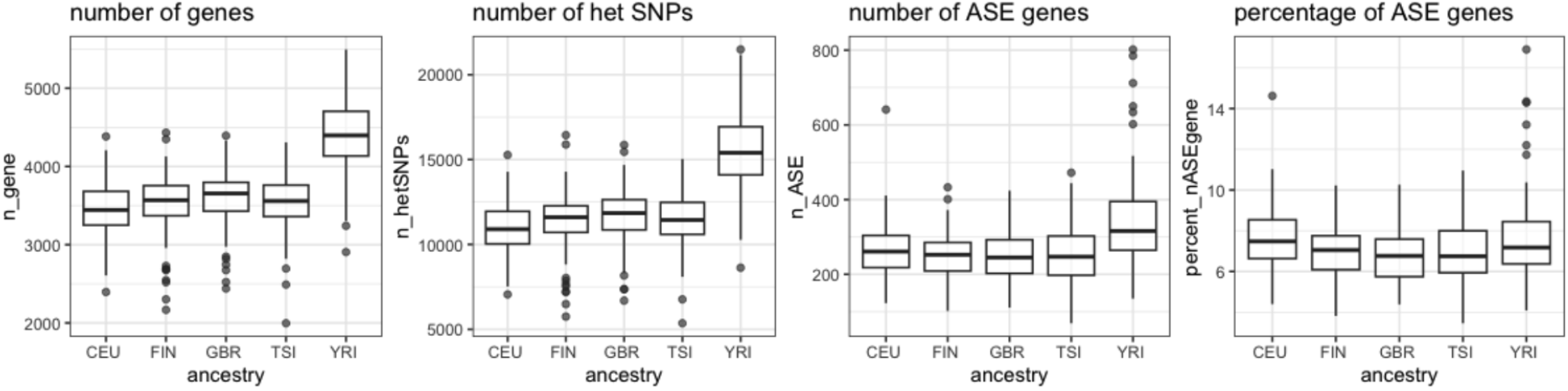
ancestry characteristics. Boxplot of number of genes, number of het SNPs, number of ASE genes, and percentage of ASE genes.

**Figure S23.**
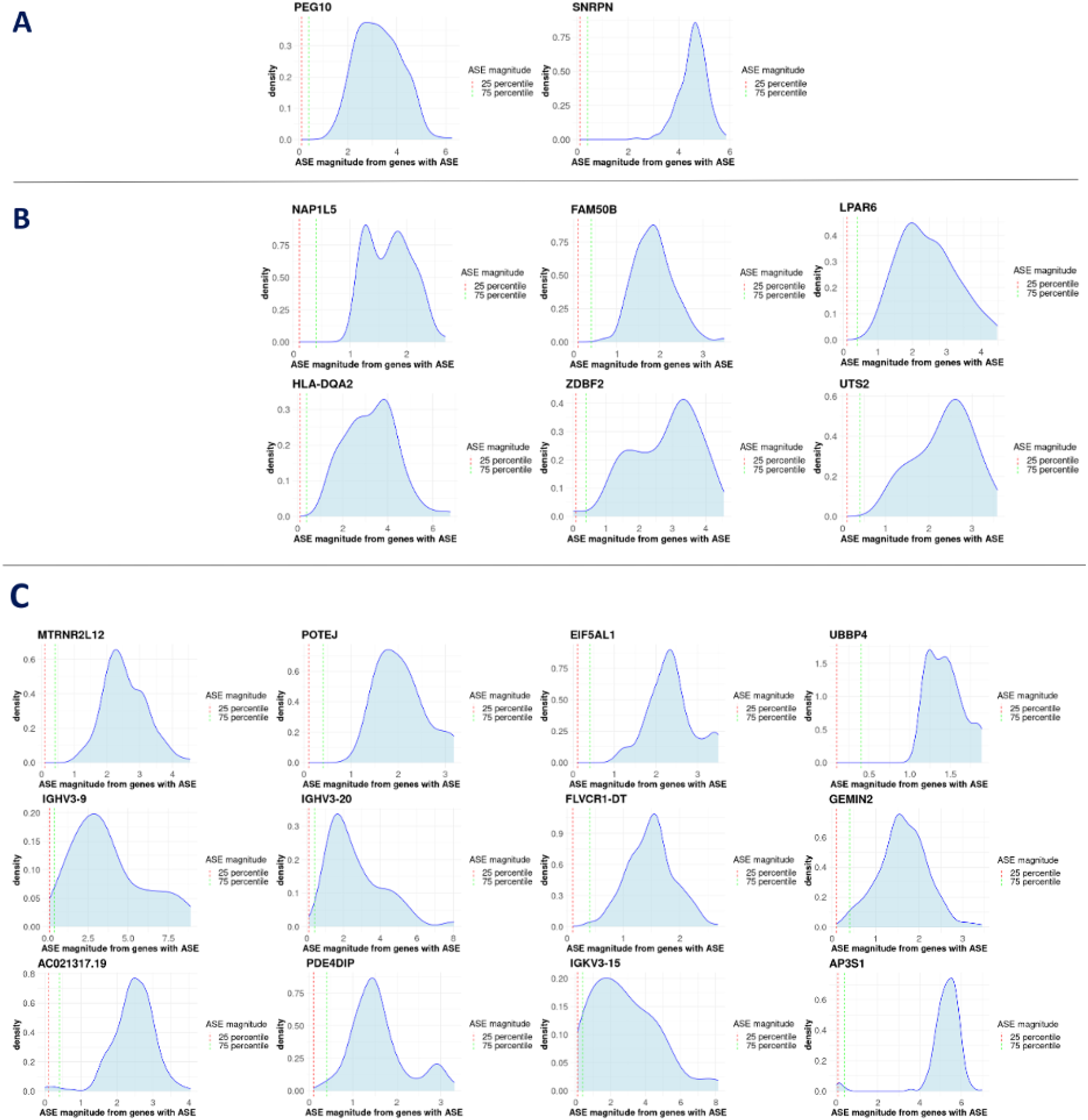
**Density plot of ASE magnitude for (A) 2 known imprinted genes, (B) 6 previously reported genes with imprinting status in specific tissues, (C) 17 genes with extreme ASE magnitude without prior literature on imprinting status**.

## References

1. Ammar J Alsheikh, Sabrina Wollenhaupt, Emily A King, Jonas Reeb, Sujana Ghosh, Lindsay R Stolzenburg, Saleh Tamim, Jozef Lazar, J Wade Davis, and Howard J Jacob. The landscape of gwas validation; systematic review identifying 309 validated non-coding variants across 130 human diseases. BMC medical genomics, 15(1):74, 2022.

2. Alexander Gusev, S Hong Lee, Gosia Trynka, Hilary Finucane, Bjarni J Vilhjálmsson, Han Xu, Chongzhi Zang, Stephan Ripke, Brendan Bulik-Sullivan, Eli Stahl, et al. Partitioning heritability of regulatory and cell-type-specific variants across 11 common diseases. The American Journal of Human Genetics, 95(5):535–552, 2014.

3. Joanna S Amberger, Carol A Bocchini, François Schiettecatte, Alan F Scott, and Ada Hamosh. Omim. org: Online mendelian inheritance in man (omim®), an online catalog of human genes and genetic disorders. Nucleic acids research, 43(D1):D789–D798, 2015.

4. Laure Frésard, Craig Smail, Nicole M Ferraro, Nicole A Teran, Xin Li, Kevin S Smith, Devon Bonner, Kristin D Kernohan, Shruti Marwaha, Zachary Zappala, et al. Identification of rare-disease genes using blood transcriptome sequencing and large control cohorts. Nature medicine, 25(6):911–919, 2019.

5. Kym M Boycott, Ana Rath, Jessica X Chong, Taila Hartley, Fowzan S Alkuraya, Gareth Baynam, Anthony J Brookes, Michael Brudno, Angel Carracedo, Johan T den Dunnen, et al. International cooperation to enable the diagnosis of all rare genetic diseases. The American Journal of Human Genetics, 100(5):695–705, 2017.

6. Fatemeh Peymani, Aiman Farzeen, and Holger Prokisch. Rna sequencing role and application in clinical diagnostic. Pediatric Investigation, 6(01):29–35, 2022.

7. Beryl B Cummings, Jamie L Marshall, Taru Tukiainen, Monkol Lek, Sandra Donkervoort, A Reghan Foley, Veronique Bolduc, Leigh B Waddell, Sarah A Sandaradura, Gina L O’Grady, et al. Improving genetic diagnosis in mendelian disease with transcriptome sequencing. Science translational medicine, 9(386):eaal5209, 2017.

8. Laura S Kremer, Daniel M Bader, Christian Mertes, Robert Kopajtich, Garwin Pichler, Arcangela Iuso, Tobias B Haack, Elisabeth Graf, Thomas Schwarzmayr, Caterina Terrile, et al. Genetic diagnosis of mendelian disorders via rna sequencing. Nature communications, 8(1):15824, 2017.

9. Xin Li, Yungil Kim, Emily K Tsang, Joe R Davis, Farhan N Damani, Colby Chiang, Gaelen T Hess, Zachary Zappala, Benjamin J Strober, Alexandra J Scott, et al. The impact of rare variation on gene expression across tissues. Nature, 550(7675):239–243, 2017.

10. Graham A Heap, Jennie HM Yang, Kate Downes, Barry C Healy, Karen A Hunt, Nicholas Bockett, Lude Franke, Patrick C Dubois, Charles A Mein, Richard J Dobson, et al. Genome-wide analysis of allelic expression imbalance in human primary cells by high-throughput transcriptome resequencing. Human molecular genetics, 19(1):122–134, 2010.

11. Pablo Llavona, Michele Pinelli, Margherita Mutarelli, Veer Singh Marwah, Simone Schimpf-Linzenbold, Sebastian Thaler, Efdal Yoeruek, Jan Vetter, Susanne Kohl, and Bernd Wissinger. Allelic expression imbalance in the human retinal transcriptome and potential impact on inherited retinal diseases. Genes, 8(10):283, 2017.

12. Monika Stachowiak, Izabela Szczerbal, and Krzysztof Flisikowski. Investigation of allele-specific expression of genes involved in adipogenesis and lipid metabolism suggests complex regulatory mechanisms of ppargc1a expression in porcine fat tissues. BMC genetics, 19:1–9, 2018.

13. Oleg Mayba, Houston N Gilbert, Jinfeng Liu, Peter M Haverty, Suchit Jhunjhunwala, Zhaoshi Jiang, Colin Watanabe, and Zemin Zhang. Mbased: allele-specific expression detection in cancer tissues and cell lines. Genome biology, 15:1–21, 2014.

14. Daniel A Skelly, Marnie Johansson, Jennifer Madeoy, Jon Wakefield, and Joshua M Akey. A powerful and flexible statistical framework for testing hypotheses of allele-specific gene expression from rna-seq data. Genome research, 21(10):1728–1737, 2011.

15. Jiaxin Fan, Jian Hu, Chenyi Xue, Hanrui Zhang, Katalin Susztak, Muredach P Reilly, Rui Xiao, and Mingyao Li. Asep: Gene-based detection of allele-specific expression across individuals in a population by rna sequencing. PLoS genetics, 16(5):e1008786, 2020.

16. Stephane E Castel, Pejman Mohammadi, Wendy K Chung, Yufeng Shen, and Tuuli Lappalainen. Rare variant phasing and haplotypic expression from rna sequencing with phaser. Nature communications, 7(1):12817, 2016.

17. Jared O’Connell, Deepti Gurdasani, Olivier Delaneau, Nicola Pirastu, Sheila Ulivi, Massimiliano Cocca, Michela Traglia, Jie Huang, Jennifer E Huffman, Igor Rudan, et al. A general approach for haplotype phasing across the full spectrum of relatedness. PLoS genetics, 10(4):e1004234, 2014.

18. Manuel A Rivas, Matti Pirinen, Donald F Conrad, Monkol Lek, Emily K Tsang, Konrad J Karczewski, Julian B Maller, Kimberly R Kukurba, David S DeLuca, Menachem Fromer, et al. Effect of predicted protein-truncating genetic variants on the human transcriptome. Science, 348(6235):666–669, 2015.

19. Stephane E Castel, Ami Levy-Moonshine, Pejman Mohammadi, Eric Banks, and Tuuli Lappalainen. Tools and best practices for data processing in allelic expression analysis. Genome biology, 16:1–12, 2015.

20. Alessandro Romanel, Sara Lago, Davide Prandi, Andrea Sboner, and Francesca Demichelis. Aseq: fast allele-specific studies from next-generation sequencing data. BMC medical genomics, 8:1–12, 2015.

21. Benjamin Deonovic, Yunhao Wang, Jason Weirather, Xiu-Jie Wang, and Kin Fai Au. Idp-ase: haplotyping and quantifying allele-specific expression at the gene and gene isoform level by hybrid sequencing. Nucleic acids research, 45(5):e32–e32, 2017.

22. Chris T Harvey, Gregory A Moyerbrailean, Gordon O Davis, Xiaoquan Wen, Francesca Luca, and Roger Pique-Regi. Quasar: quantitative allele-specific analysis of reads. Bioinformatics, 31(8):1235–1242, 2015.

23. Kimberly R Kukurba, Rui Zhang, Xin Li, Kevin S Smith, David A Knowles, Meng How Tan, Robert Piskol, Monkol Lek, Michael Snyder, Daniel G MacArthur, et al. Allelic expression of deleterious protein-coding variants across human tissues. PLoS genetics, 10(5):e1004304, 2014.

24. David A Knowles, Joe R Davis, Hilary Edgington, Anil Raj, Marie-Julie Favé, Xiaowei Zhu, James B Potash, Myrna M Weissman, Jianxin Shi, Douglas F Levinson, et al. Allele-specific expression reveals interactions between genetic variation and environment. Nature methods, 14(7):699–702, 2017.

25. Kimberly R Kukurba and Stephen B Montgomery. Rna sequencing and analysis. Cold Spring Harbor Protocols, 2015(11):pdb–top084970, 2015.

26. Jacob F Degner, John C Marioni, Athma A Pai, Joseph K Pickrell, Everlyne Nkadori, Yoav Gilad, and Jonathan K Pritchard. Effect of read-mapping biases on detecting allele-specific expression from rna-sequencing data. Bioinformatics, 25(24):3207–3212, 2009.

27. Tuuli Lappalainen, Michael Sammeth, Marc R Friedländer, Peter AC ‘t Hoen Jean Monlong, Manuel A Rivas, Mar Gonzalez-Porta, Natalja Kurbatova, Thasso Griebel, Pedro G Ferreira, et al. Transcriptome and genome sequencing uncovers functional variation in humans. Nature, 501(7468):506–511, 2013.

28. Alan Hodgkinson, Jean-Christophe Grenier, Elias Gbeha, and Philip Awadalla. A haplotype-based normalization technique for the analysis and detection of allele specific expression. BMC bioinformatics, 17:1–10, 2016.

29. Erik Garrison, Jouni Sirén, Adam M Novak, Glenn Hickey, Jordan M Eizenga, Eric T Dawson, William Jones, Shilpa Garg, Charles Markello, Michael F Lin, et al. Variation graph toolkit improves read mapping by representing genetic variation in the reference. Nature biotechnology, 36(9):875–879, 2018.

30. Bryce Van De Geijn, Graham McVicker, Yoav Gilad, and Jonathan K Pritchard. Wasp: allele-specific software for robust molecular quantitative trait locus discovery. Nature methods, 12(11):1061–1063, 2015.

31. Jing Wang Pedro Alves Debasish Raha Arif Harmanci Jing Leng Robert Bjornson Yong Kong Naoki Kitabayashi Nitin Bhardwaj Mark Rubin Michael Snyder Joel Rozowsky, Alexej Abyzov and Mark Gerstein. Alleleseq: analysis of allele-specific expression and binding in a network framework. Molecular Systems Biology, 7:522, 2011.

32. Lei Tian, Asifullah Khan, Zhilin Ning, Kai Yuan, Chao Zhang, Haiyi Lou, Yuan Yuan, and Shuhua Xu. Genome-wide comparison of allele-specific gene expression between african and european populations. Human molecular genetics, 27(6):1067–1077, 2018.

33. Olivier Delaneau and Jonathan Marchini. Integrating sequence and array data to create an improved 1000 genomes project haplotype reference panel. Nature communications, 5(1):3934, 2014.

34. Jennifer M Frost, Dave Monk, Taita Stojilkovic-Mikic, Kathryn Woodfine, Lyn S Chitty, Adele Murrell, Philip Stanier, and Gudrun E Moore. Evaluation of allelic expression of imprinted genes in adult human blood. PLoS One, 5(10):e13556, 2010.

35. Simone Rizzetto, David NP Koppstein, Jerome Samir, Mandeep Singh, Joanne H Reed, Curtis H Cai, Andrew R Lloyd, Auda A Eltahla, Christopher C Goodnow, and Fabio Luciani. B-cell receptor reconstruction from single-cell rna-seq with vdjpuzzle. Bioinformatics, 34(16):2846–2847, 2018.

36. William McLaren, Laurent Gil, Sarah E Hunt, Harpreet Singh Riat, Graham RS Ritchie, Anja Thormann, Paul Flicek, and Fiona Cunningham. The ensembl variant effect predictor. Genome biology, 17:1–14, 2016.

## Bibliography

[1] J. F. Degner et al., “Effect of read-mapping biases on detecting allele-specific expression from RNA-sequencing data,” Bioinformatics, vol. 25, no. 24, pp. 3207–3212, Dec. 2009.

[2] T. Lappalainen et al., “Transcriptome and genome sequencing uncovers functional variation in humans,” Nature, vol. 501, no. 7468, pp. 506–511, Sep. 2013.

[3] K. R. Kukurba et al., “Allelic expression of deleterious protein-coding variants across human tissues,” PLoS Genet., vol. 10, no. 5, p. e1004304, May 2014.

[4] A. Romanel, S. Lago, D. Prandi, A. Sboner, and F. Demichelis, “ASEQ: fast allele-specific studies from next-generation sequencing data,” BMC Med. Genomics, vol. 8, p. 9, Mar. 2015.

[5] J. Rozowsky et al., “AlleleSeq: analysis of allele-specific expression and binding in a network framework,” Mol. Syst. Biol., vol. 7, p. 522, Aug. 2011.

[6] S. E. Castel, A. Levy-Moonshine, P. Mohammadi, E. Banks, and T. Lappalainen, “Tools and best practices for data processing in allelic expression analysis,” Genome Biol., vol. 16, no. 1, p. 195, Sep. 2015.

[7] S. E. Castel, P. Mohammadi, W. K. Chung, Y. Shen, and T. Lappalainen, “Rare variant phasing and haplotypic expression from RNA sequencing with phASER,” Nat. Commun., vol. 7, p. 12817, Sep. 2016.

[8] O. Mayba et al., “MBASED: allele-specific expression detection in cancer tissues and cell lines,” Genome Biol., vol. 15, no. 8, p. 405, Aug. 2014.

[9] D. A. Skelly, M. Johansson, J. Madeoy, J. Wakefield, and J. M. Akey, “A powerful and flexible statistical framework for testing hypotheses of allele-specific gene expression from RNA-seq data,” Genome Res., vol. 21, no. 10, pp. 1728–1737, Oct. 2011.

[10] C. T. Harvey, G. A. Moyerbrailean, G. O. Davis, X. Wen, F. Luca, and R. Pique-Regi, “QuASAR: quantitative allele-specific analysis of reads,” Bioinformatics, vol. 31, no. 8, pp. 1235–1242, Apr. 2015.

[11] X. Wei and X. Wang, “A computational workflow to identify allele-specific expression and epigenetic modification in maize,” Genomics Proteomics Bioinformatics, vol. 11, no. 4, pp. 247–252, Aug. 2013.

[12] J. M. Frost et al., “Evaluation of allelic expression of imprinted genes in adult human blood,” PLoS One, vol. 5, no. 10, p. e13556, Oct. 2010.

[13] L. Morcos et al., “Genome-wide assessment of imprinted expression in human cells,” Genome Biol., vol. 12, no. 3, p. R25, Mar. 2011.

[14] M. Lek et al., “Analysis of protein-coding genetic variation in 60,706 humans,” Nature, vol. 536, no. 7616, pp. 285–291, Aug. 2016.

[15] S. Petrovski, Q. Wang, E. L. Heinzen, A. S. Allen, and D. B. Goldstein, “Genic intolerance to functional variation and the interpretation of personal genomes,” PLoS Genet., vol. 9, no. 8, p. e1003709, Aug. 2013.

[16] M. Bernasconi et al., “Quantitative profiling of housekeeping and Epstein-Barr virus gene transcription in Burkitt lymphoma cell lines using an oligonucleotide microarray,” Virol. J., vol. 3, p. 43, Jun. 2006.

[17] H. L. Rehm et al., “ClinGen--the Clinical Genome Resource,” N. Engl. J. Med., vol. 372, no. 23, pp. 2235–2242, Jun. 2015.

[18] B. Georgi, B. F. Voight, and M. Bućan, “From mouse to human: evolutionary genomics analysis of human orthologs of essential genes,” PLoS Genet., vol. 9, no. 5, p. e1003484, May 2013.

[19] T. J. Hayeck et al., “Improved Pathogenic Variant Localization via a Hierarchical Model of Sub-regional Intolerance,” Am. J. Hum. Genet., vol. 104, no. 2, pp. 299–309, Feb. 2019.

[20] K. E. Stanley et al., “Causal Genetic Variants in Stillbirth,” N. Engl. J. Med., vol. 383, no. 12, pp. 1107–1116, Sep. 2020.

[21] I. Zaitoun and H. Khatib, “Assessment of genomic imprinting of SLC38A4, NNAT, NAP1L5, and H19 in cattle,” BMC Genet., vol. 7, p. 49, Oct. 2006.

[22] A. Zhang et al., “Novel retrotransposed imprinted locus identified at human 6p25,” Nucleic Acids Res., vol. 39, no. 13, pp. 5388–5400, Jul. 2011.

[23] Y. Baran et al., “The landscape of genomic imprinting across diverse adult human tissues,” Genome Res., vol. 25, no. 7, pp. 927–936, Jul. 2015.

[24] F. Voorthuijzen, C. Stroobandt, W. Van Criekinge, T. Goovaerts, and T. De Meyer, “Loss-of-Imprinting of HM13 Leads to Poor Prognosis in Clear Cell Renal Cell Carcinoma,” Biomolecules, vol. 14, no. 8, p. 936, Aug. 2024.

[25] H. Kobayashi et al., “Identification of the mouse paternally expressed imprinted gene Zdbf2 on chromosome 1 and its imprinted human homolog ZDBF2 on chromosome 2,” Genomics, vol. 93, no. 5, pp. 461–472, May 2009.

[26] S. A. Carrion, J. J. Michal, and Z. Jiang, “Imprinted Genes: Genomic Conservation, Transcriptomic Dynamics and Phenomic Significance in Health and Diseases,” Int. J. Biol. Sci., vol. 19, no. 10, p. 3128, 2023.

[27] S. E. Hiby, A. King, A. Sharkey, and Y. W. Loke, “Molecular studies of trophoblast HLA-G: polymorphism, isoforms, imprinting and expression in preimplantation embryo,” Tissue Antigens, vol. 53, no. 1, pp. 1–13, Jan. 1999.

