## Supplementary Text for "Bayesian Estimation of Allele-Specific Expression in the Presence of Phasing Uncertainty"

#### Supplementary Text 1 Data Source

##### Supplementary Text 1.1 1000 Genome Project data

The RNA-seq fastq read and genomic data used in this study are publicly available. Processed VCFs were obtained from the 1000 Genome Project data Phase 3 data set, while raw RNA-seq fastq data were downloaded from the Geuvadis project's lymphoblastoid (LCL) cell line [1]. Our study population consisted of 445 individuals with complete data, including 87 Yoruba (YRI), 89 Utah (CEU), 86 British (GBR), 92 Finnish (FIN), and 91 Tuscan (TSI) individuals. Quality-controlled data included approximately 11,300 bi-allelic exonic heterozygous SNPs, covered by at least one RNA-seq read, distributed across approximately 5,000 autosomal genes. The number of SNPs and genes varied slightly due to differences in sequencing depth and ancestry. The median depth of the Geuvadis samples was 55 million mapped reads.

##### Supplementary Text 1.2 NA12878 sequence data and GIAB phased VCF

NA12878 is a female from Utah with CEU ancestry. We downloaded the latest version of the gold standard phased haplotype VCF file with GRCh37 coordinates from the National Institute of Standards and Technology (NIST) Genome-In-A-Bottle consortium FTP site [2], as well as the raw RNA-seq paired-end fastq reads [3] from LCL cell line on the Sequence Read Archive at NCBI. This sample is part of the CEPH collection of the Hapmap project [4][5]. In total, we obtained 33,449 exonic SNP pairs from 1,667,183 phase-resolved bi-allelic heterozygous sites on 22 autosomes from the GIAB VCF files, which were then phased using SHAPEIT2 for our analyses.

##### Supplementary Text 1.3 GM19440 sequence data and experimentally phased VCF

GM19440 is an AFR male from Webuye, Kenya. We obtained experimentally phased VCF files with haplotype information determined using the 10x Genomic platform in GRCh38 coordinates from Belsare et al. 2019 [6]. We downloaded the corresponding 1KGP VCF file for the GRCh38 assembly from the TGP website and used the GATK LiftoverVCF tool to convert the coordinates to GRCh37. After this conversion, we obtained a total of 65,831 exonic SNP pairs from 22 autosomes, out of the 2,813,858 phase resolved bi-allelic heterozygous sites in the VCF files. We then used SHAPEIT2 to phase the SNPs for our analyses.

##### Supplementary Text 1.4 1000 Genome Project trio dataset

We obtained a trio data set from the 1000 Genome Project data site, which consisted of the Yoruba family NA19240 (daughter), NA19238 (mother), and NA19239 (father). The data set was obtained from the pilot phase of the NCBI build 36.3 human genome reference sequence from March 2006 and DNA source of lymphoblastoid cell lines distributed by the Coriell Institute Deep Coverage WGS 2008. The SNP data for this trio were accessed from an online portal [7].

##### Supplementary Text 1.5 ASE quantification and output

quickBEAST employs subgrid algorithms to efficiently identify the mode of the posterior distribution, generating binomial p-values. Quantifications used in the study are listed below:

$$\theta = \frac{p}{1-p}$$

$$ASE\ magnitude = abs(log2(\theta))$$

$$ASE\ gene:\ p\ value\ under\ FDR\ correction < 0.05$$

high ASE value: ASE magnitude above 75% percentile (0.4034574)

low ASE value: ASE magnitude below 25% percentile (0.1020076)

#### Supplementary Text 1.6 Model Evaluation and Comparison

Models were compared based on receiver operating characteristics (ROC) curves. For ROC analyses we report the area under the ROC curve (AUC). To facilitate ROC analyses, we classify genes as positives (having ASE) or negatives (not having ASE). Positive cases were simulated by choosing a value of θ different from 1; negative cases were simulated by specifying θ equal to 1.

### Supplementary Text 2 Helper Models

#### Supplementary Text 2.1 BEASTIE STAN - Efficient Marginalization of Haplotype Phase with a Hidden Markov Model

Inference with the BEASTIE STAN model is performed via Markov Chain Monte Carlo (MCMC) as implemented in the package Stan [8]. Briefly, Stan uses a Hamiltonian Monte Carlo [9] to propose samples for the latent variables in the model (θ), and those samples are accepted with probability proportional to a ratio of likelihoods, priors, and proposal probabilities under the Hamiltonian. The likelihoods and priors of the BEASTIE model are given by:

$$\theta \sim \log_2\text{normal}(0, 0.4)$$

$$\phi \sim f(\pi, \phi^*)$$

$$X_i \sim \{binom(N_i, p) \; if \; \phi_i = 1, \; binom(N_i, 1 - p) \quad if \; \phi_i = 0$$

where p = θ / (θ + 1). The log2normal prior on θ was chosen after assessing the genome-wide distribution of ASE values across a single individual in the 1KGP [10], in which it was found that ASE values tend to vary between a halving and a doubling of gene expression compared to the other allele.

For the BEASTIE-latent model, which requires no estimates of switching error rate, a global switching error rate π is included in the model as a latent variable, and is given a beta(1,10) prior. This prior weakly favors low switching error rates, consistent with our finding that switching error rates tend to be less than 10% (Section 3.1). For this model, π is effectively integrated out by MCMC during sampling.

The vector ϕ = {ϕ_i : i = 1..n} denotes the true phasing of the n sites in the gene. The likelihood of ϕ, conditional on the predicted phasing * and the vector of site-specific phasing error rates , is evaluated by an inhomogenous hidden Markov model (HMM). Briefly, the HMM evaluates a likelihood function in which i=i* (i.e., the true phase of site i is the same as the predicted phase for that site) with probability ψ, where ψ is the probability that the number of switching errors prior to (i.e., 5' of) site i is even.

The likelihood of ϕ in the BEASTIE model is given by the function $f(\pi, \phi^*)$, which evaluates to the probability that the true phasing of heterozygous sites in the gene is ϕ, given the predicted phasing $\phi^*$ and the vector of expected switching error rates π. Because ϕ is discrete, and because the Hamiltonial Monte Carlo in Stan cannot be used with discrete latent variables, we need to integrate ϕ out of the model analytically within Stan. We now describe an efficient method for doing this, which is based on an inhomogenous (nonstationary) hidden Markov model.

We construct a simple HMM with two emitting states ("even" and "odd"), as depicted in **Figure 1B**. The emissions of the HMM consist of the alternate allele counts $X_i$ in the BEASTIE model. The start/stop state is silent (i.e., makes no emissions), and exists solely to delimit the beginning and end of the HMM's emission sequence. The even state represents having an even number of switching errors up to site *i*, while the odd state represents having an odd number of switching errors up to site *i*. Note that an even number of switching errors between the first site and site *i* results in site *i* being correctly phased, whereas an odd number of switching errors results in site *i* being incorrectly phased. This derives from the fact that each switching error reverses, or toggles, the phasing status of the current site. As such, to keep track of phasing status as we traverse the sites in the gene, we need only differentiate between an even number of switching errors and an odd number of switching errors up to the current site.

We model the occurrence of a switching error as a transition between the even state and the odd state (in either direction). In contrast, we model the occurrence of no switching error at a site as a self-transition from a state back to the same state. Thus, transitions between the even and odd states incur a probability of $\pi_i$ (switching error), while self-transitions incur $1 - \pi_i$ (no switching error). Transitions into the stop state incur probability $\varepsilon$, which is set to an arbitrary, small value, since it has a negligible effect on inference when the number of sites is fixed.

Emission probabilities are computed using a binomial distribution. Within the even state, since there have been an even number of switching errors, we take the predicted phasing $\phi_i^*$ as being the correct phase of site *i*, since an even number of switching errors results in a correct absolute phase of site *i*. Thus, if $\phi_i^* = 1$, the maternal chromosome contains the alternate allele, and we assess an emission probability given by binomial($X_i$ | $N_i$, *p*), where *p* is computed from the current estimate of $\theta$ as described in **section 2.1**. If instead $\phi_i^* = 0$, the maternal chromosome contains the reference allele, and we assess an emission probability of binomial($X_i$ | $N_i$, 1-*p*). For the odd state, since there have been an odd number of switching errors, the predicted phase is incorrect, and we need to swap the alleles when computing likelihoods. Thus, if $\phi_i^* = 1$, the emission probability is binomial($X_i$ | $N_i$, 1-*p*), and if $\phi_i^* = 0$, the emission probability is binomial($X_i$ | $N_i$, *p*).

We use the standard *forward algorithm* (Durbin *et al.*, 1998; Rabiner *et al.*, 1989) to efficiently sum over all possible state paths through the HMM_i.e., all possible phasings of the *n* sites. This effectively marginalizes out $\phi_i$, producing a new model in which the likelihood of $X_i$ is directly conditioned on $\pi_i$ and $\phi_i^*$. It can be shown by induction that this marginalization computed via the forward algorithm is exact and gives the desired likelihood. Because the number of emitting states in the HMM is fixed at 2, the time complexity of this procedure is linear in the number of sites in the gene. This is a substantial improvement over the naive approach of enumerating all possible phasings, which would incur O($2^n$). Because this process is performed for every MCMC proposal in the BEASTIE model, the efficiency of this process is crucial.

#### Supplementary Text 2.2 QuickBEAST - C++ implementation of ASE Effect Size Estimation

QuickBEAST is a highly optimized implementation that employs the same statistical model as BEASTIE. Engineered in C++ for speed, QuickBEAST is designed to be applied quickly to large numbers of null simulations, enabling the calculation of empirical p-values. These empirical p-values enable QuickBEAST to control the false discovery rate (FDR) using standard methods such as Benjamini-Hockberg [11], and facilitate the use of QuickBEAST in genome-wide applications where controlling for multiple hypothesis testing is generally required. Coding details could be found in https://github.com/x811zou/QuickBEAST.

*Subgrid Algorithm for Mode Estimation.* QuickBEAST employs a recursive subgrid refinement to estimate the MAP effect size, particularly in scenarios featuring complex likelihood surfaces with single or multiple local maxima (**Supplementary Figure S15 & S16**). This algorithm operates by systematically narrowing the parameter space to identify the MAP efficiently.

Detailed Process of the Subgrid Algorithm:

1. Initial Grid Formation: The algorithm initiates by overlaying a grid across the entire parameter space.
2. Maximum finding: For each grid point, the algorithm samples the function to maximize

3. Iterative Refinement: Based on the computed values, the algorithm identifies the region of the grid with the highest likelihood values and subdivides it for a more granular examination. As the refinement progresses, the algorithm converges to the effect size with the highest posterior probability.

*Null Simulation and P-value Calculation*. QuickBEAST conducts simulations to construct a null distribution of the binomial proportion $p = \theta / (\theta + 1)$, in order to estimate an empirical p-value of the observed data. The process involves:

1. Null Simulation Execution: For each gene, QuickBEAST performs a large number of null simulations that mimic the specific features of the test gene, such as the number of heterozygous sites and total read depth, under the null hypothesis that $\theta=1$ (no ASE).
2. Calculation of Null Statistics: The statistic $p = \theta / (\theta + 1)$ is calculated both on the test gene and on the null simulated genes.
3. Distribution Fitting: The distribution of null statistics is then fitted to a skewed-*t* distribution. This fitting process captures the characteristics of the null distribution.
4. P-value Calculation: To determine the significance of the observed data, the statistic obtained from the sample gene is compared against the skewed-t distribution derived from the null simulations. An empirical p-value is estimated based on this comparison, by determining the proportion of nulls as extreme as the observed statistic; these p-values can be FDR-corrected by standard methods.

#### Supplementary Text 2.3 SELR (Switching Error Logistic Regressor) - Switch error as a function of LD, log10(inter SNP distance), and minor allele frequency

To assess the shapeit2 switching error rate between exonic heterozygous sites i and i+1 in the NA12878 and GM19440 samples, we compared the phasing results of each sample to their respective reference phasing. For NA12878, we compared the SHAPEIT2 phasing to the gold standard VCF phasing. In contrast, for GM19440, we used the experimental VCF as the reference phasing. A switching error occurs when SHAPEIT2 incorrectly infers the phase relationship between adjacent heterozygous sites, leading to an 'anti-phase' configuration where alleles that should be on the same chromosome (in phase) are incorrectly placed on opposite chromosomes. We counted any 'anti-phase' haplotype phasing mismatch within the same gene as an error, indicating a phasing error by SHAPEIT2. This method allowed us to quantify the accuracy of SHAPEIT2 in preserving the correct chromosomal alignment of alleles across exonic regions.

Switching Error Logistic Regressor (SELR) was calibrated on data from the NA12878 sample, evenly split into training and testing datasets (50% each), consisting of 17,549 heterozygous SNP pairs annotated with ancestry-specific allele frequencies from the 1000 Genomes Project and LD metrics. Evaluating the model's efficacy, we trained it on 8,729 site pairs and validated it on an independent set of 8,730 site pairs, each set comprising approximately 3.8% incorrectly phased site pairs. This methodology facilitated the calculation of switching error probabilities for each SNP pair.

To estimate the predictors that influence phasing accuracy, we used four measures: D', r2, γ1, and γ2. D' and r2 are measures of linkage disequilibrium that provide information about the tendency of pairs of alleles at two sites to be inherited together due to local recombination rates. γ1 and γ2 represent the absolute difference and the minimum of the minor allele frequencies of sites i and i+1, respectively. We also considered the log10 transformed unspliced genomic distance between sites i and i+1, denoted as d.

D' : normalized D
$r^2$ : correlation between loci
γ1 : abs difference($maf_i$, $maf_{i+1}$)
γ2 : minimum($maf_i$, $maf_{i+1}$)
d : log10 transformed unspliced genomic distance between sites i and i+1

where $maf_i$ is the minor allele frequency of site i, and r is given by:

$$r = \frac{D}{\sqrt{p_a(1-p_a)p_b(1-p_b)}}$$

for minor allele frequency $p_a$ at site i and $p_b$ at site i+1. D' and $r^2$ are both measures of linkage disequilibrium and provide information about the tendency of pairs of alleles at two sites to be inherited together, as a result of local recombination rates [12]. For the experiments based on Genome-in-a-Bottle (GIAB) data (**Supplementary Text 1.2**), D' and $r^2$ were computed for each consecutive SNP pair via R package LDlinkR [13].

The above four predictors estimated from GIAB were collected into vector Z, and the following logistic regression model was fit, separately, for each pair of sites i and i+1:

$$E\left(\pi_i | Z\right) = \frac{1}{1+e^{-Z\beta-\alpha}}$$

Given the values of β (coefficients) and α (intercept) fit to the GIAB data, this logistic regression model can then be applied to new genomes by providing the four predictors in Z, to produce $E\left(\pi_i | Z\right)$, the expected phasing error rate specific to the pair of sites i and i+1. This is repeated for each pair of sites in the gene, and the resulting values of $_i$ are provided to quickBEAST as fixed, point estimates. Coefficients from SELF are defined in **Supplementary Table S22**.

#### Supplementary Text 2.4 Parameterized Simulator

The parameterized simulator facilitates controlled simulations by allowing specific settings for multiple parameters (**Supplementary Table S23**), including (1) θ, denoting the degree of allelic skewing (ASE); (2) m, representing the count of heterozygous sites within a gene; (3) d, indicating the total read depth for each site; (4) π, the proportional rate of switching errors between consecutive het sites; (5) n, the total number of genes in the simulation. The simulation initiates by estimating the phasing $\boldsymbol{\phi}^*$ across all gene sites, achieved through a binomial distribution drawn with π as the parameter, thus modeling the occurrence of switching errors between consecutive site pairs. Each switching error has the potential to affect the phase of all subsequent (3') sites. Subsequent to this phasing step, the simulator generates allelic read counts, drawing again from a binomial distribution, with the site's total read count (sum of alternate and reference) and binomial proportion p = θ / (θ + 1) as parameters.

This parameterized approach is instrumental for thorough explorations across diverse parameter values. It offers a robust framework for constructing power curves and efficiently planning sequencing experiments, providing a structured yet flexible platform for genomic simulations.

#### Supplementary Text 2.5 Semi-empirical simulator

In accordance with **section 3.2.2**, we adopted a logistic regression model, termed the 'truth model', to estimate error probabilities from true phasing discrepancies, contrasting SHAPEIT2 phasing against the GIAB gold standard (denoted 'GIAB error probability'). We determined binary error labels (labeled 'GIAB error label') by comparing a uniformly distributed random probability U(0,1) against the GIAB error probability—assigning 1 for values less than or equal to the GIAB error probability and 0 otherwise. These labels served as the response variable in a subsequent logistic regression, the 'prediction model,' aimed at estimating error probabilities from predicted errors (labeled 'predicted error probability').

To generate arbitrary genes based on user-defined parameters (θ, m, n, d), we integrated features of both parametric and empirical simulators, conducting SNP sampling. Our empirical simulator leverages real gene data from the NA12878 sample phased by Genome-in-a-Bottle (GIAB), focusing on genes with at least one exonic heterozygous site as annotated by GENCODE version v19. Instead of simulating the number of heterozygous sites, this approach utilizes the actual count from the selected gene in NA12878. The high phasing accuracy of the GIAB NA12878 sample allows for the precise identification of SHAPEIT2-induced switching errors in consecutive exonic heterozygous sites. This data directly informs the simulation of allelic read counts, drawn from a binomial distribution with parameter $p = \theta / (\theta + 1)$, where θ represents the predefined allelic skewing (ASE) level, and the total read count is predefined. This methodology permits a nuanced exploration of the effects of θ and read counts on predictive accuracy. Inputs for the logistic regression predictors (D', $r^2$, MAF difference, and intersite distance) are calculated directly from the GIAB sample for each site pair and are used as inputs to the logistic regression model prior to running quickBEAST. Excluding θ and read counts, all model inputs derive directly from the real data phased by GIAB.

We compared three simulation approaches: 1) non-enrichment, where gene simulation with fixed parameters involved sampling a user-specified number (m) of SNPs from GIAB processed data for (n) genes, ensuring that the MAF of a current SNP was within the range defined by the previous SNP pair's minimum MAF and the sum of this minimum and the MAF difference between the SNP pair; 2) error enrichment, where we selectively retained simulated gene with an odd count of switching error within each gene.

#### Supplementary Text 2.6 Unbiased spliced reads simulator

To scrutinize mapping bias within our computational framework, we orchestrated a simulation deploying unbiased, uniformly distributed high-coverage fastq reads, each meticulously covering SNPs across both haplotypes as depicted in **Supplementary Figure S17** (https://github.com/x811zou/spliced_simulator). We aimed for gene-specific coverage, setting it at 100 to exceed the typical 75 read depth observed in empirical datasets from 1KGP. This simulation utilized observed base quality values from actual fastq reads mapped to gene-specific regions, to enhance the authenticity of each simulated read.

For every haplotype, the number of simulated reads was methodically determined: 50 reads multiplied by the gene's length, normalized by the length of the longest transcript. Base quality strings were randomly sampled from mapped reads corresponding to the gene region in the SAM file. Each read was then constructed by selecting a fragment sequence from a transcript, randomly picked from the gencode hg19 dataset.

Following their generation, simulated reads underwent alignment to the genome, employing identical software and parameters as those used for real data to ensure consistency. To pinpoint sites with notable mapping bias, we implemented a two-sided binomial test on the mapped read counts. Sites were marked significantly biased if their associated p-values fell below the 5% threshold.

#### Supplementary Text 2.7 Genotyping error model (GEM)

Sites with a count of zero for one allele in the RNA may arise either through strong allelic imbalance or through a genotyping error in which a homozygous site is incorrectly called heterozygous. To identify possible genotyping errors while preserving accurate read count information for downstream analyses, we developed a Bayesian hierarchical model called Genotyping Error Model (GEM). The model is applied to each heterozygous variant where one allele has zero reads and the gene has at least one heterozygous site with both alleles having non-zero counts.

The intuition behind this model is as follows. For a site $i$ having zero reads for one allele, all other exonic sites in the gene jointly provide evidence for the posterior distribution of allelic imbalance. That posterior then informs the likelihood of observing zero reads for one allele at site $i$, and that likelihood can in turn be used in a statistical test for whether the zero count might be due to a genotyping error instead of allelic imbalance. While this test can be done using Fisher's exact test or the beta-binomial test, those tests do not control Type I error for this task, because the null hypothesis for this task is conditioned on one allele count being zero while allowing for variation in the other allele count; in contrast, the foregoing tests instead condition on the total count at the site. Our GEM model rectifies this mismatch by conditioning on one allele having a count of zero. Our simulation results indicate that GEM controls Type I error under a wide range of scenarios, while the Fisher's exact test and beta-binomial tests do not (**Supplementary Table S24)**. Sites identified by GEM as likely genotyping errors instead of true het sites are eliminated from all of our downstream analyses.

GEM is a hierarchical model specified as follows:

$$p \sim uniform(0, 1)$$
$$\alpha \sim binomial(p, \beta)$$
$$X \sim binomial(p, n)$$
$$n \sim negative\ binomial(1 - (Var - mu)/Var, mu * mu/(Var - mu))$$

where $p$ is a latent variable representing allele frequency; $\alpha$ is the sum of minor haplotype counts from sites other than site $i$; $\beta$ is the sum of total counts from sites other than site $i$; $n$ is a latent variable representing the total count at a hypothetical site with a zero count for one allele; $X$ is the read count for one allele at the hypothetical site and is fixed at zero; $mu$ is the sample mean of total counts across all sites, and $Var$ is the sample variance in total counts across all sites. Using MCMC, we can draw samples from the posterior distribution for $n$ and use those to assess the probability of observing values at least as extreme as the observed $n$. That probability is taken as a p-value for the null hypothesis that the zero count for one allele at site $i$ is merely due to sampling error rather than a genotyping error; rejection of that null therefore indicates a likely genotyping error at the site.

GEM is implemented in JAGS [14], parameter definitions are in **Supplementary Table S24**. In running GEM, we generated 1000 warm-up samples and 7300 keeper samples. This number of MCMC samples was chosen to ensure that the binomial proportion was estimated with sufficient precision (95% confidence interval of the binomial proportion of 0.05 will be between 0.04 and 0.06–see **Supplementary Text 4.1**).

#### Supplementary Text 2.8 1000 Genome Project sample processing pipeline

We have developed a pipeline for rapid screening of NGS datasets for allele-specific expression (ASE) studies, this pipeline also addresses other confounding issues in ASE analysis, such as reference allele mapping bias and genotyping error, as described below. Our pipeline minimizes the incorrect assignment of alternative alleles to reference sequences using STAR2 RNA-seq alignment tool in EndtoEnd alignment mode coupled with WASP filtering (**Supplementary Text 4.2**). In addition, we utilized an unbiased RNA-seq fastq read simulator to eliminate genomic sites exhibiting allelic mapping bias, fostering more even read distributions across alleles (**Supplementary Text 2.6**). For genes having a zero count for one allele at a site genotyped as heterozygous, we introduce a Genotyping Error Model (GEM) to identify potential genotyping inaccuracies (**Supplementary Text 2.7**).

This pipeline requires three inputs: paired-end raw RNA-seq fastq data, chromosome-specific 1KGP VCF files, and information about the individual's ancestry. The pipeline proceeds as follows for each individual **(Supplementary Figure S18**):
(1) The 1KGP VCF files are combined into one file containing biallelic SNPs that have passed quality control and can be phased by SHAPEIT2.
(2) Raw RNA-seq fastq reads are trimmed and aligned to the hg19 genome using STAR v.2 with WASP filtering. We discard sites with mapping bias to STAR using unbiased splice reads simulation data. Then, we use Picard to mark duplicate reads.
(3) We test SNPs with one allele having zero reads for genotyping errors.
(4) Using a fitted model trained on NA12878, we predict the SHAPEIT2 switching error rate for each SNP pair.
(5) We estimate gene-level ASE using the quickBEAST model and calculate the ASE effect size (theta) using the binomial p-value output from quickBEAST.
(6) We keep only genes that have an average read depth at least 10 and are present in at least two individuals for each ancestry.

### Supplementary Text 3 Result

#### Supplementary Text 3.1 Type 1 Error and Power Assessment of quickBEAST

Controlling type I error is crucial in ASE detection, as it reduces the risk of false positives which can mislead interpretations, particularly in clinical research. To examine our model's proficiency in Type I error management, we compared it against baseline methods, with a focus on NaiveSum due to its superiority over another baseline, MajorSite.

Utilizing a parametrized simulator (**Supplementary Text 2.4**), we generated a total of 19,207,904 gene instances with 5% phasing error under the null hypothesis (θ=1) and alternative hypothesis (θ=0.5) based on binomial confidence interval calculation. This number is calculated because the actual number of genes with heterozygous variants in the real life data, GIAB sample, is less than 10,000. Our rigorous testing framework set the alpha level at 0.05 and aimed for a confidence interval width of 1e-06, necessitating an adjusted alpha (p) of 5e-06, calculated as 0.05/10,000. This led to the simulation of approximately 19 million null gene instances to ensure substantial statistical power, calculated via the formula:

$$\text{number of null genes: } n = \frac{p(1-p)}{(width/1.96)^2}$$

We modeled two coverage scenarios for each gene (difference in coverage affecting the qb output as in **Supplementary Figure S16**): a medium coverage (with 30 reads for each of the three heterozygous sites) and a high coverage (100 reads for each of the ten heterozygous sites). In the quickBEAST model, for each gene, 1000 null instances were simulated with reads assigned to both alleles according to a binomial distribution (p=0.5). We then applied quickBEAST to these null instances to establish a distribution of posterior estimates, fitting these to a skewed t-distribution. This process facilitated the calculation of p-values for the given posterior estimates from each gene. The empirical cumulative distribution function (CDF) plot (**Supplementary Figure S19**) demonstrates the effective fit to the skewed t-distribution, leveraging 19 million null data points.

To assess the control of Type I error in our model, we evaluated p-values generated by quickBEAST, NaiveSum, MajorSite, and Pseudo Phasing using a dataset of 19 million simulations (sample size calculations detailed in **Supplementary Text 4.2**). These simulations were conducted under the null hypothesis with a 5% phasing error and high coverage (10 heterozygous sites and 100 reads per site) with simulators detailed in **Supplementary Text 2.4**).

Our analysis shows that quickBEAST maintains strict control over Type I errors, with minimal false positives recorded, as presented in **Supplementary Table S25**. This finding underscores the rigorous false positive control of quickBEAST under our test conditions. Furthermore, quickBEAST demonstrated superior statistical power in detecting true ASE events compared to NaiveSum, especially in high coverage scenarios. For example, quickBEAST achieved a statistical power of 1.000, significantly outperforming the 0.921 power observed with NaiveSum. This indicates quickBEAST's enhanced capability in identifying true positive ASE events.

Pseudo Phasing also achieved perfect power (1.000) but at the cost of a higher Type I error rate (0.926), highlighting the trade-off between sensitivity and specificity in this method. In contrast, MajorSite exhibited lower power (0.903) while maintaining better control over Type I errors (0.035). These results emphasize the effectiveness of quickBEAST in high-coverage environments, where it not only maximizes power but also maintains stringent control over Type I errors, making it a robust tool for ASE detection.

To further evaluate the Type I error rates, we analyzed p-values from the 19-million simulation dataset under the null hypothesis at various significance levels (alpha). The observed and expected Type I errors, along with their absolute differences, are summarized in **Supplementary Table S26**. The analysis shows that the observed Type I error rates generally align with the expected rates, with slight deviations quantified by the absolute difference between the observed and expected values. For instance, at an alpha level of 0.05, the observed Type I error rate is 0.06058631, with an absolute difference of 0.01058631 from the expected rate of 0.05. As the alpha level decreases, the absolute differences similarly decrease, indicating a consistent pattern across the evaluated significance levels.

#### Supplementary Text 3.2 Comparative Analysis of ASE Detection Methods in NA12878

We evaluated QuickBEAST's performance in detecting ASE genes against the two binomial-based baseline methods (introduced in **section 3.2.1**), NaiveSum (NS) and MajorSite (MS), by using the NA12878 sample from the 1000 Genome Project. QuickBEAST identified 1,008 ASE genes, constituting 13.21% of the 7,628 genes examined, each featuring at least one exonic heterozygous site and non-zero coverage per site. This outcome surpasses the results of the baseline methods, with NS identifying 842 (11.04%) genes and MS identifying 577 (7.56%) ASE genes

When we compare the QuickBEAST's 13.21% ASE detection rate for the NA12878 sample to prior literature, ASEQ [15] and MBASED [16] reported detection rates of 6% and 4.3%, respectively. Our method leverages gold standard phasing from high precision source - Whole Genome Sequencing (WGS) data from the Genome In a Bottle consortium, analyzed a broader gene set (7,628 genes) compared to the 3,071 genes and 2,560 genes assessed by ASEQ and MBASED, which used same published lymphoblastoid cell line RNA-seq data from GEO and different phased genomic variants (germline WES data from the 1000 Genomes Project collection and the constructed personal diploid genome in AlleleSeq, respectively). Compared to them, QuickBEAST showed improved detection rates, though it did not match AlleleSeq's 19% detection rate (935 out of 4,829 genes) which employs a personalized genome construction and a refined binomial test. That study benefited from having parental data which facilitated the construction of highly accurate personal genomes. Here we are considering the more general situation where parental data is unavailable, making the accurate construction of personal genomes difficult.

An analysis across QuickBEAST, NS, and MS identified 498 genes commonly detected by all three methods (**Supplementary Figure S20**), demonstrating significant concordance with MS (93.76% of MS-detected genes) and NS (95.72% of NS-detected genes). However, QuickBEAST uniquely identifies 159 genes, more than two fold more than are uniquely identified by either NS or MS.

Further analysis reveals that ASE genes can be identified by QuickBEAST but not NS or MS, or the other way around, primarily due to the handling of genes with multiple heterozygous sites (hets). MS often overlooks these genes due to its preference for the highest coverage site, potentially missing subtle ASE signals present at lower coverage sites. Similarly, NS has challenges in dealing with the genes that have multiple hets. This is because NS tends to disregard phasing errors, which are more likely to happen in genes with multiple hets, due to its simple summation approach. In contrast, quickBEAST's model accounting for phasing error allows it to capture these subtle ASE signals, showing its high sensitivity in detecting ASE genes across wide genomic complexities.

#### Supplementary Text 3.3 Correlation between bi-allelic SNPs and ASE genes in Ancestral Cohorts

There was a strong positive correlation between the number of bi-allelic heterozygous exonic SNPs and the number of ASE genes in each ancestry group ($p<0.05$), consistent with prior research [17]. The Wilcoxon rank-sum test showed that the YRI group had a higher number of heterozygous SNPs, genes, and total genes with heterozygous SNPs than European groups. A positive Pearson correlation was also found between the number of heterozygous SNPs and the number of ASE genes for each of the five ancestry groups, with YRI exhibiting substantially higher numbers of heterozygous SNPs while maintaining comparable numbers of ASE genes (**Supplementary Figure S21A**). Additionally, sequencing read depth was positively correlated with the number of heterozygous SNPs, with YRI having higher numbers of heterozygous SNPs than other groups at comparable read depth (**Supplementary Figure S22**). All individuals had a comparable positive correlation (0.86) between sequencing read depth and the number of ASE genes regardless of ancestry (**Supplementary Figure S21CD**).

#### Supplementary Text 3.4 Ancestral-associated differences in ASE rates in Lymphoblastoid cell lines

Our finding reveals the significant variations in ASE rate influenced by genetic ancestry. Yoruba (YRI) individuals showed an average of 4,387 genes per individual that have averaged 10 reads per site, with an average ASE gene percentage of 7.64%, about 3.5 hets per gene, and a total of 15,474 heterozygous SNPs. Conversely, Utah (CEU) samples demonstrated slightly higher ASE rates at 7.68%, but with fewer genes on average (3,430) and a total of 10,945 het SNPs, alongside an average of 3.18 het SNPs per gene. The CEU population had a higher rate of genes with ASE than other groups (**Supplementary Figure S22C**), a difference that was statistically significant ($p = 0.001449$, one-sided Wilcoxon test). This significance also show in comparisons between the CEU group and each other ancestry groups ( Tuscan (TSI): $p = 0.002627$, Great Britain (GBR): $p = 0.02082$, Finland (FIN): $p = 0.02454$, YRI: $p= 0.004648$), despite comparable read depths in all ancestral groups (**Supplementary Figure S21B**). Meanwhile, individuals from other European groups (FIN, GBR,TSI) showed a narrower range in the number of testable genes (between 3,503-3,583), total het SNPs (11,294-11,658), hets per gene (3.21-3.24), and ASE rates (between 6.86%-7.03%).

### Supplementary Text 4 Calculation and code

#### Supplementary Text 4.1 Sample Size Determination

$$CI = p \pm z\sqrt{p}(1-p)/n$$

$$p\ is\ the\ sample\ proportion\ (p = 0.05,\ etc)$$

$$z\ is\ the\ zscore\ for\ 95\%\ CI\ (z = 1.96,\ etc)$$

$$n\ is\ the\ sample\ size$$

(1) $p = 0.05$, $z = 1.96$, marginal error (CI width) = 0.01 →
$0.01 = 2 \times 1.96\sqrt{(0.05 \times 0.95)}/n$, $n \cong 7,300$

(2) $Number\ of\ genes\ with\ hets\ we\ seen\ in\ GIAB\ sample < 10K;\ p = 0.05/10K = 5e-06,$
$z = 1.96$, marginal error (CI width) = 1e-06 →
$n\_NULL = p(1-p)/(width/1.96)\ **\ 2,\ n \cong 19,207,904$

#### Supplementary Text 4.2 STAR alignment tool code

```
STAR --twopassMode Basic --runThreadN 24 --genomeDir $star_ind \
    --readFilesIn $fastqDir/${sample}_FWD_paired.fq.gz $fastqDir/${sample}_REV_paired.fq.gz
\
    --alignEndsType EndToEnd \
    --waspOutputMode SAMtag \
    --varVCFfile $VCF \
    --outFilterMismatchNmax 10 \
    --outSAMtype BAM SortedByCoordinate \
    --outReadsUnmapped Fastx \
    --outSAMattributes NH HI NM MD AS nM jM jI XS vA vG vW \
    --readFilesCommand "gunzip -c" \
    --outFileNamePrefix $output_prefix
```

#### Supplementary Text 4.3 Logistic regression model and ANOVA R code

```
model <- glm(error ~ num_hets, data = grouped_df, family = binomial)
Call:
glm(formula = error ~ num_hets, family = binomial, data = grouped_df)

Deviance Residuals:
    Min       1Q   Median       3Q      Max
-2.5650  -0.4321  -0.4184  -0.4052   2.2542

Coefficients:
            Estimate Std. Error z value Pr(>|z|)
(Intercept) -2.52588    0.08868 -28.482  < 2e-16 ***
num_hets     0.06718    0.01764   3.809  0.00014 ***
```

```
---
Signif. codes:  0 ‘***’ 0.001 ‘**’ 0.01 ‘*’ 0.05 ‘.’ 0.1 ‘ ’ 1

(Dispersion parameter for binomial family taken to be 1)

    Null deviance: 1734.2  on 2864  degrees of freedom
Residual deviance: 1718.1  on 2863  degrees of freedom
AIC: 1722.1

Number of Fisher Scoring iterations: 5
```

anova_result <- anova(model, test="Chisq")

```
Analysis of Deviance Table

Model: binomial, link: logit

Response: error

Terms added sequentially (first to last)


         Df Deviance Resid. Df Resid. Dev  Pr(>Chi)
NULL                      2864     1734.2
num_hets  1   16.136      2863     1718.1 5.897e-05 ***
---
Signif. codes:  0 ‘***’ 0.001 ‘**’ 0.01 ‘*’ 0.05 ‘.’ 0.1 ‘ ’ 1
```

Bibliography


[1] BioStudies, “BioStudies < The European Bioinformatics Institute < EMBL-EBI.” Accessed: Aug. 03, 2024. [Online]. Available: https://www.ebi.ac.uk/arrayexpress/experiments/E-GEUV-1/samples/

[2] “[No title].” Accessed: Aug. 03, 2024. [Online]. Available: ftp://ftp-trace.ncbi.nlm.nih.gov/giab/ftp/release/NA12878_HG001/latest/GRCh37/

[3] “Gm12878-Illumina - SRA - NCBI.” Accessed: Aug. 03, 2024. [Online]. Available: https://www.ncbi.nlm.nih.gov/sra/SRX457730%5Baccn%5D

[4] “HapMap Project.” Accessed: Aug. 03, 2024. [Online]. Available: https://www.coriell.org/1/NIGMS/Collections/HapMap-project

[5] “Browse the CEPH Database.” Accessed: Aug. 03, 2024. [Online]. Available: https://www.coriell.org/1/NIGMS/Collections/CEPH-Resources

[6] S. Belsare *et al.*, “Evaluating the quality of the 1000 genomes project data,” *BMC Genomics*, vol. 20, no. 1, p. 620, Aug. 2019.

[7] “[No title].” Accessed: Jul. 31, 2024. [Online]. Available: ftp://ftp.1000genomes.ebi.ac.uk/vol1/ftp/pilot_data/release/2010_07/trio/snps/

[8] B. Carpenter *et al.*, “Stan: A Probabilistic Programming Language,” *J. Stat. Softw.*, vol. 76, Jan. 2017, doi: 10.18637/jss.v076.i01.

[9] C. P. Robert, V. Elvira, N. Tawn, and C. Wu, “Accelerating MCMC algorithms,” *Wiley Interdiscip. Rev. Comput. Stat.*, vol. 10, no. 5, p. e1435, Jun. 2018.

[10] 1000 Genomes Project Consortium *et al.*, “A global reference for human genetic variation,” *Nature*, vol. 526, no. 7571, pp. 68–74, Oct. 2015.

[11] J. D. Storey, *The False Discovery Rate: A Bayesian Interpretation and the Q-value*. 2001.

[12] O. Delaneau, J. Marchini, 1000 Genomes Project Consortium, and 1000 Genomes Project Consortium, “Integrating sequence and array data to create an improved 1000 Genomes Project haplotype reference panel,” *Nat. Commun.*, vol. 5, p. 3934, Jun. 2014.

[13] T. A. Myers, S. J. Chanock, and M. J. Machiela, “: An R Package for Rapidly Calculating Linkage Disequilibrium Statistics in Diverse Populations,” *Front. Genet.*, vol. 11, p. 157, Feb. 2020.

[14] “GitHub - michaelnowotny/pyjags: PyJAGS: The Python Interface to JAGS,” GitHub. Accessed: Jul. 31, 2024. [Online]. Available: https://github.com/michaelnowotny/pyjags

[15] A. Romanel, S. Lago, D. Prandi, A. Sboner, and F. Demichelis, “ASEQ: fast allele-specific studies from next-generation sequencing data,” *BMC Med. Genomics*, vol. 8, p. 9, Mar. 2015.

[16] O. Mayba *et al.*, “MBASED: allele-specific expression detection in cancer tissues and cell lines,” *Genome Biol.*, vol. 15, no. 8, p. 405, Aug. 2014.

[17] G. Luoni *et al.*, “Population-specific patterns of linkage disequilibrium in the human 5q31 region,” *Genes Immun.*, vol. 6, no. 8, pp. 723–727, Dec. 2005.
